# Machine learning engineered PoLixNano nanoparticles overcome delivery barriers for nebulized mRNA therapeutics

**DOI:** 10.1101/2024.11.03.621713

**Authors:** Dandan Zhang, Qin Xiao, Jie Tang, Kai Xiao, Tingting Chen, Yang Chen, Yuheng Liu, Lifeng Xu, Chao Li, Larry Cai, Shouyuan Huang, Ying Wang, Liusheng Peng, Joseph Rosenecker, Dan Shao, Quanming Zou, Shan Guan

## Abstract

There continues to be a dearth of competent inhalable mRNA delivery although it holds great potential for addressing a wide variety of refractory diseases. The huge advances seen with parenteral-administered lipid nanoparticle (LNP) have not been translated into nebulized mRNA delivery due to the aggressive nebulization process and insurmountable barriers inherent to respiratory mucosa. Here, we show amphiphilic block copolymers revealed by machine learning (ML) can spontaneously form stabilized nanoparticles (PoLixNano) with the lipids components of LNP and simultaneously impart the PoLixNano with “shield” (shear force-resistant) and “spear” (pulmonary barriers-penetrating abilities) capabilities. We present a ML approach that leverages physicochemical properties and inhaled mRNA transfection profiles of a chemically diverse library of polymeric components to validate the integration of “shield” and “spear” properties as highly predictive indicators of transfection efficiency. This quantitative structure-mRNA transfection prediction (QSMTP) model identifies top-performing amphiphilic-copolymers from more than 10000 candidates and suggests their mucus-penetrating ability outweights the shear force-resistant property in contributing to efficient mRNA transfection. The optimized PoLixNano substantially outperforms the LNP counterpart and mediates up to 1114-times higher levels of mRNA transfection in animal models with negligible toxicities. The PoLixNano promotes overwhelming SARS-CoV-2 antigen-specific sIgA antibody secretion and expansion of T_RM_ cells which collectively confers 100% protection in mice against lethal SARS-CoV-2 challenges and blocks the transmission of Omicron variant between hamsters. PoLixNano also displays versatile therapeutic potential in lung carcinoma and cystic fibrosis models. Our study provides new insights for designing delivery platforms of aerosol-inhaled mRNA therapeutics with clinical translation potential.

## Introduction

In vitro transcribed mRNA based technology have revolutionized the field of biomedicine due to its record-breaking speed of development as well as its flexible design to express any proteins that target various diseases^1,2^. The approved COVID-19 mRNA vaccines have been administered billions of times, saved millions of lives and became the Nobel Prize in 2023, all in just three short years^3^. Yet the field shows no signs of slowing down because mRNA technology has stimulated overwhelming interest in preventing and treating a wide variety of infectious, cancer-related, and genetic diseases that are refractory to current treatments^4–7^. Most notably, next-generation mRNA vaccines inoculated via airway route are urgently needed because it may not only provoke systemic immune responses that prevent severe disease but also elicits strong mucosal immunity—characterized by secretory IgA (sIgA) and lung tissue resident memory T cells (T_RM_)—capable of blocking viral transmission, a benefit beyond that of parenteral injected counterparts^8–12^. Additionally, inhaled delivery of mRNA therapeutics holds immense promise to treat lung cancers and diverse pulmonary diseases (e.g. cystic fibrosis, α-1 antitrypsin deficiency, and primary ciliary dyskinesia) in which causative treatments remain elusive even though the genetic origin has been clearly identified^13–16^.

Although the notion of inhaled delivery of mRNA is highly appealing, the expanding applications of mRNA therapeutics in the airway remain restricted by delivery challenges due to barriers posed by nebulization process as well as the sophisticated pulmonary system^17–19^. The nebulizer produces strong shear forces which destroy the integrity of nanoparticles encapsulating mRNA, leading to aggregation, disintegration, and mRNA leakage that completely abolish its biological functions^20–22^. Even if the mRNA nanoparticles were successfully aerosolized with intact structure, they still encounter physiological barriers inherent to the respiratory mucosa that expel the nanoparticles from reaching its target cells^23^. Among which, the most notorious and insurmountable barrier is the mucus layer that traps the nanoparticles and acts as a chemical and steric barrier to their diffusion^24,25^. While each of the aforementioned barrier presents a potential rate-limiting step, their relative impact on the efficacy of nebulized mRNA delivery remains poorly understood, hampering rational design of inhaled delivery systems.

The cutting-edge lipid nanoparticle (LNP) is the only clinically validated delivery system for mRNA therapeutics^26,27^. Albeit it has demonstrated substantial efficacy following parenteral route of administration, the LNP displays limited capability in addressing the aforementioned nebulization-related and lung-specific barriers. This limitation is evident by a latest CF gene therapy clinical trial initiated by TranslateBio, in which the aerosolized LNP formulations failed to show sufficient effects due to inadequate mRNA expression in bronchial epithelia^28^. Major efforts in improving this delivery modality have been focused on ameliorating the shear-force resistant properties of LNP via optimizing the type/ratio of lipid components and altering the nebulization buffer with addition of polymeric excipient^21,29–32^. Despite of an improvement in pulmonary mRNA transfection over traditional LNP^29,32^, these charge-repulsion or stability-enhancing excipient strategies fails in addressing the insurmountable airway related barriers (particularly the mucus layer), and their potency in clinical translation for mucosal vaccine, cancer treatment and gene therapy yet needs to be proved. Consequently, a groundbreaking design with considerations on the unique physiological structure and aggressive mucosal environment inherent to the respiratory tract, moving beyond the classic four-component lipid design paradigm originated from parenteral applications, is urgently needed to fully unlock the therapeutic potential of nebulized mRNA.

To address the above-mentioned challenges, we developed a rationally designed delivery platform termed PoLixNano (Poloxam-lipid mixed nanoparticles) by integrating machine learning (ML) discovered neutral amphiphilic block copolymers with lipid composition to yield compact nanoparticles displaying dual functional properties: stability and shear force-resistant during nebulization (“shield”) and enhanced mucus-penetrating and transepithelial transport capabilities (“spear”) (**Fig. 1a**). Compared to high throughput screening approaches which often involve redundant processes, high costs, and barely provide acurate structure-activity design rules, ML-driven design can personalize formulation development through more precise and efficient computational models^33,34^. To this end, we established an experimental training dataset via collecting the physicochemical features and transfection profiles of PoLixNano prepared by a library of polymeric components with diverse chemical structures. This dataset enabled the development of a proficient ML platform that revealed the “spear” penetrating property as more critical than the “shield” stability protection in maximizing transfection efficiency for inhaled mRNA nanoparticles. The ML-guided screening further identified critical structure features within the polymeric component of PoLixNano, highlighting amphiphilic block copolymers— both the well-known poloxamers being listed in Pharmacopoeia and their tetrafunctional analogs, poloxamines^12,35^—as the most effective candidates for achieving the necessary “shield” and “spear” functionalities. Whereas neither hydrophilic polymers nor hydrophobic polymers could fulfill the task.

**Fig. 1.**
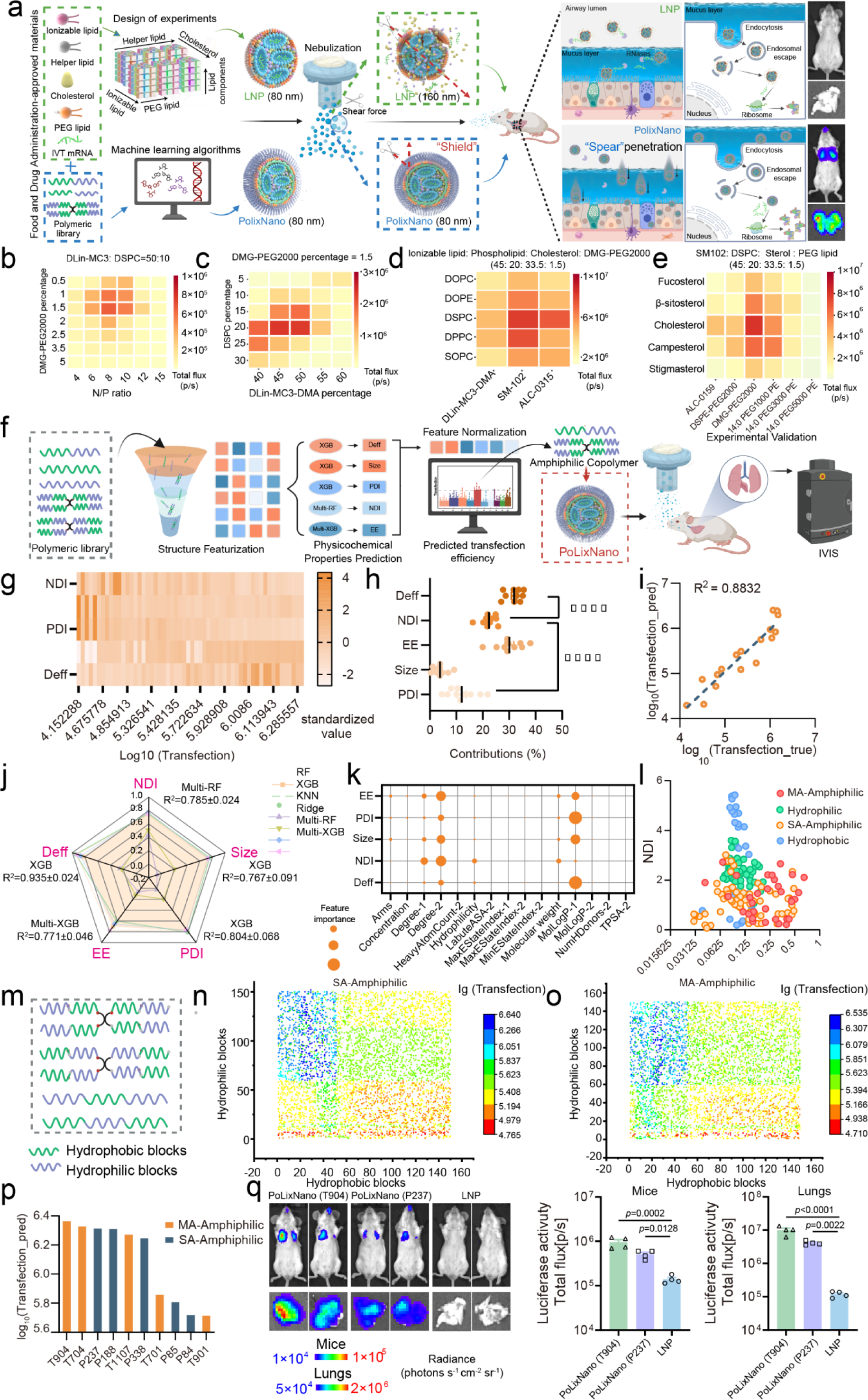
Optimization of PoLixNano formulations for nebulized mRNA delivery using machine learning algorithms. **a,** Schematic illustration of the preparation, optimization and nebulization of PoLixNano as well as LNP with a proposed pulmonary delivery process of the PoLixNano across airway mucus layer and cellular barriers for successful mRNA transfection. **b-e**, Heatmaps illustrating the transfection efficiency of PoLixNano encapsulating mRNA encoding firefly luciferase (Fluc-mRNA) with different lipid compositions, the concentration of T704 in (**b-e**) was fixed at 5 mg/mL, the ratio between nitrogen residues in ionizable lipid and nucleic acid phosphate groups (N/P ratio) in (**c-e**) was fixed at 8, the molar ratio of ionizable lipid: phospholipid: sterol-lipid: PEG-lipid in (**d**,**e**) was fixed at 45:20:33.5:1.5. The PoLixNano was administered to mice via nebulization, the bioluminescence signal in the lungs of mice was detected 6 h post-dosing (0.1 mg Fluc-mRNA per mouse). (n=3 biologically independent mice). The PoLixNano in (**b**) were prepared by various N/P ratios ranging from 4 to 15, along with 0.5 to 5 molar ratio of DMG-PEG2000. The molar ratio of DLin-MC3-DMA: DSPC was fixed at 50: 10. The PoLixNano in (**c**) were prepared by 5-30% (molar ratio) DSPC and 40-60% Dlin-MC3-DMA, with fixed 1.5% DMG-PEG2000. The PoLixNano in (**d**) were generated by incorporating various types of helper phospholipids (DSPC, DOPE, DPPC, DOPC, SOPC) and ionizable lipids (DLin-MC3-DMA, SM-102, ALC-0315). **e**, PoLixNano were further formulated with distinct sterol-components (fucosterol, β-sitosterol, cholesterol, campesterol and stigmasterol) and different types of PEG-lipids (ALC-0159, DSPE-PEG2000, DMG-PEG2000, 14:0 PEG1000 PE, 14:0 PEG3000 PE and 14:0 PEG5000 PE). Results in **b-e** suggested that the top performing PoLixNano candidate across the lipids library was composed of SM-102, DSPC, cholesterol, and DMG-PEG2000 (molar ratio = 45:20:33.5:1.5) with a N/P ratio of 8. **f**, Workflow of ML-guided PolixNano optimizing. It consists of four key parts: structure featurizing, multi-objective optimization, transfection efficiency prediction, and experimental verification. **g**, A heatmap illustrating the Log_10_(transfection efficiency) of PoLixNano with different physicochemical properties. **h**, Contributions normalized from regression coefficients of NDI (22.284 % ± 0.889 %), Size (3.828 % ± 0.828 %), PDI (12.058 % ± 1.722 %), EE (30.047 % ± 1.621 %), Deff (31.783 % ± 0.971 %) on ten-folds cross-validation. **i**, The scatter contains the true Log_10_(transfection efficiency) value and the predicted Log_10_(transfection efficiency) value made by the ensemble for each PoLixNano in the test set. **j**, Perfomance of different ML algorithms for Size, PDI, EE, NDI and Deff prediction on ten-folds cross-validation. **k**, Mean feature importance from ML of each task (Size: XGB, PDI: XGB, EE: Multi-XGB, NDI: Multi-RF, Deff: XGB) on five-folds cross-validation. Names of features are derived from rdkit.Chem.Descriptors._desclist or manually labeled polymer structure descriptors (X-1: descriptor X for hydrophobic monomer, X-2: descriptor X for hydrophilic monomer). **l**, Deff and NDI distributions of PolixNano prepared by hydrophilic, hydrophobic, single-arm amphiphilic (SA-Amphiphilic) and multi-arm amphiphilic (MA-Amphiphilic) polymers. **m**, Schematic illustration of structure space for four-arms amphiphilic and single-arm amphiphilic copolymer optimizing, including length, position and proportion of hydrophilic and hydrophobic blocks in polymers. **n**-**p**, Predicted Log10(transfection efficiency) values of PoLixNano prepared by SA-Amphiphilic (**n**), MA-Amphiphilic (**o**) and top 10 best-performing commercially available amphiphilic (**p**) copolymers. **q**, Representative bioluminescence images (left) and quantitative comparison (right) of Fluc-mRNA expression in live mice and excised lungs 6 h post-nebulization mediated by the top-perforiming T904-based PoLixNano, poloxamer 237 (P237)-based PoLixNano or LNP counterpart (n=4 biologically independent animals). A one-way analysis of variance (ANOVA) with Tukey’s post-hoc test was applied to determine significance in **q**.

Upon nebulization under optimized conditions, PoLixNano demonstrated excellent stability and structural integrity without size changes and notable mRNA leakage, while the size of LNP counterpart doubled with almost one third encapsulated mRNA leaked out. Improved cellular uptake and endosome escape were observed in PoLixNano, resulting in levels of target-protein expression from PoLixNano increased up to 401 times higher in cultured cells and 30 times enhancement in organoids compared to the LNPs. By virtue of its promising mucus-penetrating and transepithelial transporting ability, nebulized PoLixNano is able to induce 35-to 1114-times higher levels of mRNA transfection than the LNP counterpart in the airway of a range of animal models, and substantially outperforms the recently described LNP-HP08^Loop^ strategy^32^. The PoLixNano primarily targets pulmonary dendritic cells and bronchial epithelium with negligible toxicities, making it an ideal delivery platform for prophylactic mucosal vaccines and gene therapy. Indeed, PoLixNano encapsulating mRNA encoding the receptor binding domain (RBD) of SARS-CoV-2 induced durable and comprehensive adaptive immune responses, including mucosal immunity (sIgA antibody in the airways), systemic immunity (neutralizing serum IgG antibody), cellular immunity (lung-T_RM_ cells), and trained innate immune responses. All these merits conferred virtually 100% protection of susceptible mice against lethal SARS-CoV-2 challenges and significant benefits against host–host transmission of the Omicron variant between hamsters than the parenteral vaccination regimen, offering robust protection against breakthrough infection. Additionally, PoLixNano exhibited therapeutic potential in a metastatic lung cancer model and enabled abundant expression of mRNA encoding the cystic fibrosis transmembrane conductance regulator (CFTR) in the bronchial epithelia of *CFTR*-deficient mice, underscoring its versatility for monogenic disorder gene therapy. Considering all the components of PoLixNano could switch to US Food and Drug Administration-approved materials, this platform offers a translatable solution for pulmonary mRNA therapeutics across multiple disease applications.

## Results

### ML-guided design of PoLixNano with shear force-resistant and mucus-penetrating properties

The PoLixNano could be prepared by a simple self-assembly of lipids, in vitro transcribed messenger RNA (mRNA) and a polymeric component via microfludic mixing. Based on our previous experiences, we selected poloxamine 704 (T704) as a prototype copolymer since it has proven to be extremely promising in mediating efficient pulmonary gene transfection within small and large animals than were achieved with the state-of-the-art liposome-based formulations^12,14,36^. Starting from a classic lipid composition used in approved COVID-19 mRNA vaccines together with optimized concentration of T704 (**Supplementary Fig. 1a**), we employed the orthogonal design of experiments (DOE) to create a library of PoLixNano formulations aimed at modulating lipid compositions and varying their molar ratio in order to find out the top-performing candidate (**Fig. 1b-1e**).

After optimizing the lipid compositions of PoLixNano, we planned to screen potent polymeric compounents with diverse chemical sturctures, ranging from hydrophilic, hydrophobic, amphiphilic, positively-charged, negatively-charged and neutral polymers. Considering their structural space is vast, and alterations in their concentrations can drastically impact the biological functions of PoLixNano, the subsequent screening of these polymeric compounents remains a labour-intensive and time-consuming task, with limited chance to reveal underlying mechnisms in terms of structure-activity relationships from those sophisticated chemical and formulation parameters. Given machine learning (ML) techniques have demonstrated considerable promise in drug discovery and novel lipids design^33,34^, we aimed at exploiting ML to facilitate the discovery of polymeric candidates within a formulation library (**Supplementary Table 2**) that comprises 264 PoLixNano prepared by 66 polymers with four distinct concentrations (**Fig. 1f**). The physicochemical properties of each PoLixNano both pre- and post-nebulization were subsequently evaluated, including particle size (Size), polydispersity index (PDI) and mRNA encapsulation efficiency (EE). Importantly, nebulization shear force deformation index (NDI), and diffusion coefficient (Deff) respectively representing the ability of shear force-resistant and mucus-penetrating, were collected as two key paraments of PoLixNano. The Pearson correlation coefficents (r) confirmed that each of these five parameters was significantly correlated with in vivo transfection profiles (**Fig. 1g** and **Extended Data Fig. 1a-e**). We constructed a linear regression model to assess the contribution of each feature towards transfection efficiency using ten-folds cross-validation. Interestingly, Deff accounted for approximately 31.78% of the variance in transfection efficiency, surpassing the 22.28% contribution from NDI (**Fig. 1h**). Next, the transfection dataset was randomly splitted into training set and test set at a ratio of 7:3. The fitted equation (Eqs. 1) performed with R^2^ value of 0.8832 on test set (**Fig. 1i**), suggesting this model might effectively capture the relationship between those five features and transfection efficiency.

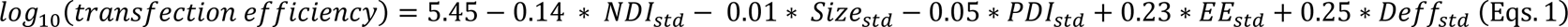

Encouraged by these findings, we sought to develop quantitative structure-mRNA transfection prediction (QSMTP) model to discover best-performing polymers by the structural data. We assessed the learning capability of different ML algorithms, and five-folds cross-validation were used to aviod the accidental bias caused by dataset splitting. eXtreme Gradient Boosting (XGB) yielded the best predictions on the Size, PDI and Deff, while Multi-task random forest regressor (Multi-RF) and Multi-task XGB regression (Multi-XGB) offered better performance on the prediction of NDI and EE, respectively (**Fig. 1j** and **Extended Data Fig. 1f-j**). Next, we deciphered the mechanisms underlying the relationship between structural properties and physicochemical characteristics by calculating feature importance (**Fig. 1k**). It reveals that “MolLogP” (representing hydrophobic properties) and “degree” (representing quantity of hydrophilic and hydrophobic monomers) of a certain polymer contributed largely to both shear force-resistant and mucus-penetrating ability of PoLixNano. These findings indicate that the composition and pattern of hydrophilic and hydrophobic blocks in the polymer might be the key factors that influencing the delivery properties of PoLixNano in the airway^37^.

Further analysis indicates that PoLixNano prepared from amphiphilic polymers, particularly multi-arm amphiphilic polymers (MA-Amphiphilic), simultaneously exhibited superior “shield” and “spear” characteristics (**Fig. 1l** and **Extended Data Fig. 1k-o**). Using such poloxamine and poloxamer frameworks as templates, the ensembled algorithm of ML and linear regression was applied for in silico optimizing the best-performing polymer (**Fig. 1m**). From more than 10000 potential canditates, we identified the top-performing commercially available copolymers (T904) (**Fig. 1n**), and found T904-based PoLixNano significantly outperforms the nebulized LNP (LNP-neb) counterpart (**Fig. 1o**) as well as recently reported iLNP-HP08^Loop^ featuring a high helper lipid ratio, acidic dialysis buffer, and excipient-assisted nebulization buffer (**Extended Data Fig. 2a-e**). Although being relatively less efficient, we found PoLixNano prepared with one lipid component removed still displayed robust mRNA transfection in the respiratory tract of mice whereas the three-components LNP counterparts barely mediated any observable transfection signal (**Extended Data Fig. 2f**), indicating the necessity of integrating polymeric components. Collectively, we selected T904-based PoLixNano as the best-performing candidate for all following investigations.

### Characterizations of the best-performing PoLixNano with shear force-resistant and mucus-penetrating properties

We next characterized the shear force-resistant properties of the best-performing PoLixNano. Surprisingly, the particle size of nebulized PoLixNano (PoLixNano-neb) remained at around 87.2 nm (no size change) with a low polydispersity index (PDI) of 0.06 while the aerosolization process drastically increased the size of LNP from 73.4 nm to 154.6 nm (more than 110.5% size changes) with a large PDI of 0.23 (**Fig. 2a**). Above 84.6% of mRNA remains encapsulated within PoLixNano-neb (less than 8.8% loss), in contrast to a less than 62% of mRNA encapsulation efficiency of LNP-neb (more than 36% leakage) (**Fig. 2b**). Besides, PoLixNano maintained negative zeta potential and consistent viscosity both pre- and post-nebulization (**Extended Data Fig. 3a**,3b), and remained stable even being stored at room temperature for 14 days (**Extended Data Fig. 3c-e**). Agarose gel electrophoresis assay confirms similar results and suggests PoLixNano maintained superior structural integrity post-nebulization over LNP counterpart which showed distinct mRNA leakage (**Extended Data Fig. 3f**). In mRNA release analysis, PoLixNano required at least 15 mM sodium dodecyl sulfonate (SDS) for dissociation, compared to 8 mM SDS for LNP (**Extended Data Fig. 3g**). Transmission electron microscopy (TEM) directly unveiled a compact spherical morphology with monodisperse status of non-nebulized and nebulized PoLixNano, as well as the non-nebulized LNP (**Fig. 2c**). However, LNP appeared to aggregate post-nebulization and displayed irregular, clustered and amorphous structure with uneven sizes (**Fig. 2c**).

**Fig. 2.**
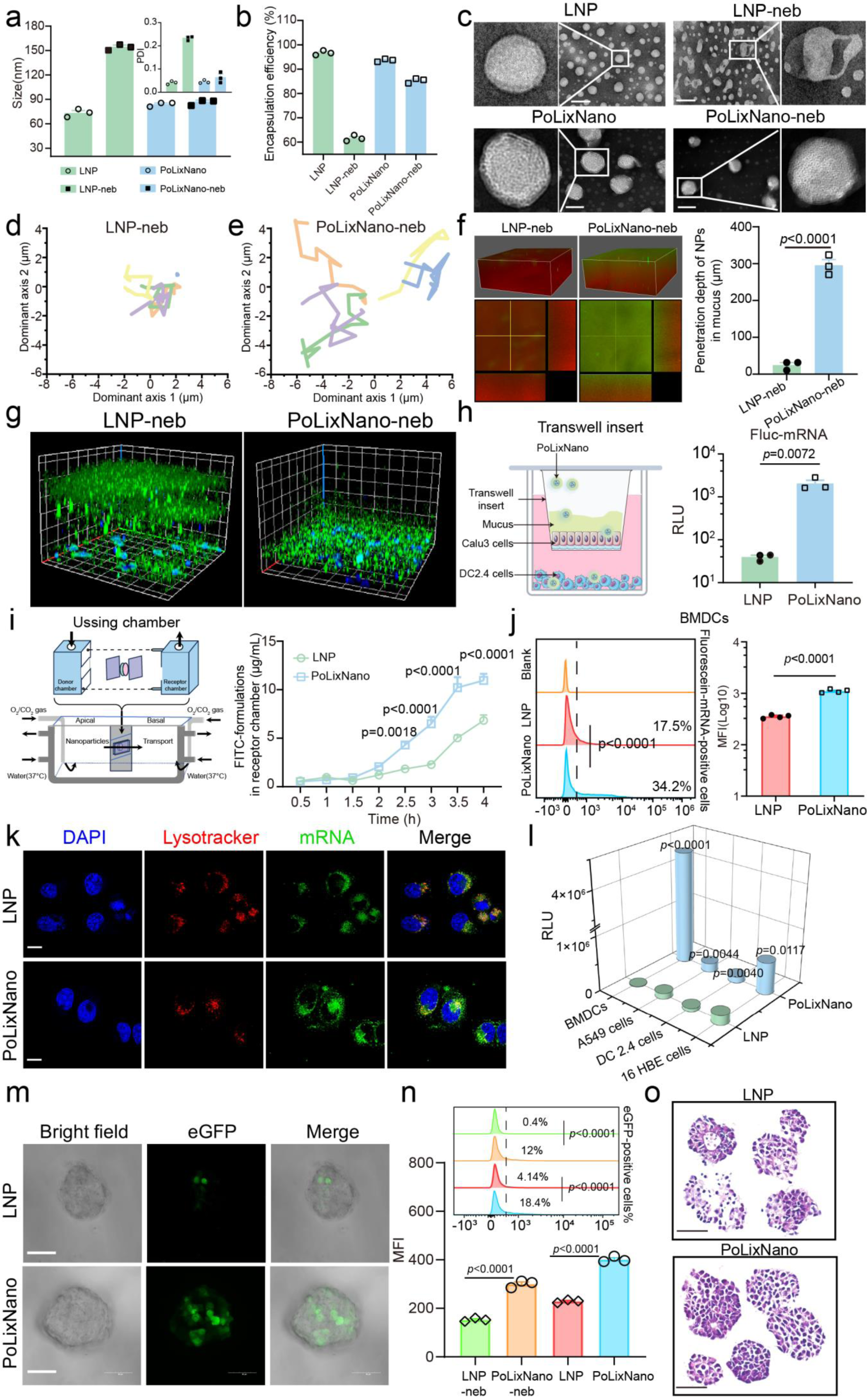
Characterization and in vitro evaluations of the best-performing PoLixNano. **a-c,** Physicochemical characterization of PoLixNano and LNP counterpart before and after nebulization. **a,** The dynamic light scattering (DLS) measurements for the particle size and polydispersity index (PDI) of PoLixNano or LNP pre- and post-nebulization. **b**, The mRNA encapsulation efficiency (EE%) of PoLixNano and LNP before and after nebulization. **c**, Transmission electron microscopy (TEM) micrograph of non-nebulized and nebulized PoLixNano (scale bar: 100 nm) or LNP (scale bar: 200 nm). **d**,**e**, Representative trajectories of LNP (**d**) or PoLixNano (**e**) nanoparticles after nebulization in mucus mimicking gel detected by a multiple particle tracking (MPT) assay. Each color represents the trajectory of one independent particle. **f**, 3D fluorescence images and sectional fluorescence images depicting the penetrating profiles of nebulized PoLixNano (PoLixNano-neb) or LNP (LNP-neb) in the mucus-simulating hydrogel, along with the corresponding quantitative penetration depths comparison. **g**, 3D confocal microscopy of the mucus penentrating profiles of PoLixNano-neb or LNP-neb in Calu-3 cells with secreted thick mucus layer. Green channel, fluorescence-labeled nanoparticles; blue channel, nuclei stained by DAPI. **h**, Schematic showing the transportation and successful expression of Fluc-mRNA encapsulated in PoLixNano or LNP across a monolayer formed by Calu-3 cells within a Trans-well insert assay system (left). The luciferase expression within the basolateral seeded DC2.4 cells was evaluated 24 h after the indicated samples were added to the apical compartment (right) (n=3 independent experiments). **i**, Schematic diagram of the Ussing Chamber System to assess transepithelial penetration profiles of PoLixNano or LNP (left). Concentrations of fluorescence-labeled nanoparticles in the receptor chamber over time was measured (right) (n=3 independent experiments). **j**, Cellular uptake of Fluorescein-mRNA encapsulated in PoLixNano or LNP counterpart in bone marrow-derived dendritic cells (BMDCs) measured by flow cytometry (n=4 biologically independent samples). Bar chart represents the mean fluorescence intensity (MFI) per cell, and the histogram displays the representative ratio of Fluorescein-mRNA positive cells. Untreated BMDCs were used as the blank control. **k**, Subcellular location of the Fluorescein-mRNA@PoLixNano and Fluorescein-mRNA@LNP after 4 h incubation with 16HBE cells. Green channel, Fluorescein-mRNA; blue channel, nuclei stained by DAPI; red channel, endocytic vesicles marked by LysoTracker™ Red DND-99; Merge, combination of the aforementioned channels. Scale bars: 15 µm. **l**, Transfection efficiency of MetLuc-mRNA@PoLixNano in various cell lines. The luciferase activity was represented as relative light units (RLU) (n=3 biologically independent samples). **m**, Representative expression of eGFP-mRNA in organoids after 36 h incubation of PoLixNano or LNP based formulations. Images from left to right depict bright-field microscopy images of individual organoid, eGFP signal (green channel), and an overlay of the two images. Scale bars: 50 µm. **n**, Quantification of eGFP-positive cells (histogram) and MFI (bar chart) in organoids transfected by non-nebulized LNP or PoLixNano, and post-nebulization counterparts (LNP-neb or PoLixNano-neb) (n=3 biologically independent samples). **o**, Hematoxylin and eosin (H&E) staining of organoids treated with PoLixNano or LNP. Scale bar: 50 μm. The results of **c-g**, **k**, **m** and **o** are representative images from at least three independent experiments with similar results. Statistical significance was analyzed using a one-way ANOVA with Tukey’s post-hoc test in **n**, two-tailed unpaired *t*-tests in **f**, **h**, **j** and **l**, and two-way ANOVA with Sidak’s multiple comparisons test in **i**.

To reveal the penetration capacity of PoLixNano, a multiple particle tracking (MPT) assay was performed. The trajectories of the PoLixNano and LNP based nanoparticle motions were recorded, with representative trajectories shown in **Fig. 2d,2e** and **Extended Data Fig. 3i**. The distinct movement patterns suggest PoLixNano hold the capability of efficient mucus penetrating. In contrast, LNP was almost trapped in a small area by the mucus network. This trend could be further proved by the mean square displacement (MSD) and the effective diffusivities (Deff), both of which were substantially enhanced in PoLixNano group compared to the LNP counterpart (**Extended Data Fig. 3h**,3i). We further assessed PoLixNano in a fluorescence labeled mucus-simulating hydrogel via the confocal microscopy. The penetration depth of PoLixNano was 12-fold higher post-nebulization (**Fig. 2f**) and 2.9-fold higher pre-nebulization **(Extended Data Fig. 3j**) compared to the LNP counterpart. Additionally, a large number of LNP was trapped and localized far from a mucus secreting cell layer, while both pre- and post-nebulized PoLixNano substantially penetrated deeper into the mucus layer and successfully reached the cells (**Fig. 2g** and **Extended Data Fig. 3k**). Improved transmucosal penetration and transepithelial transportation of PoLixNano could further be reflected by the significantly enhanced mRNA transfection within cells seeded at the basolateral side of a Trans-well insert system compared to LNP counterpart (**Fig. 2h**). We also found the PoLixNano diffused much faster across excised bronchial tissues of rats than the LNP did using an Ussing Chamber System (**Fig. 2i**). These findings collectively highlight the essential roles of the amphiphilic block copolymer within the PoLixNano in facilitating efficient mucus diffusivity and transepithelial transportation.

### PoLixNano facilitates efficient mRNA tranfection in cultured cells and organoids with negligible cytotoxicity

As an efficient cellular uptake is the prerequisite of successful mRNA transfection, we quantified the internalization of different formulations within cultured cells using fluorescence labeled mRNA (Fluorescein-mRNA). Flow cytometry analysis revealed that the PoLixNano group exhibited significantly higher uptake efficiency compared to the LNP in all tested cell lines (**Fig. 2j and Extended Data Fig. 4a**). To further elucidate underlying mechanism, cell entry pathways were investigated. As shown in **Extended Data Fig. 4b**, PoLixNano could utilize efficient energy-dependent cellular uptake mediated by electrostatic absorptive endocytosis, micropinocytosis and clathrin-mediated endocytosis. Additionally, confocal microscopy indicated abundant Fluorescein-mRNA locolized in the cytosol when delivered by PoLixNano, while the LNP encapsulated counterpart was limited and confined within lysosome, implying the PoLixNano efficiently enables Fluorescein-mRNA to facilitate endosome escape (**Fig. 2k**).

In order to assess the overall transfection efficiency of PoLixNano in various cell lines, mRNA encoding Metridia luciferase (MetLuc-mRNA) was loaded into PoLixNano (MetLuc-mRNA@PoLixNano) and LNP (MetLuc-mRNA@LNP). MetLuc-mRNA@PoLixNano mediated a 401-fold increased transfection in bone marrow derived dendritic cells (BMDCs) and significantly higher levels of luciferase expression in other tested cell lines compared to MetLuc-mRNA@LNP (**Fig. 2l**). A similar trend was observed when using mRNA encoding enhanced green fluorescent protein (eGFP-mRNA) (**Extended Data Fig. 4c**,4d). A viability assay revealed that 93.06% of 16HBE cells and 77.13% of DC2.4 cells survived after PoLixNano transfection, while only 48.31% and 62.87% of which remained viable in the LNP counterpart (**Supplementary Fig. 1b**). Besides, we investigated the ability of different formulations in promoting dendritic cell (DC) maturation. BMDCs treated by PoLixNano exhibited higher expression of activation markers, including CD40, CD80, CD86 and MHCII, compared to LNP treated BMDCs (**Extended Data Fig. 4e**).

We also employed an organoid model with three-dimensional (3D) structures, which are extremly difficult to transfect because it recapitulate key characteristics of the in vivo status (such as differentiation and tight junctions)^38,39^, to further evaluate the transfection profiles. Both confocal microscopy and flow cytometry revealed substantially improved ratios of eGFP positive cells in organoids incubated with eGFP-mRNA@PoLixNano (18.4%) compared to eGFP-mRNA@LNP (4.1%) (**Fig. 2m,2n** and **Supplementary Fig. 1c**,1d). Even though the eGFP signal displayed by nebulized PoLixNano (PoLixNano-neb, 12.0%) was relatively lower than non-nebulized one, it remained significantly higher than the post-nebulized LNP counterpart (LNP-neb, 0.38%) (**Extended Data Fig. 4f** and **Supplementary Fig. 1c**,1d). A similar trend was observed when using Fluc-mRNA, in which LNP mediated transfection immediately deminished to the backgroud level once nebulization was involved (**Extended Data Fig. 4g**). Histopathological analysis revealed that organoids treated by PoLixNano and LNP were similar to the PBS control (**Fig. 2o and Extended Data Fig. 4h**). These findings collectively indicated that PoLixNano hold the shear force resistant properties and offers superior transfection efficiency in in vitro models compared to LNP counterpart.

### Aerosolized PoLixNano mediates safe and significantly superior mRNA transfection in animal models

For in vivo evaluations, we first investigated the bio-distribution of nebulized PoLixNano in mice, together with the LNP control. The fluorescence labelled formulations primarily localized in the lung, with PoLixNano mediated significantly higher levels of fluorescence accumulation than LNP counterpart (**Fig. 3a**). Flow cytometry revealed superior cellular uptake of PoLixNano compared to LNP among various airway related cells (**Fig. 3b**). Efficient internalization of PoLixNano into epithelial cells (EpCAM) and antigen-presenting cells (CD11C) was further validated by immunofluorescent staining (**Fig. 3c**).

**Fig. 3.**
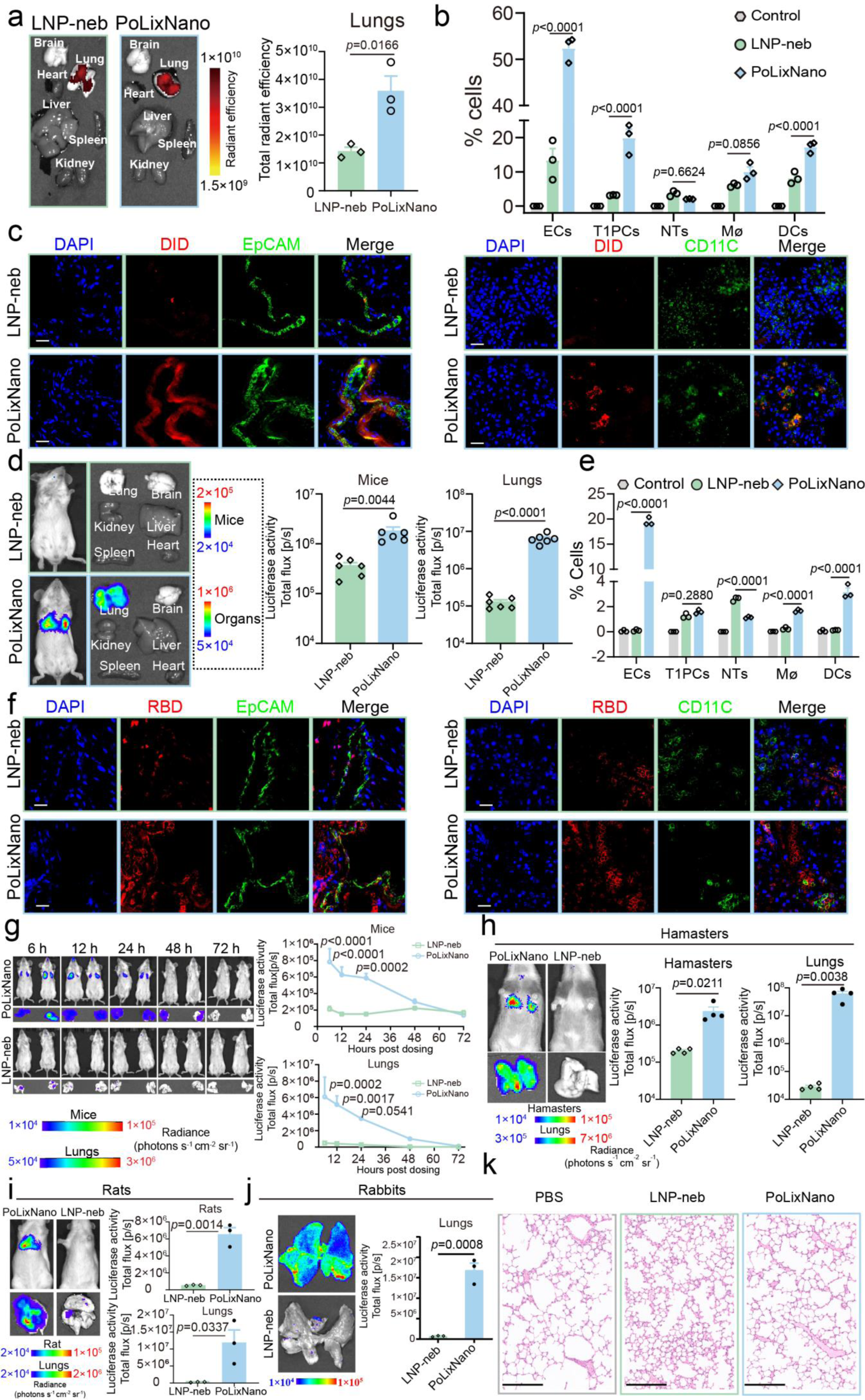
Bio-distribution and transfection profiles of nebulized PoLixNano in animal models. **a**, Distribution of fluorescence labeled formulations in excised organs of mice 6 h post-nebulization with PoLixNano or LNP (left panel), with a quantitative comparison of their signal intensities (right panel) (n=3 biologically independent samples). **b**, Quantitative analysis of fluorescence labeled PoLixNano or LNP in different types of pulmonary cells collected from the lung of treated mice 6 h post-nebulization (n=3 biologically independent mice). ECs: Epithelial cells; T1PCs: Type 1 pneumocytes; NTs: Neutrophils; Mø: Macrophages; DCs: Dendritic cells. Samples from untreated mice were served as the Control. **c**, Representative immunostaining images showing the distribution of fluorescence labeled PoLixNano or LNP in lung sections 6 h post-nebulization. For each panel, the images from left to right show nuclei (DAPI, blue), formulations (DiD, red), EpCAM (epithelial cell adhesion molecule which represents the epithelial cells, green in the left panel) or CD11C (a marker of dendritic cells, green in the right panel), and an overlay of the three images (Merge). Scar bar: 20 μm. **d**, Bioluminescence imaging of Fluc-mRNA expression (6 h post-nebulization) mediated by nebulized PoLixNano or LNP in mice and excised organs (left), with corresponding quantitative signal intensities (right) (n=7 biologically independent animals). **e**, Quantitative analysis of different types of pulmonary cells successfully transfected by eGFP-mRNA within lung samples collected 6 h post-transfection of nebulized LNP or PoLixNano, including ECs, T1PCs, NTs, Mø and DCs (n=3 biologically independent mice). Samples from untreated mice were served as the Control. **f**, Immunohistochemical analysis of the receptor binding domain (RBD) protein of SARS-CoV-2 (red channel) encoded by the successfully transfected RBD-mRNA in the lungs of mice mediated by the nebulized LNP or PoLixNano, the green channel represents EpCAM or CD11C, the bule channel shows nuclei stained by DAPI. Scar bar: 20 μm. **g**, Expression kinetics of Fluc-mRNA encapsulated in nebulized PoLixNano or LNP over time in mice and excised lungs (n=4 biologically independent animals). **h-j**, Bioluminescence imaging and quantification of Fluc-mRNA expression in hamsters (**h**), rats (**i**), and rabbits (**j**) 6 h post-nebulization with PoLixNano or LNP (n=3 biologically independent animals). **k**, Hematoxylin and eosin (H&E) staining of lung sections harvested from mice treated with nebulized PoLixNano or LNP 24 h post-nebulization (scale bar: 200 μm). The images of **a**, **c**, **d** and **f**-**k** are representative results from at least three independent experiments with similar results. Significant differences are assessed using two-tailed unpaired *t*-tests in **a**, **d**, **h**, **i** and **j**; Two-way ANOVA with Sidak’s multiple comparisons test in **g**; Two-way ANOVA withTukey’s multiple comparisons test in **b** and **e**.

We then examined the in vivo transfection using Fluc-mRNA. Bioluminescence signals from live mice and excised organs demonstrate that PoLixNano only transfected the lungs of mice and significantly outperformed the LNP counterpart (**Fig. 3d**). Quantification of eGFP-mRNA expression among various types of pulmonary cells revealed that eGFP-mRNA@PoLixNano successfully transfected substancially more airway epitheliums (19.5%) and pulmonary DCs (3.17%) compared to eGFP-mRNA@LNP (0.10% and 0.12%). PoLixNano also slightly transfected other cell types including macrophages, type I pneumocytes and on rare occasions neutrophils (**Fig. 3e**). Immunofluorescent analysis of lung sections further confirms that PoLixNano induced prosperous mRNA transfection signal in bronchial epithelial cells and antigen-presenting cells with a widespread distribution pattern in contrast to LNP counterparts (**Fig. 3f**).

We further investigated the kinetics of mRNA expression mediated by PoLixNano and LNP. In contrast to the background level signal in LNP treated mice since 6 h post-dosing, overwhelmingly stronger bioluminescence signals both in live mice and excised lungs could be consistantly observed in PoLixNano counterparts up to 72 h post-dosing (**Fig. 3g**). The mRNA expression mediated by PoLixNano also exhibited dose-dependency (**Extended Data Fig. 5a**). The delivery potential of PoLixNano were further evaluated in other animal models, in which PoLixNano facilitated more than 1114.5-fold, 52.6-fold, 35.2-fold higher mRNA transfection in the lungs of hamasters (**Fig 3h**), rats (**Fig 3i**), and rabbits (**Fig 3j**), respectively, compared to the LNP counterpart.

To investigate potential toxicity of PoLixNano, we performed a histopathological analysis and found no sign of inflammation or toxicity neither in lung sections (**Fig. 3k**) nor in other major organs of mice (**Extended Data Fig. 5b**). The expression levels of four key pro-inflammatory cytokines (i.e. interleukin-6 (IL-6), tumor necrosis factor-α (TNF-α), interleukin-4 (IL-4) and interleukin-17 (IL-17)) in lung homogenates, serum and bronchoalveolar lavage fluid (BALF) samples also showed no significant differences compared to the control (**Extended Data Fig. 5c and Supplementary Fig. 2a**). The complete blood chemistry metrics suggest that all parameters remained within the normal range over 21 days post-dosing (**Extended Data Fig. 5d and Supplementary Fig. 2b**). We also confirmed no detectable anti-PoLixNano antibodies were found in serum within 21 days (**Extended Data Fig. 5e**), which further proved the safety profile of PoLixNano.

### PoLixNano provides comprehensive and long-lasting immunity against SARS-CoV-2 antigen

To explore the potential of PoLixNano in mediating mucosal immune responses, the mRNA encoding RBD (RBD-mRNA) was loaded into PoLixNano (RBD-mRNA@PoLixNano) and immunized BALB/c mice via a prime-boost strategy as illustrated in **Fig. 4a**. We observed RBD-mRNA@PoLixNano triggered the highest levels of RBD-specific IgG antibody in serum samples which is even higher than intramuscularly injected LNP counterpart (RBD-mRNA@LNP-i.m), whereas RBD-specific antibodies were almost undetectable in nebulized LNP counterpart (RBD-mRNA@LNP-neb) (**Fig. 4b**). Notably, PoLixNano maintained persistently and relatively high levels of anti-RBD specific IgG antibody over 380 days (**Extended Data Fig. 6a**) probably due to the significantly elevated amounts of germinal center B cells (GCB cells, CD19+B220+CD95+Gr1+) and T follicular helper cells (Tfh cells, CD4+CXCR5+PD1+) which coordinated to facilitate B cell maturation^40^ (**Fig. 4c** and **Extended Data Fig. 6b**). The anti-RBD secretory immunoglobulin A (sIgA) antibody titer detected in BALF samples on day42 reached more than 1/3200 **(Extended Data Fig. 6c**), which are even higher than those induced by intranasally vaccinated counterparts using adenovirus-vector (∼1/100) or adjuvanted subunit vaccines (∼1/1000) according to previous publications^41,42^. Consistently, RBD-mRNA@PoLixNano vaccinated mice showed significantly higher levels of RBD-specific sIgA in BALF and nasal lavage fluid (NLF) samples even on day380 compared to other groups displaying a background level signal (**Fig. 4d**). Pseudovirus neutralization assays suggest the neutralizing antibody titer at 50% inhibition (pVNT_50_) in PoLixNano vaccinated mice reached approximately ∼1/71450 in serum and ∼1/422 in BALF samples (**Fig. 4e** and **Extended Data Fig. 6d**), which even outperforms LNP-i.m. counterpart. We additionally found RBD-mRNA@PoLixNano triggered the highest expansion of RBD-specific IgG antibody secreting plasma cells (ASCs) in bone marrow cells (BMCs), mediastinal lymph node cells (MLNs), and splenocytes, concurrent with high levels of RBD-specific IgA ASCs being detected in BMCs and MLN (**Fig. 4f** and **Extended Data Fig. 6e**). Altogether, the high amounts of PoLixNano induced ASCs may attribute to the persistent systemic and mucosal responses^43^.

**Fig. 4.**
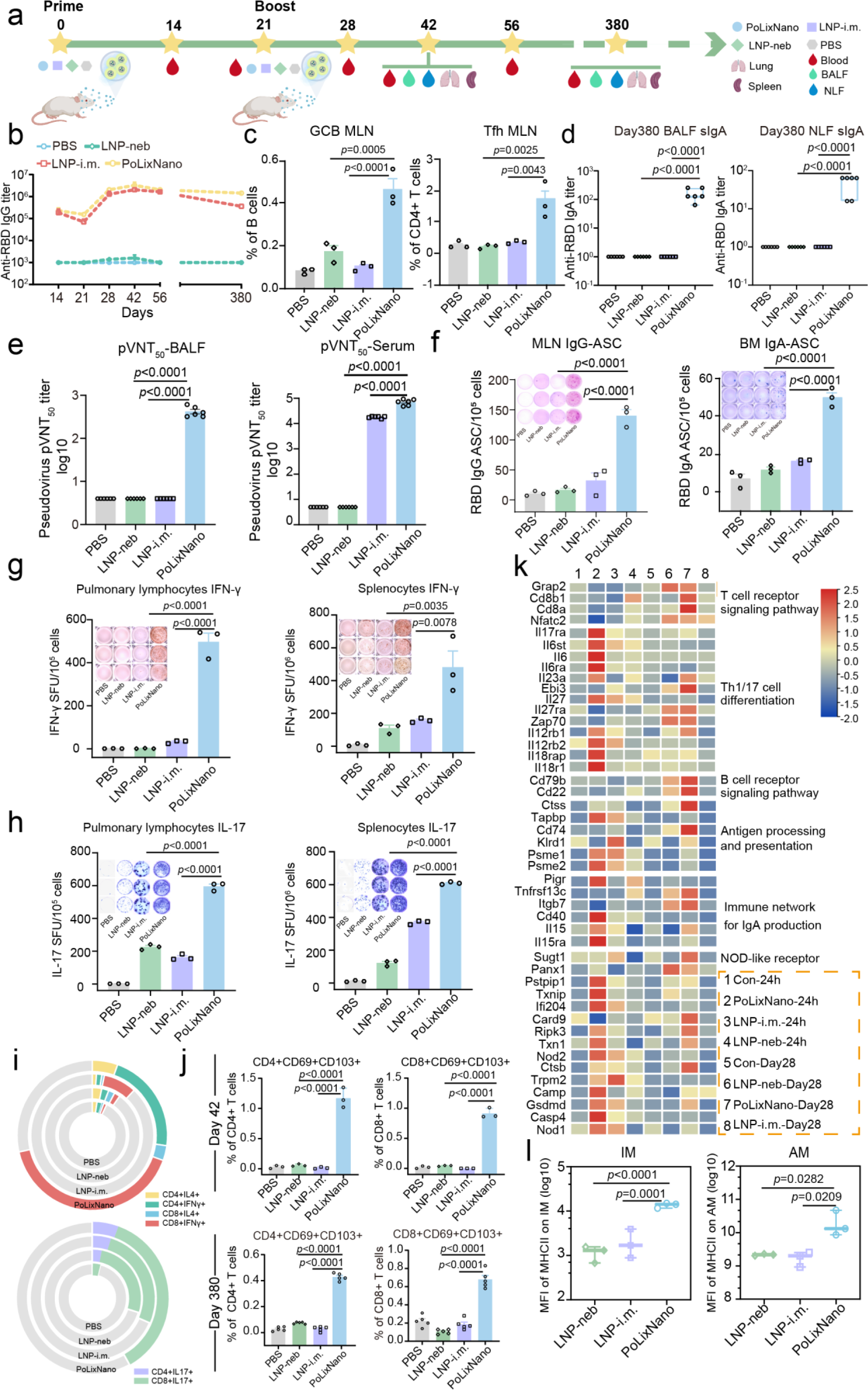
Immunologic evaluations of mucosal mRNA vaccine against SARS-CoV-2 mediated by the PoLixNano. **a**, Schematic illustration of immunization and sampling. BABL/c mice were immunized twice with RBD-mRNA encapsulated in PoLixNano (PoLixNano) or LNP (LNP-neb) via nebulization or intramuscularly injected LNP counterpart (LNP-i.m.) on day 0 and day 21. Mice treated by nebulized PBS were served as PBS control. **b**, The endpoint titers of RBD-specific immunoglobulin G (IgG) antibody in serum samples collected on day 14, 21, 28, 42, 56 and 380 post-prime via the ELISA analysis (n=9 biologically independent animals). **c**, Flow cytometry analysis of germinal center B cells (GCB) and T follicular helper cells (Tfh) in mediastinal lymph nodes (MLNs) on day 28 (n=3 biologically independent samples). **d**, The RBD-specific secretory immunoglobulin A (sIgA) antibody titer in the BALF and NLF samples collected on day 380 (n=6 biologically independent animals). **e**, Measurements of pseudo-virus neutralization titer 50 (pVNT_50_) from the serum and BALF samples of vaccinated mice on day 42 (n=6 biologically independent animals). **f**, Quantification of RBD-specific IgG and IgA antibody secreting cells (ASCs) in MLNs or bone marrow (BM) on day 35 via ELISpot analysis (n=3 biologically independent animals). Inserts: imagines of each well. **g**, **h**, ELISpot assay of the IFN-γ (**g**) and IL-17 (**h**) producing cells within the lung (left) and spleen (right) of mice treated by indicated formulations on day 42 after re-stimulation with a RBD peptide pool (details can be found in Supplementary Notes) (n=3 biologically independent animals). **i**, The proportion of RBD-specific intracellular Th1 (IFN-γ), Th2 (IL-4) or Th17 (IL-17) cytokine secreting CD8+ and CD4+ T cells in the lung on day 42 after re-stimulation with the RBD peptide pool (n=3 biologically independent animals). **j**, The proportion of tissue-resident memory T cells (T_RM_, CD69+CD103+) in the lung among CD4+ and CD8+ T cells on day 42 and day 380 (n=3 biologically independent samples on day 42 and n=5 biologically independent animals on day 380). **k**, A heatmap of immune pathway-related gene expression in lung samples collected at 24 h and 28 days post-prime. Lung samples from mice treated with PBS served as the control (Con). **l**, Mean fluorescence intensity (MFI) of MHC II expression on alveolar macrophages (AMs) and interstitial macrophages (IMs) from vaccinated mice at 42 days post-prime (n=3 biologically independent animals). A one-way ANOVA with Tukey’s post-hoc test was applied for statistical analysis of data in **c-h**, **j** and **l**.

We further evaluated the cellular immune responses induced by PoLixNano. ELIspot revealed that PoLixNano elicited more abundant interferon-gamma (IFN-γ) (**Fig. 4g**) and IL-17 (**Fig. 4h**) secreting cells than LNP-neb and LNP-i.m. counterparts, with background level of IL-4 secreting cells (**Supplementary Fig. 3a**). PoLixNano group simultaneously exhibited substantially increased proportions of IFN-γ (T helper 1 (Th1)-type cytokine) secreting- and IL-17 (T helper 17 (Th17)-type cytokine) secreting-CD4+ T cells and CD8+ T cells in pulmonary lymphocytes **(Fig. 4i** and **Extended Data Fig. 7a,b**). Unfortunately, all vaccinated groups failed to induce CD4+ or CD8+ T cells expressing IL-4 (**Extended Data Fig. 7c** and **Supplementary Fig. 3b**). Compelling evidences indicate critical roles of lung tissue resident memory T cells (T_RM,_ CD69+CD103+) in eliminating SARS-CoV-2^44,45^, we found RBD-mRNA@PoLixNano vaccinated mice demonstrated significantly higher proportion of CD8+T_RM_ and CD4+T_RM_ on both day42 and day380, compared to RBD-mRNA@LNP-neb and RBD-mRNA@LNP-i.m. counterparts (**Fig. 4j** and **Supplementary Fig. 4a**). Apart from that, we also observed an increased proportion of the effector memory CD4+ T cells (CD4+T_EM_, CD44+CD62L-), the central memory CD4+ T cells (CD4+T_CM_, CD44+CD62L+), CD8+ T_EM_ and CD8+ T_CM_ cells in the lung of PoLixNano vaccinated mice compared to other groups (**Extended Data Fig. 7d, 7e**), but PoLixNano did not show benefits on T_EM_ and T_CM_ in the spleen (**Supplementary Fig 4b**).

### Explorations on the underlying mechanism of the innate and adaptive immune responses induced by PoLixNano

RNA-sequencing (RNA-Seq) was adopted to investigate the underlying mechanism of the superior extent and duration of adaptive immune responses induced by PoLixNano (**Extended Data Fig. 8a**)., it revealed upregulated expression of T and B cell activation-associated genes in PoLixNano group, such as *cd8b1*, *cd8a*, *cd79b*, and *cd22* (**Fig. 4k)**. Since PoLixNano specifically up-regulated genes related to NOD receptor (*Nod2* and *Nod1*), Th1 cell differentiation (*Ifnlr1, Ifnar2, IL27, IL12rb1 and IL18r1*) and Th17 cell differentiation (*IL6*, *IL6ra* and *IL23a*) (**Fig. 4k** and **Extended Data Fig. 6f**), it may uniquely exert adjuvant effects via the Nod-Th1/Th17 axis signaling pathway (**Supplementary Fig. 5**).

PoLixNano vaccine showed distinct abundance of differentially expressed genes (DEGs) at 24 h and 28 days post-dosing compared to LNP-neb counterpart. There were 1544 upregulated genes and 2569 downregulated genes at 24 h, whereas 558 genes were upregulated and 1118 genes were downregulated after the second immunization at 28 days (**Extended Data Fig. 8b**,8c, left panel). Gene set enrichment analysis (GSEA) identifies enrichment of the *FOXN1* gene set at 24 h and the *PHC1* gene set at 28 days in PoLixNano group compared to LNP-neb (**Extended Data Fig. 8b**,8c, right panel). Notably, *FOXN1* is crucial for T cell development in postnatal thymic epithelial cells^48^. Venn analysis identifies 202 overlapping DEGs common to both PoLixNano and LNP-neb vaccinated groups at 24 h and 28 days (**Extended Data Fig. 8d**), and gene ontology (GO) analysis of these overlapping genes revealed that, compared to LNP-neb vaccinated groups, the upregulated genes in the PoLixNano vaccinated groups are primarily involved in interferon-gamma (IFN-γ)-mediated signaling pathway and cytokine-mediated signaling pathway signaling pathways at 24 h, including *PARP9*, *PARP14*, and *DDX58* (**Extended Data Fig. 8e**), which play important roles in regulating both innate and adaptive immunity^49–51^. Furthermore, weighted gene co-expression network analysis (WGCNA) isolated key gene modules associated with LNP-neb, LNP-i.m. or PoLixNano across different time points (**Extended Data Fig. 8f**), and reactome enrichment analysis further reveals that PoLixNano vaccinated group uniquely activated pathways including the “Adaptive Immunie System”, “Immunoregulatory Interactions between Lymphoid and Non-Lymphoid Cells”, “Antigen Processing and Cross-Presentation” and “Assembly and Peptide Loading of Class I MHC” at 24 h compared to LNP-neb or LNP-i.m. counterparts (**Extended Data Fig. 8g**).

Venn diagram analysis of temporal DEGs from 24 h to 28 days identifies specific genes associated with LNP-neb, LNP-i.m. and PoLixNano (**Extended Data Fig. 9a**). The kyoto encyclopedia of genes and genomes (KEGG) pathway analysis demonstrates that PoLixNano significantly regulated T helper 17 (Th17) cell differentiation, NF-κB signaling pathway, T cell receptor signaling pathway, MAPK signaling pathway, and antiviral defense pathways including those related to COVID-19, compared to LNP-neb and LNP-i.m. counterparts (**Extended Data Fig. 9b**). Combined with the observations that PoLixNano specifically upregulated the expression of genes related to NOD receptor (*Nod2* and *Nod1*), *Ripk3*, Th1 cell differentiation (*Ifnlr1, Ifnar2, IL27, IL12rb1, IL12rb2, IL18rap and IL18r1*) and Th17 cell differentiation (*IL6, IL6ra* and *IL23a*) (**Fig. 4k** and **Extended Data Fig. 6f**) and PoLixNano significantly promoted the number of IFN-γ and IL-17 secreting cells within the pulmonary lymphocytes (**Fig. 4h,4i**), the PoLixNano may exert adjuvant effects in the airway of mice via the Nod-Th1/Th17 axis signaling pathway **(Supplementary Fig. 5**). Previous studies has suggested that pathogen recognition mediated by the innate immune receptor Nod2 is crucial for the activity of mucosal adjuvants^52–54^. NOD-like receptors interact with Ripk family which in turn activates the NF-κB and MAPK signaling pathways to trigger the production of pro-inflammatory cytokines (such as IFN-γ, IL-4 and IL-6) and initiate immune responses^55,56^. It has been demonstrated that Th17 cells can differentiate and proliferate through the NOD receptor signaling pathway in the early stages of mucosal infection under the stimulation of cytokines such as IL-6 and IL-23^57^. Th17 cells primarily secret the characteristic cytokine IL-17 which plays a crucial role in the mucosal defence through multiple pathways, enhancing the tight junctions of epithelial cells in the mucosal area and promoting tissue repair at the site of injury^58^. After binding to the receptors of epithelial cells, IL-17 can trigger the production of cytokines such as Granulocyte-Macrophage Colony-Stimulating Factor (GM-CSF) and chemokines such as CCLs and CXCLs, recruiting neutrophils and natural killer cells to the site of mucosal infection to help eliminate invading pathogens^52^. Apart from that, IL-17 is also indispensable in mediating the generation of sIgA antibodies in respiratory mucosal site, the production of sIgA depends on the interaction between Th17 cells and B cells in the germinal center of pulmonary drainage lymph node^52,56,56,57,57–60^. All these clues align with our findings that PoLixNano vaccine induced high levels of mucosal sIgA antibody and robust Th17-biased T cell immunity (**Fig. 4d,4i and Extended Data Fig. 6c**,7a).

Our data also suggest PoLixNano vaccine substancially elevated the proportions of IFN-γ secreting CD4+ T cells (Th1 cells) and CD8+ T cells (cytotoxic T cells, CTLs) in the lung of vaccinated mice (**Fig. 4i** and **Extended Data Fig. 7b**). Th1 cells are crucial in controlling viral infection and cancer development, they preferentially produce IFN-γ and are the principal regulators of type I immunity against intracellular pathogens, such as invading virus and tumors^61^. IFN-γ stimulates recruitment of macrophages, neutrophils and T cells to the site of infection and plays a critical role in conveying antiviral signals from the innate to the adaptive immune response in order to fully activate host antiviral immunity^62^. Upon receiving the IFN-γ signal, antigen presenting cells increase expression of MHC class II and costimulatory molecules which in turn facilitate CD4+ T cell activation and initiation of the adaptive immune response against viral infection^63^. IFN-γ also drives cytotoxic T cell (CTLs) development. CD8+ CTLs can directly induce target cell death by the interaction between Fas/Fas ligand, and secretion of cytolytic mediator perforin, which creates pores in the target cells allowing the delivery of granule serine proteases (granzymes), to induce apoptosis^64^.

Furthermore, Venn diagram analysis of differentially expressed metabolites (DEMs) identifies specific metabolites for LNP-neb, LNP-i.m. and PoLixNano vaccinated groups, exhibiting temporal variations from 24 h to 28 days **(Extended Data Fig. 9c**). The KEGG enrichment reveals subcluster_3 was significantly associated with the PoLixNano group, including starch and sucrose metabolism, ABC transporters, galactose metabolism, carbohydrate digestion and absorption and cysteine and methionine metabolism (**Extended Data Fig. 9d**). The metaboAnalyst (https://www.metaboanalyst.ca) shows that PoLixNano-related DEMs were enriched in drug metabolism-cytochrome P450 and glycerolipid metabolism pathways, whereas LNP-related DEMs were primarily enriched in sphingolipid metabolism and lipoic acid metabolism pathways (**Extended Data Fig. 9e**).

The differentially expressed genes (DEGs) and differentially expressed metabolites (DEMs) related to the PoLixNano, LNP-neb, and LNP-i.m. vaccinated groups were further constructed into metabolic processes, with a visual analysis of the metabolic pathways conducted using iPath3.0 (http://pathways.embl.de). The results indicate that PoLixNano vaccinated group was involved in distinct metabolic processes compared with the other two LNP counterparts. (**Extended Data Fig. 9f**). Cytoscape visualization on the correlation between the DEMs and their associated genes reveals specific changes of drug metabolism, cytochrome P450, vitamin 6 metabolism, and fatty acid related metabolic pathways in PoLixNano group (**Extended Data Fig. 9g**). Notably, the significant down-regulation of cytochrome P450 related genes and metabolites suggests a decreased detoxification of xenobiotics was occurred in PoLixNano group, emphasizing the outstanding biocompatibility and stealthy properties of PoLixNano^46,47^.

Suggested by the above-mentioned RNA-seq results, we further found PoLixNano significantly activated (MHC II expression) both alveolar macrophages (AMs) and interstitial macrophages (IM) for trained innate immunity, while neither the LNP-neb nor LNP-i.m. showed benefits (**Fig. 4l**). Collectively, the above data indicate that PoLixNano is efficient in promoting a long-lasting and comprehensive immunity consisting of trained innate immunity, systemic, cellular, and mucosal immune responses.

### PoLixNano confers robust protection against lethal SARS-CoV-2 challenge and prevents transmission of Omicron strain

Given the promising immune responses elicited by PoLixNano, we assessed its protective effects against both a lethal SARS-CoV-2 ancestral strain C57MA14 (**Extended Data Fig. 10a**) and the Omicron BA.5 variant in mice respectively (**Fig. 5a**). Regarding the C57MA14 challenge, PoLixNano-vaccinated mice did not experience significant body-weight loss (**Extended Data Fig. 10b**) and exhibited robust protection with a 100% survival rate compared to the LNP-neb (0%), LNP-i.m. (75%), and PBS (0%) groups (**Extended Data Fig. 10c**). Viral RNA levels in the lung at 8 days post-challenge (dpc) and turbinate at 14 dpc were undetectable in the PoLixNano treated mice (**Extended Data Fig. 10d**), which is in stark contrast to LNP-neb and LNP-i.m. counterparts (**Extended Data Fig. 10e**). According to the histopathological assays, mice treated by LNP-neb or PBS not only displayed typical lung lesions, but also retained obvious amount of SARS-CoV-2 nucleocapsid protein (N protein) in the lung sections, whereas the pulmonary histopathologic lesions were significantly relieved in LNP-i.m. and PoLixNano groups with the absence of SARS-CoV-2 N protein (**Extended Data Fig. 10f**). Detection of cytokines in the lung homogenates further confirmed the above-mentioned observations, low levels of pro-inflammatory cytokines, including IL-4, TNF-α, IL-13, were only observed in the PoLixNano vaccinated mice (**Extended Data Fig. 10g**).

**Fig. 5.**
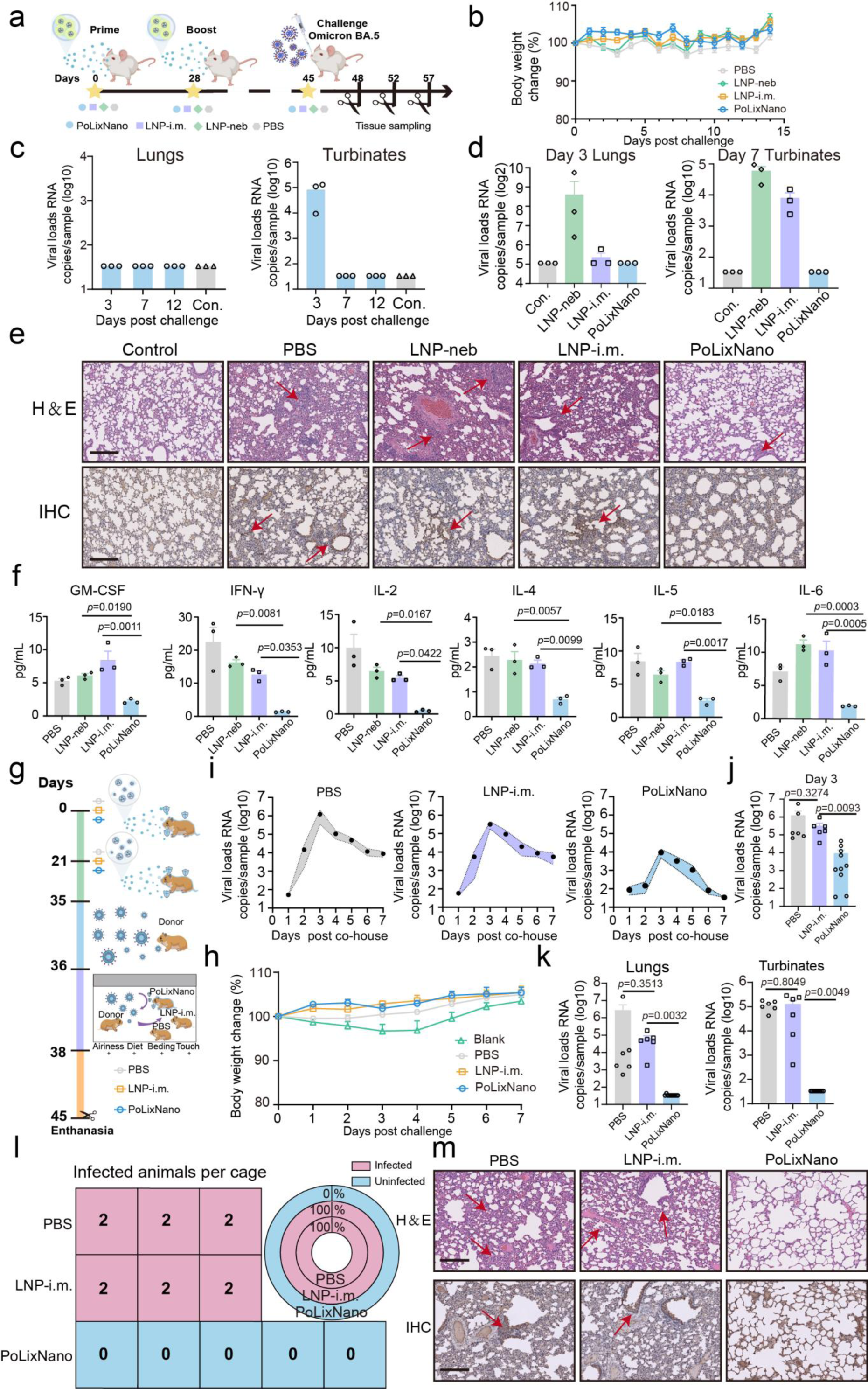
Robust protection conferred by PoLixNano-based mRNA vaccine against SARS-CoV-2 Omicron variant challenge in mice and its performance against the close contact transmission of Omicron variant in hamsters. **a**, Schematic illustration of immunization, sampling and live SARS-CoV-2 Omicron BA.5 variant challenge in mice. PoLixNano: nebulized RBD-mRNA@PoLixNano, LNP-neb: nebulized RBD-mRNA@LNP, LNP-i.m.: intramuscularly injected RBD-mRNA@LNP, PBS: nebulized PBS. **b**, Body weight changes of mice among all treated groups after the Omicron variant challenge (n=8 biologically independent animals). **c**, Viral RNA loads in the lung and nasal turbinate of mice treated by the PoLixNano at 3-, 7- and 12-days post-challenge (dpc). **d**, The viral RNA loads in lung tissues at 3 dpc and in nasal turbinate at 7 dpc of mice immunized with indicated formulations (n=3 biologically independent experiments). **e**, Histopathology (H&E) analysis (top) and immunohistochemical (IHC) assay of the SARS-CoV-2 nucleocapsid (N) protein (bottom) in lung tissues at 3 dpc. Red arrows indicate pathological changes and SARS-CoV-2 nucleocapsid (N) protein positive areas. Scale bars, 200 μm. Samples collected from untreated mice without challenge were served as the control (Con.) in **c**-**e**. **f**, Cytokine levels in lung homogenates collected from indicated groups at 3 dpc (n=3 biologically independent animals). **g**, Schematic diagram of the close contact infection assay using Omicron BA.5 variant strain. One donor hamster was administered with the live Omicron variant then co-housed with two recipient hamsters vaccinated by the same formulation (i.e. PBS, LNP-i.m. and PoLixNano) for 48 h close contact respectively, the recipient hamsters were analyzed for 7 days since co-house. **h**, Body weight changes of hamsters among all the indicated groups after contact with infected donor hamsters (n=6 biologically independent animals). The infected donor hamsters were used as the control group. **i**, Viral RNA loads measured in the nasal irrigation of PBS (n=6 biologically independent animals), LNP-i.m. (n=6 biologically independent animals) and PoLixNano (n=10 biologically independent animals) vaccinated recipient hamsters, data are shown as mean (symbols) and standard error of the mean (shade) over 7 days post-exposure (dpe) to the infected donors (n=11 biologically independent animals). **j**, The viral RNA loads in daily nasal irrigation of recipient hamsters at 3 dpe. **k**, Viral RNA loads in the lung homogenates and nasal turbinate were measured in recipient hamsters from PBS groups (n=6, biologically independent animals), LNP-i.m. (n=6, biologically independent animals), and PoLixNano (n=10, biologically independent animals) at 7 dpe. **l**, Numbers of infected recipient hamsters after exposure to hamster donor in each cage at 7 dpe. Infection rates of recipient hamsters in each group were calculated (Doughnut chart). **m**, The histopathology assay (H&E, top) and immunohistochemical (IHC) analysis of SARS-CoV-2 N protein (bottom) of lung tissues collected at 3 dpe. Red arrows indicate areas positive for the pathological lesions and SARS-CoV-2 N protein. Scale bars, 200 μm. The results of **e** and **m** are representative images from three independent experiments with similar results. Statistical analysis was determined by the one-way ANOVA with Tukey’s post-hoc test in **f** and by two-tailed unpaired *t*-test in **j** and **k**.

In terms of Omicron challenge, all vaccinated mice did not show body-weight loss or mortality (**Fig. 5b** and **Extended Data Fig. 10h**). PoLixNano-vaccinated mice were able to completely eliminate the virus in the lower respiratory tract as early as 3 dpc and in the upper airway 7 dpc (**Fig. 5c**), while obvious levels of viral loads still could be detected in LNP-neb and LNP-i.m. counterparts at the same time points (**Fig. 5d**). Lung sections from PoLixNano-treated mice exhibited occasional mild peribronchial inflammation and no detectable expression of SARS-CoV-2 N protein, whereas LNP-neb and LNP-i.m. counterparts exhibited abundant SARS-CoV-2 N protein, substantial thickening of alveolar septa and prominent infiltration of inflammatory cells (**Fig. 5e**). We also observed significant higher levels of pro-inflammatory cytokines (such as IL-2, IL-4, TNF-α, IL-6) in the lung homogenates of LNP-neb and LNP-i.m. groups compared to PoLixNano (**Fig. 5f** and **Extended Data Fig. 10i**).

Inspired by the robust protection of PoLixNano, we further utilized hamsters to assess its potential benefits against host-host transmission of Omicron BA.5 strain in a close contact transmission protection model (**Fig. 5g**). After exposure to the donor, no weight loss was observed in the exposed hamsters (**Fig. 5h**). The viral loads in nasal irrigation samples collected from PBS and LNP-i.m. inoculated hamsters were drastically increased to above 10^5^ RNA copies/sample during 1 to 3 days post-exposure (dpe) without significant sign of relief even 7 dpe (**Fig. 5i**), whereas PoLixNano counterparts displayed 33-131 times lower viral loads at 3 dpe (**Fig. 5j**) and declined to background level 7 dpe. PoLixNano-vaccinated hamsters exhibited the lowest viral loads in lung homogenates and nasal turbinate 7 dpe compared with PBS and LNP-i.m. counterparts (**Fig. 5k**). It is worth to note that 100% of hamsters inoculated with PBS or LNP-i.m. were infected even 7 dpe, whereas no infections were observed in PoLixNano vaccinated group (**Fig. 5l**). Mild lung lesions and obvious SARS-CoV-2 N protein expression could be detected in PBS and LNP-i.m. group 7 dpe, while PoLixNano vaccination displayed robust protection against lung pathology, without detectable SARS-CoV-2 N protein (**Fig. 5m**).

Although pioneering studies have demonstrated intranasal vaccination with COVID-19 mRNA induced potent immune protection against SARS-CoV-2 challenge^9,10^, our findings for the first time suggest that mucosal mRNA vaccines confers substantial benefit against host-host transmission of the Omicron variant apart from inhibiting disease development. These encouraging data may expedite the clinical translation process of mucosal mRNA vaccines and potentially enable us to be better prepared in combating the outbreak of future pandemics.

### PoLixNano displays versatility in lung cancer treatment and cystic fibrosis gene therapy

To demonstrate the versatility of PoLixNano as a cancer therapeutic to inhibit lung metastasis, we established a B16F10-based luciferase expressing cells (B16F10-luc) induced lung metastatic melanoma model and employed mRNA encoding interleukin-12 (IL-12-mRNA) which demonstrates robust potency in lung tumor suppression^13^ (**Fig. 6a**). The PBS and LNP-neb treated mice showed drastically weight loss since Day12, whereas PoLixNano treatment prominently alleviated this phenomenon (**Fig. 6b**). Monitoring B16F10-luc tumor development further revealed that PoLixNano significantly suppressed tumor progression (**Fig. 6c,6d**), corresponding to the delayed onset of mortality in mice treated by PoLixNano with more than 50% mice survived at the end of investigation (**Fig. 6e**). On the other hand, PBS and LNP-neb treated mice experienced early mortality, with an averaging 26-day lifespan (**Fig. 6e**). In comparison to LNP-neb counterpart, PoLixNano treatment substantially reduced the number of metastatic foci and lung weights (**Fig. 6f,6g**). Owing to the therapeutic benefits of PoLixNano, the expression of a melanocyte-specific gene *Tyrp1*, which represents a quantitative indicator of B16F10 tumor metastasis and progression in the lung^65^, significantly mitigated in PoLixNano treated samples in contrast to the LNP-neb and PBS controls (**Fig. 6h**).

**Fig. 6.**
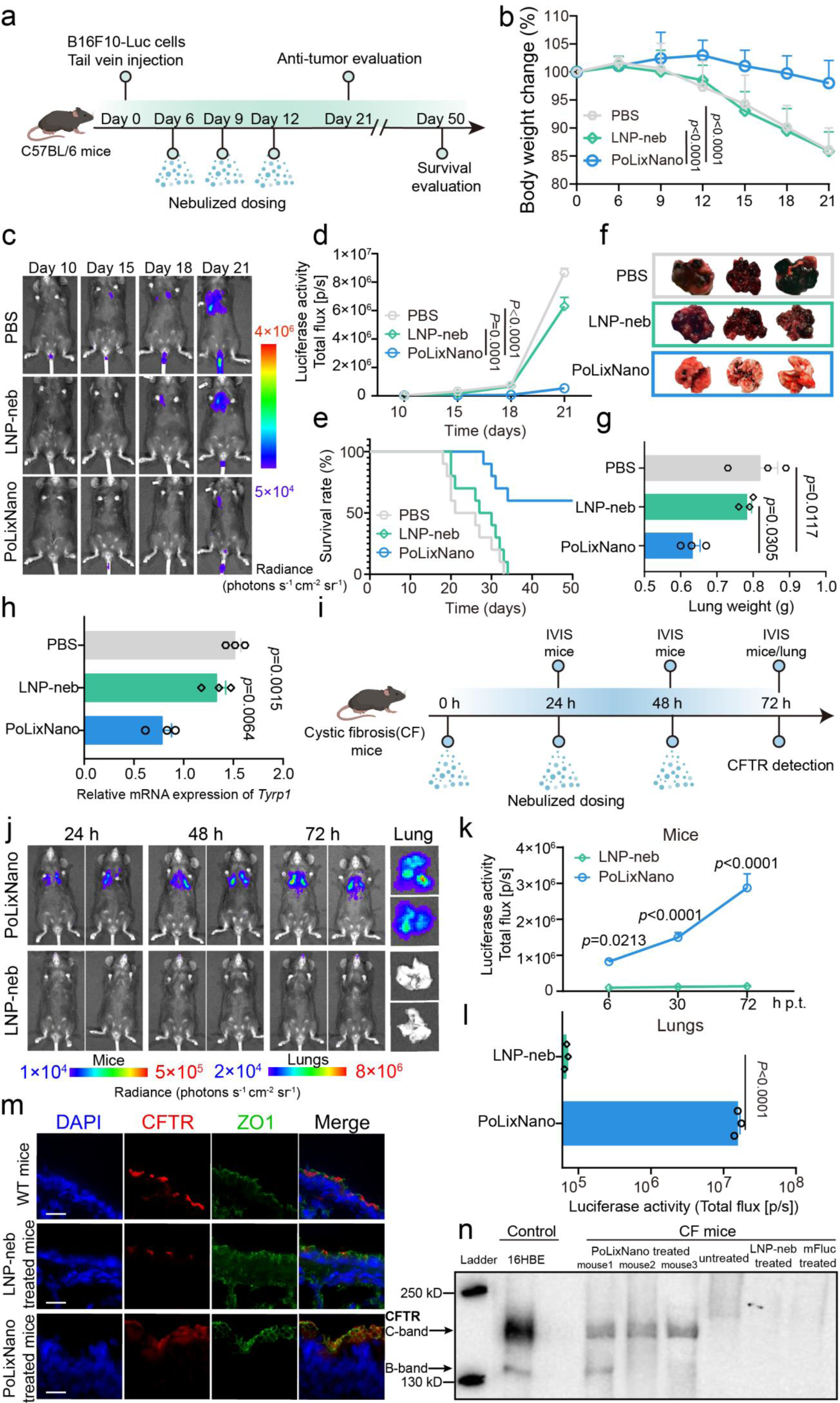
Evaluations on therapeutic efficacy of nebulized PoLixNano in metastatic tumor and cystic fibrosis (CF) models. **a,** Schematic illustrating the establishment of the B16F10 melanoma lung metastatic tumor model and anti-tumor efficiency assessments after mice were nebulized with PBS, LNP (LNP-neb) and PoLixNano. **b**, Body weight changes over time for mice treated by indicated formulations (n=10 biologically independent mice). **c**,**d**, Bioluminescence imaging (**c**) of B16F10-luc tumor growth over time in mice treated by indicated formulations, and the corresponding quantification (**d**) of bioluminescence intensities at the indicated time points (n=10 biologically independent mice). Statistical significance was evaluated on day 21 in **b** and **d** using one-way ANOVA with Tukey’s post-hoc test. **e**, Survival curves of mice for each treatment group (n = 10 biologically independent mice). **f**, Representative morphologies of lungs excised at day 21 post-tumor inoculation. Black points represent the metastatic foci (n = 3 biologically independent samples). **g**, Lung weights measured at 21 days after tumor inoculation (n=3 biologically independent animals). **h**, Quantitative real-time PCR analysis on the mRNA expression of tyrosine phosphatase type 1 (Tyrp1) at 21 days post-tumor inoculation (n = 3 biologically independent samples). **i**, Schematic depicting the dosing regimen in cystic fibrosis mouse model of inhaled Fluc-mRNA or CFTR-mRNA, encoding cystic fibrosis transmembrane conductance regulator (CFTR), encapsulated in PoLixNano or LNP (LNP-neb). **j**, Representative bioluminescence images of Fluc-mRNA expression mediated by PoLixNano or LNP-neb in live CF mice over time and in excised lungs 72 h post first dosing. **k**, Quantification of bioluminescence intensities in live CF mice over time as described in (j). (n = 3 biologically independent mice). **l**, Quantitative analysis of bioluminescence signal intensities of excised lungs collected in (j). (n = 3 biologically independent samples). **m**, Representative immunofluorescence staining of CFTR protein in the lung sections collected from CFTR-mRNA@LNP-neb or CFTR-mRNA@PoLixNano treated CF mice. Samples obtained from untreated wild-type mice (WT) were used as the positive control. Red channel: CFTR, green channel: zonula occludens-1 (ZO1), blue channel: nuclei. Scale bars: 10 µm. **n**, Western blot analysis of CFTR expression in lungs of CFTR-mRNA@PoLixNano (PoLixNano lanes) or CFTR-mRNA@LNP-neb (LNP-neb lane) treated CF mice. 16HBE cells, which endogenously express wild-type human CFTR, were used as a positive control (16HBE lane). Untreated CF mice (untreated lane) and Fluc-mRNA@PoLixNano (mFluc lane) treated CF mice were served as negative controls. Signal detection was performed using Pierce Femto, with 30 seconds detection for 16HBE and 60 seconds detection for all mouse samples. The results of **c**, **j** and **m** are representative images from three independent experiments with similar results. Statistical significance was analyzed using a one-way ANOVA with Tukey’s post-hoc test in **b**, **d**, **g** and **h**. A two-way ANOVA with Sidak’s multiple comparisons test was applied in **k**, and a two-tailed unpaired *t*-test was adopted in **l** for calculating the statistical significance.

To evaluate the potential of PoLixNano in protein-replacement gene therapy, we investigated the mRNA expression profiles in a cystic fibrosis transmembrane conductance regulator (*CFTR*) knock-out mouse model (CF mice). Considering repeated dosing is necessary in cystic fibrosis (CF) gene therapy to maintain the CFTR expression within the therapeutic window, we first employed Fluc-mRNA to test whether the PoLixNano can be given repeatedly (**Fig. 6i**). It turned out that Fluc-mRNA expression levels of PoLixNano group were maintained (**Fig. 6j-l**). Three repeated doses of mRNA encoding CFTR (CFTR-mRNA) were administered to CF mice via nebulization (**Fig. 6i**). CFTR immune-staining revealed consistent and clear expression of CFTR on the apical side of the superficial epithelium of bronchial airways in CF mice treated by PoLixNano, which was comparable to the wild-type counterparts but was in contrast to LNP-neb counterparts that display faint CFTR signal (**Fig. 6m**). Western blot analysis further confirmed samples from PoLixNano treated mice displayed strong signals at approximately 170-180 kDa in size, which were similiar to the C-band signal (mature CFTR) of 16HBE controls that endogenously express wild-type CFTR^14^ (**Fig. 6n**). However, no CFTR signal was observed in untreated CF mice, nor in CFTR-mRNA@LNP-neb and Fluc-mRNA@PoLixNano (mFluc) treated CF mice (**Fig. 6n**).

## Conclusion

In summary, we have developed a safe and effective delivery strategy, i.e. PoLixNano, for nebulized mRNA via leveraging machine learning algorithms. Our dataset reveals the crucial roles of the “shield” and “spear”-like properties for inhaled mRNA therapeutics. By virtue of these barriers-overcoming properties, the nebulized PoLixNano display promising mRNA transfection in a serial of in vitro and in vivo models without observable risk of vehicle-induced pathology. We also systematically demonstrated PoLixNano is a versatile platform in mediating nebulized mRNA therapeutics to address the transmission of airborne pathogen, lung cancer and congenital lung disease such as cystic fibrosis. Although further validation and optimization for human use are still needed, we believe PoLixNano technology would be one of the cutting edge in vivo nebulized mRNA delivery systems and exhibit potential to revolutionize the inhaled mRNA therapeutics ranging from mucosal vaccines to gene therapy for treating refractory pulmonary diseases.

## Methods

### Reagents

Nuclease-free water (129117) was purchased from QIAGEN. Opti-MEM I reduced serum medium (31985070), riboRuler high range RNA ladder (SM1821), lithium chloride (AM9480), protein transport inhibitor cocktail (00-4980-93), 1,1’-dioctadecyl-3,3,3’,3’-tetramethylindodicarbocyanine perchlorate (DiD) (D7757) and LysoTracker Red DND-99 (L7528) were purchased from Thermo Fisher Scientific. Agarose (A118880) and Pluronic block copolymers (P181 (P131343), P184 (P131345), P401 (P131346), P10R5 (P434424) and L31 (P131340)) were purchased from Aladdin. 3-Morpholinopropanesulfoinc Acid (MOPS) buffer (M1010) was purchased from Solarbio. Pre-rinsed dialysis bags with a molecular weight cut-off of 10-14kDa were purchased from Viskase. The ionizable lipid SM-102 (O02010), ALC-0315 (O02008), DLin-MC3-DMA (O02006) and dipalmitoylphosphatidylcholine (DPPC) (S01004) were purchased from AVT (Shanghai, China). 1,2-distearoyl-sn-glycero-3-phosphocholine (DSPC) (850365P), 1,2-dioleoyl-sn-glycero-3-phosphoethanolamine (DOPE) (850725P), 1,2-dioleoyl-sn-glycero-3-phosphocholine (DOPC) (850375P), 1-stearoyl-2-oleoyl-sn-glycero-3-phosphocholine (SOPC) (850467P), 1,2-dimyristoyl-rac-glycero-3-methoxypolyethylene glycol-2000 (DMG-PEG2000) (880151P), campesterol (70012P6), stigmasterol (700062P), DSPE PEG2000 (880128P), ALC-0159 (880155P), 14:0 PEG1000 (880710P), 14:0 PEG3000 (880310P) and 14:0 PEG5000 (880210P) were purchased from Avanti Polar Lipids. Poloxamine 704 (T704) was a gift from InCellArt (Nantes, France). The poloxamine (tetronic) block copolymers (T304, T707, T803, T901, T908, T1107, T1301, T1304, T1307, T1508 and T150R1) and Pluronic P85 were kindly gifts from BASF GmbH (Ludwigshafen, Germany). Poloxamine T90R4, poloxamine T904 (435546), poloxamine T701 (435511), cholesterol (C3045), fucosterol (F5379), β-sitosterol (43623), dioleoyl Phosphatidylserine (DOPS) (840035P), Pluronic block copolymers (P407 (P2443), P188 (15759), P338 (P2164021), P105 (435414), P124 (P2163920), P84 (713538), P237 (P2164020), P401 (435457) and P335(86216)), various poly(ethylene glycol) (PEG) compositions (4 arm PEG compositions (2K(JKA7032), 5K(JKA7109), 10K(JKP2003R), 20K(JKP2005R)), 8 arm PEG compositions (10K(JKA8008), 20K (JKA8009), 40K (JKA8012)), PEG6K (901397), PEG20K (81298)) and various poly(propylene glycol) (PPG) compositions (PPG400 (81350), PPG1K (202320), PPG2K (202339) and PPG4K (202355)) were purchased from Sigma-Aldrich. PPG600 (P874988) and PPG1K (P815573) were purchased from Macklin. D-Luciferin potassium salt (122799) was purchased from PerkinElmer. CFTR antibodies were obtained for J. Riordan (University of North Carolina at Chapel Hill). All other solvents and small molecular reagents were obtained at analytical or HPLC grade from Sigma-Aldrich.

### In vitro transcribed mRNA

In vitro transcribed messenger RNA (IVT mRNA) were prepared and purified as previously described^1,2^. Briefly, IVT mRNA (the open reading frame (ORF) sequences of IVT mRNA used in this study can be found in Supplementary Table 1) was synthesized by linearized plasmid templates using the T7 High Yield RNA Synthesis Kit (E131-01A, novoprotein) with N1-methylpseudouridine (Glycogene, Wuhan, China) substitution and followed by the addition of a Cap 1 structure using Cap 1 Capping System (M082-01B, novoprotein). The synthesized IVT mRNA was purified using an affinity chromatography-based purification^1^. The concentration and purity of IVT mRNA were measured using a NanoDrop^TM^ One (Thermo Fisher Scientific).

### Preparation of PoLixNano and LNP

PoLixNano was prepared by mixing an aqueous phase with an ethanol phase using a microfluidic device (AceNANOSOME, ACMEI LIFESCIENCE, Shanghai, China) at a volume ratio of 3:1 with a total flow rate of 12 mL/min. Ethanol phase: lipid components, including ionizable lipid (SM-102, DLin-MC3-DMA or ALC-0315), helper phospholipid (DOPC, DOPE, DSPC, DPPC or SOPC), sterol (fucosterol, β-sitosterol, cholesterol, campesterol or stigmasterol) and PEG lipid (ALC-0159, DSPE-PEG2000, DMG-PEG2000, 14:0 PEG1000, 14:0 PEG3000 or 14:0 PEG5000), were dissolved in ethanol with a molar ratio of 45:20:33.5:1.5, or at specific ratios as indicated in certain investigations. Optionally, hydrophobic polymeric components were dissolved in ethanol phase. Aqueous phase: IVT mRNA and corresponding hydrophilic or amphiphilic polymeric components were dissolved in citric acid buffer (50 mM, pH=4). The nitrogen to phosphorus (N/P) ratio between IVT mRNA and ionizable lipid (SM-102, DLin-MC3-DMA or ALC-0315) was 8. The nebulized LNP counterparts were prepared as same as PoLixNano but without the addition of polymeric components. For intramuscular route administered LNP, lipid components were composed of DLin-MC3-DMA, DSPC, cholesterol and DMG-PEG2000 at a molar ratio of 50: 10: 38.5: 1.5, with a N/P ratio of 6. DiD labelled nanoparticles (DiD-PoLixNano or DiD-LNP) were prepared by adding DiD to the ethanol phase at 1 mol % relative to the total lipid. The PoLixNano and LNP were dialyzed against 1×PBS (pH 7.4) overnight using dialysis bag with molecular weight cutoff of 8-14 kDa at 4 °C for 16 h before further use. For iLNP-HP08^LOOP^ formulation^3^, lipid components were composed of ALC-0315, DSPC, cholesterol and DMG-PEG2000 at a molar ratio of 60: 20: 19: 1. The iLNP-HP08^LOOP^ were dialyzed against a dialysis buffer composed of 20 mM HEPES at pH 6.0 for 16 h. P188 was added to the iLNP-HP08^LOOP^ at a final concentration of 8 mg/mL for subsequent nebulization.

### Characterizations of the PoLixNano formulations

The particle size and zeta potential of the PoLixNano were determined using a Zetasizer Nano ZS (Malvern Instruments) at 25 °C. The morphology of PoLixNano nanoparticles was investigated by transmission electron microscope (TEM, JEM-1400plus, JEOL Ltd., Japan). The viscosity of the PoLixNano was measured by a viscometer (HAAKE^TM^ MARS^T^, Thermo Fisher Scientific). The encapsulation efficiency (EE) of IVT mRNA within PoLixNano was assessed using a Quant-iT^TM^ RiboGreen RNA Assay kit (R11490, Thermo Fisher Scientific). PoLixNano samples were dispersed in 1 ×Tris-EDTA (TE) buffer or 2 % Triton X-100 surfactant respectively, incubated at 37 °C for 30 min, and diluted with 1×TE buffer in final concentration of 0.04 µg/mL to 2 µg/mL. RNA standard solutions was prepared in five concentrations (0 µg/mL, 0.04 µg/mL, 0.2 µg/mL, 1 µg/mL, and 2 µg/mL) using 1×TE buffer. 100 μL PoLixNano samples or RNA standard solutions were mixed with 100 μL of RiboGreen reagent (1:200 diluted) in a clear-bottom black plate and incubated at room temperature (RT) for 3 min. Fluorescence intensity was measured using a microplate reader (varioskan^TM^ LUX, Thermo Fisher Scientific) at an excitation wavelength of 480 nm and an emission wavelength of 520 nm. The mRNA EE was calculated as follows: EE (%) = [(concentration of total IVT mRNA - concentration of free IVT mRNA) / concentration of total IVT mRNA] ×100%).

### Nebulization of IVT mRNA formulations

The PoLixNano or LNP formulations containing IVT mRNA were nebulized using a vibrating mesh nebulizer (Aerogen^®^ Pro, Aerogen, Ireland). For in vitro investigations, a fraction of formulations were kept apart and used as a “pre-nebulized” sample. The rest of the formulations were aerosolized, and nebulized solution (“nebulized” sample) was collected in a separate tube for further use. In terms of in vivo studies, the mice were loaded into custom-built nose-only exposure chamber with animal restraints. The Aerogen^®^ Pro Nebulizer was placed on the upward-facing the exposure chamber. The formulations were added dropwise to the nebulizer at a flow rate of 100 μL per droplet. After each individual droplet was nebulized, the exposure chamber was inspected until the vaporized dose had cleared (approximately 60-120 s per drop). The droplets were added until the desired dose per mouse was achieved, and the mice were removed from the restraints for further evaluations. For the inhalation of IVT mRNA to rats and hamsters, the exposure system was modified with larger animal restraints and a larger nose cone to fit the size of animals. For the delivery of IVT mRNA to the rabbits, a modified nebulizer mask was used along with a larger restraint.

### Machine learning

The PoLixNano library were generated using a series of polymeric components (Supplementary Table 2). The models that we developed were from the conda installable package Sci-kit Learn in Python 3.9.19 and developed in JetBrains PyCharm. The linear regression model was evaluated on ten-folds cross-validation by R-Squared. All of the final models in use were trained on 70% dataset and tested on the remaining 30%. Nebulization shear force deformation index (NDI) was calculated by size, polydispersity index (PDI), encapsulation efficiency (EE) before and after nebulization using equation 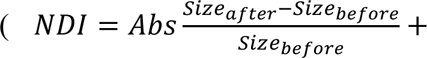 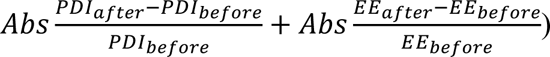. To compare the contribution of each feature to transfection efficiency, the diffusion coefficient (Deff), NDI, size, PDI and EE were normalized by a StandardScaler from Sci-kit Learn. We employed a multiple linear regressor to model the relationship between the physicochemical properties of PoLixNano and Log_10_ (transfection efficiency). The contribution of each feature was represented by calculating the proportion of its regression coefficient relative to the sum of all regression coefficients. Chemical structures for all monomers in simplified molecular-input line-entry system (SMILES) representations were extracted from Pubchem. Compounds were described according 12 calculated physicochemical properties (rdkit.Chem.Descriptors._descList, rdkit.org, details can be found at: https://github.com/rdkit/rdkit/issues/3105) for each monomer in polymers and 6 manually labeled features (molecular weight of polymer, arms, degree of each monomer and the polymer concentration used to prepare PoLixNano). These 30-dimensional numerical characterizations of a polymer served as input for KNR, RF, XGB, Ridge, Multi-RF and Multi-XGB implemented with optimized parameters in scikit-learn. For Deff, NDI, size, PDI and EE of PoLixNano, we compared the performance of different regression models on five-fold cross-validation and selected the most suitable model for each of them as the component to be incorporated into the final screening and prediction platform. The feature importance to Deff, NDI, size, PDI and EE were determined by calculating the mean importance of the best-suited model using five-fold cross-validation.

### Agarose gel retardation assay

The binding affinity and stability of IVT mRNA encapsulated in PoLixNano or LNP were evaluated by gel retardation assay. The electrophoresis was conducted with 1% (w/v) agarose gel containing 200 ng/well of naked IVT mRNA and 4 µg/well of IVT mRNA encapsulated in PoLixNano or LNP, both before and after nebulization. All samples were mixed with equal volumes of 2×loading dye buffer (R0641, Thermo Fisher Scientific), and loaded into the formaldehyde denaturing gels in 1×MOPS buffer, and followed by electrophoresis at 120 V for 20 min. For the IVT mRNA release study, PoLixNano or LNP samples were incubated with different concentrations of sodium dodecyl sulfonate (SDS) (28364, Thermo Fisher Scientific) solution (8 mM to 25 mM) at RT for 30 min. The gel electrophoresis was performed as described above.

### Multiple-particle tracking (MPT) analysis

3% (w/v) porcine mucin (M2378, Sigma-Aldrich) was dissolved in nuclease-free water and stirred overnight at RT to obtain mucin gel mimicking mucus layer. 20 μL of DiD-PoLixNano or DiD-LNP was incubated within 400 µL of mucin gel for 30 min at 37 ℃. The motion of nanoparticles in mucus solution was recorded using a Leica SP8 (Leica, Germany) microscope equipped with a 40×water objective. Fifteen-second videos were captured using the LAS4.5 software (Leica). Three-dimensional time-lapse microscopy images were processed using Imaris software (imaris version 9.8.2, Oxford Instruments). Images underwent deconvolution and background subtraction before particle detection and tracking. Tracks were reconstructed using the autoregressive motion algorithm, with quality filters applied to exclude spurious trajectories. The resulting particle trajectories, containing spatial coordinates (X, Y, Z) and temporal information for each detected particle, were exported as CSV files for subsequent analysis.

Particle trajectories were analyzed using custom R scripts (R version 4.4.1). Raw tracking data was imported from Excel files containing particle positions (X, Y, Z coordinates) and time points. For each particle track, we performed a comprehensive motion analysis that included calculation of step-wise displacement between consecutive positions (δr = √[(x₂-x₁)² + (y₂-y₁)² + (z₂-z₁)²]), cumulative displacement along the trajectory (Σδr), and total displacement from the origin in three dimensions (Δr = √[Δx² + Δy² + Δz²], where Δx = x - x₀, Δy = y - y₀, Δz = z - z₀). Mean Square Displacement (MSD) analysis was conducted by calculating the square of the total displacement from the origin (⟨ r²(τ)⟩ = ⟨ [r(t + τ) - r(t)]²⟩), with results normalized by time differences (τ) from the start of each track. These measurements were averaged across all particles within each experimental group to generate representative mobility profiles. Diffusion coefficients were subsequently derived using the Einstein relation for three-dimensional diffusion (⟨ r²(τ)⟩ = 6Dτ), where D is the diffusion coefficient (Deff) and τ is the time interval. Values were determined through linear regression of MSD versus time, with the diffusion coefficient calculated as D = slope/6. Standard errors (SE) were calculated from the regression analysis using SE(D) = SE(slope)/6 to assess the precision of these measurements. For visualization, trajectories were normalized to common starting points and projected onto optimal two-dimensional planes determined by the principal axes of motion, and plotted via Prism 8.0.2. All analyses were performed using the tidyverse suite of R packages with error bars in all plots representing standard error of the mean unless otherwise noted.

### Mucus-penetration in three-dimensional (3D) fluorescence labeled mucus-simulating hydrogel

The mucus-penetrating capacity of PoLixNano was evaluated in an artificial mucus model. Briefly, Rhodamine B (1 μg/mL) (A13572-30, Thermo Fisher Scientific) was added to the mucin gel (3% (w/v) porcine mucin) mimicking mucus layer and 150 μL of the mixture was transferred into each well of 96-well plate. Subsequently, 50 μL of PoLixNano (20 μg/mL of Fluc-mRNA), formulated with FITC-DOPE (M31161, AbMole), was applied to the apical side of the well and incubated at 37 ℃ for 10 min. The movement of PoLixNano through the mucus was imaged using confocal microscope (FV3000, Olympus, Japan) and analyzed via a live cell imaging system (IX83, Olympus, Japan).

### Mucus-penetration in mucus secreting cell model

Calu-3 human lung adenocarcinoma (Calu-3) cells were seeded into the apical side of the PET trans-well inserts of 24-well plates (3524, Corning) at a density of 1×10^4^ cells. Calu-3 cells were cultivated for up to 21 days in order to produce a thick mucus layer. The cell nuclei was labelled by adding 4′,6-diamidino-2-phenylindole (DAPI, D1306, Thermo Fisher Scientific) for 15 min. Subsequently, 20 μL of PoLixNano (50 μg/mL of Fluc-mRNA) formulated with FITC-DOPE was added into the apical chambers and incubated at 37 ℃ for 30 min. The distribution of PoLixNano in the mucus layer was visualized using the z-stack mode of confocal laser scanning microscope (Zeiss LSM880, Germany).

### Transepithelial transport ability assays

#### Trans-well insert model

A Trans-well insert model was established to investigate the transepithelial ability of PoLixNano across the mucus layer and airway epithelial cells. Briefly, Calu-3 cells were seeded at a density of 3×10^4^ cells in the apical side of the PET trans-well inserts of 24-well plates. On day 21 post-seeding of Calu-3 cells, DC 2.4 cells (1×10^5^ cells/well) were seeded into the basolateral compartment. PoLixNano or LNP encapsulating Fluc-mRNA (400 ng/well) were added to the apical chambers. After 24 h incubation, DC 2.4 cells were washed three times with 1×PBS and luciferase activity was measured using a luciferase assay kit (RG006, Beyotime Biotechnology). Firefly luciferase activity was recorded on a microplate reader (GloMax, Promega).

#### Ussing Chamber assay

The Ussing Chamber system was used to test the transepithelial transporting ability of PoLixNano^4,5^. A thin film of excised bronchial tissue from rats was evenly distributed in the space between a pair of semi-penetrating membrane filters (Merck Millipore, 2.0 μm) placed between the donor and acceptor chambers. 3 mL of HEPES solution (1×10^-3^ M, pH 7.4) was added to the receptor chamber, and an equal volume of PoLixNano (10 μg/mL of Fluc-mRNA) formulated with FITC-DOPE was added to the donor chamber. At indicated time points, 0.2 mL samples were collected from the receptor chamber for detection.

### Cell culture

Human bronchial epithelial cells 16BE14o-(16HBE) and DC2.4 (a mouse dendritic cell line) were cultured in complete medium comprising PRMI 1640 (11875093, Gibco) supplemented with 10% fetal bovine serum (FBS, A5670701, Gibco) and 1% penicillin/streptomycin (15140122, Thermo Fisher Scientific). B16F10-Luc cells, Calu-3 cells and A549 (a human non-small cell lung carcinoma cell line) cells were cultured in Dulbecco’s Modified Eagle Medium (DMEM, 11965118, Thermo Fisher Scientific) with 10% FBS and 1% penicillin/streptomycin. All cell lines were passaged using 0.25% trypsin-EDTA (25200072, Gibco) when they reached 90% confluence. B16-F10 transgenic cell line expressing firefly luciferase (B16F10-Luc cells) were purchased from Cellverse (shanghai, China), and other cell lines were obtained from the American Type Culture Collection (ATCC, Manassas, VA, USA). All cell lines were tested negative for mycoplasma contamination. All experiments were performed on cells in the logarithmic growth phase.

### Cellular uptake and in vitro IVT mRNA transfection

Fluorescence labeled IVT mRNA (Fluorescein-mRNA) was prepared using Label IT^®^ Nucleic Acid Labeling Kit (MIR 3200, Mirus bio, USA) according to the instructions. For cellular uptake study, DC2.4 cells (1×10^5^ cells/well), 16HBE cells (1.5×10^5^ cells/well), A549 cells (2×10^5^ cells/well) and bone marrow-derived dendritic cells (BMDCs, 5×10^5^ cells/well) were seeded on 24-well plates and incubated overnight. Once the cells reached approximately 70% confluence, PoLixNano encapsulating 200 ng of Fluorescein-mRNA was added to each well. The cells were incubated for 4 h and washed twice with PBS for analysis of fluorescein positive cells via flow cytometer (BD FACSCanto^TM^ II, BD Biosciences). To further investigate the internalization mechanism of PoLixNano, DC2.4 cells (1×10^5^ cells/well), 16HBE cells (1.5× 10^5^ cells/well) or BMDCs (5×10^5^ cells/well) were treated with specific inhibitors (0.5 mg/mL sodium azide, 0.5 mM amiloride, 1 μg/mL β-cyclodextrin, 1 mM protamine sulfate or 10 μg/mL chlorpromazine) at 37 ℃ or pre-incubated at 4 ℃ for 1 h. Subsequently, Fluorescein-mRNA@PoLixNano (200 ng mRNA/well) was added to the above pre-treated cells in the presence of inhibitors, followed by incubation at 37 ℃ or at 4 ℃ for an additional 4 h. The cellular uptake of the Fluorescein-mRNA@PoLixNano without any treatments was adopted as the positive control. The relative uptake rate of various groups was normalized according to the positive control. The fluorescein positive cells were analyzed via flow cytometry (BD FACSCanto^TM^ II, BD Biosciences).

To investigate the subcellular fate of PoLixNano, 16HBE cells were seeded at a density of 3× 10^5^ cells/well in 6-well plates (3516, Corning). The cells were incubated with Fluorescein-mRNA@PoLixNano (1.5 µg IVT mRNA/well) or LNP counterpart (1.5 µg IVT mRNA/well) for 4 h, then subjected to the incubation with LysoTracker™ Red DND-99 (L7528, Invitrogen™) for 30 min. The cell nuclei was labelled with DAPI for 15 min. The fluorescence imaging was performed using confocal microscope (FV3000, Olympus, Japan).

To explore the transfection efficiency of PoLixNano, 16HBE cells (3.5×10^4^ cells/well), DC2.4 cells (2×10^4^ cells/well), A549 cells (2.5×10^4^ cells/well) and BMDCs (3×10^4^ cells/well) were seeded in 96-well plates (3599, Corning) 24 h before transfection. 170 μL of serum-free Opti-MEM was added each well, followed by the addition of 30 μL of PoLixNano (400 ng of MetLuc-mRNA per well), in triplicate or more. After 6 h of incubation, the medium was replaced with complete medium and the cells were incubated for another 24 h. 50 μL of supernatants from transfected cells were incubated with 30 μL of luciferase substrate (rep-qlc2, QUANTI-Luc^TM^, InvivoGen), and the luciferase activity was measured using a spectrophotometer (Varioskan lux, Thermo Fisher Scientific).

### Cell viability assay

The PoLixNano was prepared with a lipid composition of SM-102, DSPC, cholesterol, and DMG-PEG2000 (molar ratio = 45:20:33.5:1.5, N/P ratio of 8) and 5 mg/mL T904. The 16HBE cells (3.5×10^4^ cells/well) or DC 2.4 cells (2×10^4^ cells/well) were seeded in 96-well plates and incubated with PoLixNano or LNP counterpart (400 ng of IVT mRNA per well) at 37 ℃ for 6 h. The cytotoxicity of PoLixNano or LNP counterpart was investigated using the cell counting kit-8 (CCK8, HY-K0301, MedChemExpress).

### Preparation and analysis of bone marrow-derived dendritic cells (BMDCs)

Bone marrow cells were isolated from the femurs of female BALB/c mice by rinsing with RPMI 1640 medium (C11875500BT, Gibco), lysing with the red blood cell lysis buffer (C3702, Beyotime) and centrifuging (1200 rpm, 5 min). The cells were cultured in RPMI 1640 complete medium supplemented with 10% FBS, 1% penicillin/streptomycin, 20 ng/mL granulocyte-macrophage colony-stimulating factor (GM-CSF) (315-02, PEPROTECH), and 10 ng/mL Interleukin 4 (IL-4) (214-14, PEPROTECH). The medium was half-replaced every 2 days. On day 7, adherent cells were collected and transferred into a 24-well plate at a density of 5×10^5^ cells/well for investigating uptake efficiency and internalization mechanism using approaches as above-described. For the maturation assay, BMDCs were seeded in 24-well plates at a density of 5×10^5^ cells/well. PoLixNano or LNP formulations containing 3 μg of RBD-mRNA were added to each well and then incubated for 24 h. After the incubation, BMDCs were harvested and washed twice with PBS. Subsequently, anti-mouse-CD11C-FITC (11-0114-82, Invitrogen, 1:100), anti-mouse-CD40-PE (12-0401-82, Invitrogen, 1:100), anti-mouse-CD86-APC (17-0862-81, Invitrogen, 1:100), anti-mouse-CD80-Percp (105026, Bio-legend, 1:100), and anti-mouse-MHCII-eFluor^TM^ 450 (48-5321-82, Invitrogen, 1:100), were used to stain the cells at 4 ℃ for 30 min. After washing twice with PBS, flow cytometry (BD FACSCanto^TM^ II, BD Biosciences) was used to assess BMDC maturation.

### Organoid cultures

#### Organoid culture

Gastric cancer (GC) organoids were derived from peripheral blood and tissue samples of patients with GC receiving surgical resection at the Second Affiliated Hospital of Army Medical University. The study was approved by the Medical Ethics Committee of the Second Affiliated Hospital of Army Medical University (Approved Protocol ID: 2022-096-01). All experiments were performed in compliance with the Declaration of Helsinki. The expansion conditions of GC organoids were conducted as follows: organoids were suspended with cell recovery solution (354253, BD Falcon), incubated for 30 min on ice in a constant shaking incubator (100 rpm), and collected by centrifugation (200 g, 5 min). TryPLE Express (12605-028, Gibco) was used digested organoids into units. Then units were seeded into 6-well plates at a cell density of 2,000 cells coated by 20 µL Matrigel (356231, Corning), and maintained in GC organoid basal medium (K2179-GC, bioGenous) in a 5% CO_2_ environment at 37 °C. Organoids were regenerated on day 7 to day 10 after medium change every other day. Experiments were performed between passage 7 to passage10.

### Transfection of IVT mRNA in organoids

GC organoids cultured in 6-well plates were suspended in organoid cryopreservation medium (E238023, bioGenous) and incubated for 30 min on ice in a constant shaking incubator (100 rpm). The organoids were centrifuged (200 g, 5 min) and added into 24-well plates (pre-treated by anti-adherence rinsing solution (E238002, bioGenous)) at a density of 200 organoids (diluted in 100 µL organoid basal medium) per well. The PoLixNano or LNP was mixed with organoid basal medium based on 1:200 (w/v) ratio of IVT mRNA to organoid basal medium. The mixture solution containing 1 µg IVT mRNA was added per well in replicates of 3. After 12 h incubation in a 5% CO_2_ environment at 37 °C, organoids were re-coated with Matrigel. After another 24 h of incubation, the eGFP expression was assayed using confocal microscopy (FV3000, Olympus, Japan) and flow cytometry (BD FACSCanto^TM^ II, BD Biosciences), and the luciferase activity was evaluated by a microplate reader (Varioskan lux, Thermo Fisher Scientific).

### Animal studies

All in vivo studies were approved by the Laboratory Animal Welfare and Ethics Committee of Third Military Medical University (AMUWEC20210929) and were performed in accordance with the institutional and national policies and guidelines for the use of laboratory animals. Specific pathogen-free (SPF) female BALB/c mice (6–8 weeks old), C57/B6 mice (6–8 weeks old), New Zealand white rabbits (2-2.5 kg), Sprague-Dawley (SD) rats (8-10 weeks old) and Syrian golden hamsters (7–8 weeks old) were purchased from the Vital River Laboratory Animal Technology Co., Ltd. (Beijing, China). B6CF mice were ordered from Cyagen Biosciences (Guangzhou, China). All animals were kept under SPF conditions undergoing a 12 h/12 h light/dark cycle with free access to food and water. Animals were conceded an adaption time of at least 7 days and randomly divided into groups before experiments.

### In vivo bioluminescence

Unless otherwise specified, PoLixNano or LNP was administered to mice, Syrian golden hamsters and SD rats via nebulized dose per animal of 100 μg, 200 μg and 100 μg Fluc-mRNA respectively. At indicated time point post-administration, the animals were anesthetized with isoflurane and subsequently injected intraperitoneally (i.p.) with D-luciferin potassium salt (122799, PerkinElmer) dissolved in PBS (150 mg/kg body weight). Bioluminescence imaging was performed using an In Vivo Imaging System (IVIS) (Lumina Series III, PerkinElmer). Unlike mice, Syrian golden hamsters and SD rats, rabbits were anesthetized with urethane before nebulization. PoLixNano or LNP was administered to rabbits at a dose of 600 μg NanoLuc-mRNA per rabbit. At 12 h post-administration, the lungs of rabbits were isolated and the luciferase activity was detected by Nano-Glo^TM^ Luciferase Assay (N1110, Promega).

### In vivo biodistribution and cellular uptake of formulations

DiD-PoLixNano or DiD-LNP was administered to mice via nebulization, major organs (brain, lung, heart, liver, spleen and kidney) were isolated 6 h post-administration for fluorescence imaging using the IVIS. Meanwhile, the isolated lungs were minced into small pieces, and digested in complete 1640 medium containing 150 U/mL Collagenase Ⅴ (C8170, Solarbio) and 150 μg/mL of DNase I (D8071, Solarbio) at 37 ℃ for 60 min with gently shaking. The resulting cell suspension was filtered through 70 μm cell strainer (352350, Corning) and lysed with red blood cell lysis buffer (C3702-500mL, Beyotime). Following centrifugation (300 g, 5 min), the cells were collected, washed with 1×PBS, and re-suspended in a cell staining buffer (554656, BD). Subsequently, the cells were incubated with anti-mouse-CD11C-PE-Cyanine7 (25-0114-82, Invitrogen, 1:100), anti-mouse-EpCAM-BV421 (118225, Bio-Legend, 1:100), anti-mouse-CD-Podoplanin-PE (127407, Bio-Legend, 1:100), anti-mouse-CD-F4/80-PerCp-Cy5.5 (123127, BioLegend, 1:100), anti-mouse-CD-Ly6G-APC-eFluor™ 780 (47-9668-82, Invitrogen, 1:100) and anti-mouse-CD11B-FITC (11-0112-82, Invitrogen, 1:100) for 30 min at 4 ℃ to identify DC cells, epithelial cells, type 1 pneumocytes, macrophages and neutrophils. The DiD labelled positive cells in various pulmonary cell types were analyzed by the flow cytometry (BD LSRFortessa^TM^, BD).

### Immunohistochemical assay

DiD-PoLixNano or RBD-mRNA@PoLixNano was administered to mice via nebulized dose per mouse of 100 μg RBD-mRNA respectively. At 6 h post-administration, the lungs were immersed in 4 % paraformaldehyde for 24 h, embedded in paraffin and sectioned at a thickness of 5 μm. The lungs were deparaffinized, rehydrated, and incubated with 3% bovine serum albumin (BSA) for 30 min to block non-specific binding sites. The lung sections were incubated with primary antibodies anti-CD11C (GB11059, Servicebio, 1:500) or anti-EpCAM (ab221552, Abcam, 1:200) at 4 °C overnight, followed by incubation with specific secondary antibodies (GB25303, Servicebio, 1:400) for 1 h at RT. The lung sections from RBD-mRNA@PoLixNano group were additionally incubated with primary antibodies anti-SARS-CoV-2 Spike (40592-MM135, Sino Biological, 1:300) overnight, followed by incubation with Cy3-conjugated goat-anti-mice IgG (GB21301, Servicebio, 1:300) secondary antibody. The lung sections were counterstained with DAPI to visualize cell nuclei. The image information was collected using a Pannoramic MIDI system (3DHISTECH, Budapest) and FV1200 confocal microscopy (Nikon Eclipse C1).

### In vivo eGFP-mRNA expression

PoLixNano or LNP was administered to mice via nebulization at a dose of 100 μg eGFP-mRNA per mouse. Lungs were excised 6 h post-nebulization, dissociated, and treated with red blood cell lysis solution to obtain single-cell suspensions. Following centrifugation (300 g, 5 min), cells were re-suspended in a cell staining buffer and incubated with anti-mouse-CD11C-APC (17-0114-82, Invitrogen, 1:100), anti-mouse-EpCAM-BV421 (118225, Bio-Legend, 1:100), anti-mouse-CD-Podoplanin-PE/Cyanine7 (127417, BioLegend, 1:100), anti-mouse-CD-F4/80-Brilliant Violet 650^TM^ (123149, BioLegend, 1:100), anti-mouse-CD-Ly6G-APC-eFluor™ 780 (47-9668-82, Invitrogen, 1:100) and anti-mouse-CD11B-PE (101207, Bio-Legend, 1:100) for 30 min at 4 ℃ to identify DC cells, epithelial cells, type 1 pneumocytes, macrophages and neutrophils, respectively. The quantification of eGFP positive cells was performed by flow cytometry (BD LSRFortessa^TM^, BD).

### Histopathology analysis

24 h post-administration of nebulized PoLixNano or LNP, major organs of mice including lungs, brains, intestines, livers, spleens and hearts were collected and fixed overnight in 4 % paraformaldehyde overnight at RT. The fixed organs were dehydrated, embedded in paraffin and sliced into 5 μm thickness. The lung sections were stained with hematoxylin and eosin (H&E) and visualized using light microscope (Eclipse Ci-L, Nikon).

### Cytokine detection in lung homogenates, BALF and serum

At indicated time points post-administration of PoLixNano or LNP, the levels of interleukin 4 (IL-4), interleukin-6 (IL-6), interleukin-17 (IL-17) and tumor necrosis factor α (TNF-α) in lung homogenates, bronchoalveolar lavage fluid (BALF) and serum were quantified using Enzyme-Linked Immunosorbent Assay (ELISA) kits (Dakewe Biotech, China). Briefly, serially diluted samples were added to the corresponding ELISA plates (IL-4 (61210402, Dakewe Biotech), IL-6 (1210602, Dakewe Biotech), IL-17 (1211702, Dakewe Biotech) and TNF-α (1217203, Dakewe Biotech)) and incubated for 90 min at 37 ℃. Subsequently, horseradish peroxidase (HRP)-linked streptavidin was added and incubated for 30 min at 37 ℃. Tetramethylbenzidine (TMB, ab171522, Abcam) substrate was added and incubated for 5 min at 37 ℃. The reaction was stopped with stop solution (P0215, Beyotime). The absorbance at 450 nm was measured using a microplate reader (Varioskan lux, Thermo Fisher Scientific).

### Mouse serum chemistry metrics and anti-PoLixNano ELISA

Mice were dosed with PoLixNano on day 0 and day 21. Serum samples were collected at indicated time points (day 1, day 3, day 7, day 14 and day 28) and serum chemistry metrics including alanine aminotransferase (ALT), aspartate aminotransferase (AST), total protein, urea nitrogen, alkaline phosphatase (ALP), triglycerides (TG), phosphorus, creatinine were analyzed by Servicebio company (Wuhan, China).

For anti-PoLixNano antibody ELISA, a 50 μL aliquot of serum were isolated at indicated time points (day 14, day 28 and day 42). High-binding 96-well plates (9018, Corning) were coated with 20 μg/well copolymer, lipids and PoLixNano, incubated overnight at 4 ℃ and washed with PBS containing 0.05 % Tween-20 (PBST) for 3 times. The plates were blocked with 1% bovine serum albumin (BSA) for 2 h at 37 ℃. The serum samples diluted in BSA at a ratio of 1:100 (v/v) were added and incubated for 1 h at 37 ℃. The goat anti-mouse IgG (ab205719, Abcam, 1:10000) was added and incubated for another 1 h at 37 ℃. The signal was developed using TMB substrate, the reaction was stopped using stop solution and the absorbance at 450 nm was measured using a plate reader (Varioskan lux, ThermoFisher).

### Immunological analysis of RBD-mRNA vaccines

#### Immunization scheme

BALB/c mice were immunized on day 0 and boosted on day 21 with 100 μg/mouse RBD-mRNA encapsulated in PoLixNano or LNP (LNP-neb) via nebulization, or 5 μg/mouse RBD-mRNA loaded within LNP (prepared as same LNP as the approved COVID-19 mRNA vaccine mRNA-1273^6^) via intramuscular injection. Serum samples were collected on day 14, day 21, day 28, day 42, day 56, and day 380, respectively. BALF was collected on day 42 or day 380 by lavaging the lungs with 1000 μL of 1×PBS via a polyethylene tube cannulated into the trachea (downward flushing). Similarly, nasal lavage fluid (NLF) was similarly obtained by lavaging with 1000 μL of 1×PBS through a polyethylene tube (upward flushing).

#### Determination of RBD-specific antibody titers using enzyme-linked immunosorbent assay (ELISA)

The antigen-specific IgG or IgA antibodies in the serum, BALF and NLF were analyzed by ELISA. Briefly, high-binding 96-well plates were coated with 20 ng/well of RBD protein overnight at 4°C in coating buffer (421701, BioLegend), and the plates were blocked with 1% BSA in PBST for 1 h at 37°C. Serially diluted samples were added to the 96-well plates and incubated for 1 h at 37 °C. After incubation, the plates were washed five times with PBS containing 0.05% Tween 20 (PBST), followed by the addition of 100 μL of goat anti-mouse IgG (ab205719, Abcam, 1:10000) or goat anti-mouse IgA (ab97235, Abcam, 1:10000) secondary antibody at 37°C for 40 min incubation. Subsequently, 50 μL TMB substrate was added to each well and incubated for 10 min at 37°C. The absorbance at 450 nm (OD_450_) was measured using a microplate reader (Varioskan lux, Thermo Fisher Scientific). The endpoint titer was defined as the highest reciprocal dilution of the detected sample that gave an absorbance greater than 2.1-fold of the OD_450_ value from the PBS group.

#### Pseudovirus neutralization assay

The neutralizing ability of serum and BALF was assessed using a pseudovirus neutralization assay. Angiotensin-Converting Enzyme 2 (ACE2)-expressing HEK-293T cells (hACE2-293T) (41107ES03, Yeasen) were cultured in DMEM (11965118, Thermo Fisher Scientific) supplemented with 10% of FBS and 1% of penicillin/streptomycin. hACE2-293T cells were seeded in 96-well plates at a density of 2×10^4^ cells/well. Serum samples were serially diluted two-fold starting at a 1:800 in Opti-MEM, and BALF samples were diluted two-fold starting at a 1:32 in Opti-MEM. 100 μL of diluted serum or BALF sample were mixed with 10 μL of SARS-CoV-2 Omicron (B.1.1.529) pseudovirus encoding firefly luciferase (11991ES70, Yeasen) and incubated for 1 h at 37 ℃. The mixtures were subsequently added to the pre-seeded hACE2-293T cells and incubated for 48 h. After incubation, cells were lysed and luciferase activity was measured using luciferase assay reagent (RG006, Beyotime Biotechnology). The bioluminescence was recorded on a microplate reader (GloMax, Promega). The pseudovirus neutralizing antibody titers at 50% inhibition (pVNT_50_) were calculated as the serum dilution at which relative light units (RLU) were reduced by 50% compared to RLU in virus control wells after subtraction of background RLU in cell control wells.

#### Enzyme-linked immunosorbent assay (ELISpot)

The cellular immune responses in the lung and spleen of vaccinated mice were analyzed by mouse IFN-γ ELISpot kit (CT317-PR2, U-CyTech), mouse IL-4 ELISpot kit (CT319-PR2, U-CyTech) and IL-17 ELISopt PLUS plates (3521-4HPW-2, MABTECH). Briefly, on day 42 post-immunization, lungs were collected from immunized mice and digested with digestion buffer composed of RPMI 1640 medium supplemented with 150 U/mL Collagenase Ⅴ and 150 μg/mL of DNase I at 37 ℃ for 1.5 h with gently shaking at 220 rpm, and filtered through a 70 μm cells strainer. Pulmonary lymphocytes were harvested using density gradient centrifugation using Percoll (17-0891-09, GE Healthcare). Spleens were filtered using a 70 μm cell strainer in complete RPMI 1640, centrifuged (1200 rpm, 5 min), and lysed in red-cell lysis solution (C3702-500mL, Beyotime) for 5 min on ice, followed by washing with PBS to obtain a clear single-cell suspension of splenocytes. The pre-coated plates containing anti-mouse IFN-γ, IL-4 and IL-17 antibody were incubated with RPMI 1640 medium with 10% FBS at RT for 30 min. Subsequently, 1×10^6^ splenocytes or 1×10^5^ pulmonary lymphocytes were plated and stimulated with 5 μg/well of RBD peptide pool (Supplementary Table 3). The cells were incubated for 36 h at 37 ℃. Following incubation, the plates were washed four times with PBST and incubated with biotinylated anti-IFN-γ, anti-IL-4 antibody or anti-IL-17 antibody and streptavidin-HRP, all diluted in PBS containing 0.5% FBS at RT for 40 min. After adding the substrate solution, the spots were visualized and counted using an Immuno-Spot CLT (Bio-Rad, Germany).

To investigate antibody-secreting cells (ASC), MultiScreen-HA filter 96-well plates (MAHAS4510, Merck) were pre-coated with 30 ng/well of SARS-CoV-2 RBD protein overnight at 4 °C. Plates were rinsed with PBST and blocked with culture medium (RPMI 1640, 10% FBS, 1% penicillin-streptomycin) for 4 h at 37 °C. On day 35 post-prime, single-cell suspensions of splenocytes, bone marrow cells (which were flushed from femurs and tibia into PBS and filtered through a 70 μm strainer) and mediastinal lymph-nodes cells were prepared from various vaccinated groups (i.e. PBS-treated, LNP-neb, LNP-i.m. and PoLixNano groups). The cells were added to the RBD protein-coated plates and incubated at 37 °C for 4 h. After washing, plates were incubated with anti-IgG (3825-2H, MABTECH) or anti-IgA (3865-2H, MABTECH), followed by the incubation with streptavidin-conjugated horseradish peroxidase, each for 1 h at RT. After five washes with PBST, TMB substrate was added for spot development. The reaction was stopped by rinsing with a generous amount of ultrapure water, and the plates were dried overnight. Spots were counted using an automated ELISpot reader (Immuno-Spot-CLT, Bio-Rad).

#### Flow cytometry analysis

To investigate intracellular cytokine expression, pulmonary lymphocytes from immunized mice were isolated on day 42 post-prime and plated into 96-well plates at a density of 1×10^6^ cells per well. Cells were incubated for 6 h at 37 °C in the presence of eBioscience™ Protein-Transport-Inhibitor-Cocktail (00-4980-93, Thermo Fisher Scientific, 1:500), followed by stimulation with the RBD peptide pool (5 μg/well) at 37 °C for an additional 24 h. Subsequently, cells were stained with anti-mouse-CD3e-FITC (553061, BD Biosciences, 1:200), anti-mouse-CD4-PercpCy 5.5 (550954, BD Biosciences, 1:200), and anti-mouse-CD8α-PE-Cy7 (552877, BD Biosciences, 1:200) antibodies in staining buffer at 4 °C for 30 min. Cells were fixed with the Cytofix/Cytoperm Fixation/Permeabilization Kit (554714, BD Biosciences), followed by the incubation with anti-mouse-IFN-γ-eFluor™ 450 (48-7311-82, Invitrogen, 1:50), anti-mouse-IL-4-APC (17-7041-82, Invitrogen, 1:50) or anti-mouse-IL-17-eFluor™ 450 (48-7177-82, Invitrogen, 1:50) antibodies at 4 °C for an additional 30 min.

To detect the effector memory T Cells (T_EM_), central memory T cells (T_CM_) and tissue-resident memory T cells (T_RM_), pulmonary lymphocytes or splenocytes were prepared from immunized mice on day 42 or day 380 post-immunization as previously described. Subsequently, cell samples were stained with the following antibodies in cell staining buffer for 40 min at 4 °C: anti-mouse-CD3e-FITC (553061, BD Biosciences, 1:200), anti-mouse-CD4-PercpCy5.5 (550954, BD Biosciences, 1:200), anti-mouse-CD8α-PE-Cy7(552877, BD Biosciences, 1:200), anti-mouse-CD62L-PE (553151, BD Biosciences, 1:100), anti-mouse-CD44-APC (559250, BD Biosciences, 1:100), anti-mouse-CD69-BV421 (562920, BD Biosciences, 1:80), anti-mouse-CD103-BV510 (563087, BD Biosciences, 1:80).

For germinal center B cells (GCB) and T follicular helper cells (Tfh) analysis, mice were euthanized by exsanguination on day 28 post-immunization, and the MLNs were collected, homogenized and filtered through a 70 μm cell strainer. Cell samples were incubated with cell staining buffer containing anti-mouse-CD3e-eFluor^TM^ 506 (69-0032-82, Invitrogen, 1:200), anti-mouse-CD185(CXCR5)-APC (17-7185-82, Invitrogen, 1:200), anti-mouse-CD279 (PD-1)-PE (12-9985-82, Invitrogen, 1:100), anti-mouse-CD4-PercpCy5.5 (45-0042-82, Invitrogen, 1:100) and anti-mouse-CD44-FITC (11-0441-82, Invitrogen, 1:100) at 4 ℃ for 30 min. For GCB cell analysis, cell suspensions were incubated with a mixture of anti-mosue-CD95-PE (12-0951-81, Invitrogen, 1:100), anti-mouse-Gr1-FITC (11-5931-82, Invitrogen, 1:100), anti-mouse-B220-V450(48-0452-82, Invitrogen, 1:100), and anti-mouse-CD19-APC (17-0913-82, Invitrogen, 1:100) at 4 °C for 40 min.

To quantify the alveolar macrophages (AMs) and interstitial macrophages (IMs), single-cell suspensions were prepared from BALF of immunized mice collected on day 42 post-immunization, stained with the biotin-anti-mouse-Ly-6C (128003, Bio-Legend, 1:50) for 40 min at 4 °C, and subsequently incubated with the following antibodies in cell staining buffer containing anti-mouse-CD11B-PE (12-0112-8, Invitrogen, 1:100), anti-mosue-Ly6G-APC-eFluor™780 (47-9668-82, Invitrogen, 1:100), anti-mosue-CD11C-FITC (11-0114-82, Invitrogen, 1:100), anti-mouse-MHC II-eFluor™450 (48-5321-82, Invitrogen, 1:100), anti-mouse-CD24-SuperBright 600 (63-0242-82, Invitrogen, 1:100), anti-mouse-CD64-Percp-eFluor^TM^710 (46-0641-82, Invitrogen, 1:100), anti-CD170 (SiglecF)-PE-Cy7 (25-1702-82, Invitrogen, 1:100), and APC streptavidin (405307, Bio-Legend, 1:100) for 30 min at 4 °C.

All stained cell samples were rinsed five times with PBS before analyzing on a BD LSRFortessa^TM^ (BD Biosciences). Data was analyzed and illustrated by FlowJo software V10.8.1.

### RNA sequencing (RNA-seq)

RNA sequencing analysis was performed as previously described ^7^. Briefly, BALB/c mice were immunized with PoLixNano, LNP-neb, LNP-i.m. or PBS as described above. Mice were euthanized 24 h or 28 days post-prime. Lungs were extracted and minced with scissors. Total RNA extraction, purification, reverse transcription, and library construction, were prepared by Shanghai Majorbio Biotechnology Co., Ltd. (Shanghai). RNA libraries were constructed using the TruSeq™ RNA Sample Preparation Kit (Illumina, San Diego). High-throughput sequencing was conducted on the Illumina HiSeq XTEN/NovaSeq 6000 sequencer with a read length of 2×150 bp. To ensure data quality, SeqPrep (https://github.com/jstjohn/SeqPrep) and Sickle (https://github.com/najoshi/Sickle) were employed for trimming and quality control. Clean reads were mapped to mouse genome using HISAT2 software ^8^.

The levels of differential expression genes (DEGs) were calculated using transcripts per million (TPM) reads, and gene abundances were quantified using expectation-maximization (RSEM). DEGs were identified based on |log2FC| ≧ 1 and FDR≤ 0.05 (DESeq2)^9^, or FDR ≤0.001 (DEGseq)^10^. The Benjamini-Hochberg correction method was applied, and DEGs with an adjusted *P*-value < 0.05 were deemed statistically significant. Gene Ontology (GO) analysis was conducted using Goatools, and Kyoto Encyclopedia of Genes and Genomes (KEGG) pathways analysis was performed using KOBAS to identify substantially enriched DEGs^11^. Enrichment was determined using a Bonferroni-corrected *P-*value below 0.05.

### Metabolomics

Metabolome sequencing was conducted as previously described^12^. BALB/c mice were immunized with PoLixNano, LNP-neb, LNP-i.m., or PBS. The mice were euthanized 24 h or 28 days post-prime. Lungs were extracted using 400 μL 80% methanol. The extracts were homogenized using the frozen tissue grinder (Shanghai wanbo biotechnology co., LTD) at - 10 ℃, 50 Hz. After ultrasonic extraction for 30 min at 5 ℃, the samples were stored at -20 ℃ for 30 min. Subsequently, the samples were centrifuged at a rate of 13, 000 g for 15 min at 4° C, and the supernatant was stored in an injection vial for LC-MS/MS analysis. The chromatographic conditions were as follows: column, Waters ACQUITY HSS T3 C18 (100 mm × 2.1 mm, 1.8 μm); mobile phase, ultrapure water (with 0.1% formic acid in water:acetonitrile (95:5, v/v)) for phase A, and acetonitrile (with 0.1% formic acid in acetonitrile:isopropanol:water (47.5:47.5:5, v/v/v)) for phase B. The flow rate was maintained at 0.40 mL/min with a column temperature set at 40 ℃. An injection volume of 3 μL was utilized.

The raw data from the UHPLC-MS was processed using Progenesis QI software (Waters, Milford, USA), which involved baseline filtering, peak identification, integration, retention time correction, and peak alignment. The resulting data matrix, which included sample names, m/z values, retention times, and peak intensities, was exported for further analysis. Concurrently, metabolites were identified by searching relevant databases, primarily the HMDB (http://www.hmdb.ca/)^13^, Metlin (https://metlin.scripps.edu/), and the self-compiled Majorbio Database (MJDB) from Majorbio Biotechnology. The final data was normalized and analyzed using the Majorbio cloud platform (www.majorbio.com)^14^.

### Live SARS-CoV-2 virus challenge and virus transmission studies

#### Live SARS-CoV-2 virus challenge

Studies with SARS-CoV-2 ancestral strain C57MA14 and Omicron BA.5 were performed in biosafety level 3 laboratories. BALB/c mice were immunized as above mentioned. On day 45, the pre-immunized mice were challenged intranasally under anesthesia with either 10^5^ 50% tissue culture infective dose (TCID_50_) in 50 μL volume of a mouse-adapted lethal SARS-CoV-2 strain C57MA14 (NCBI GenBank number: OL913104.1, details can be found in: https://www.ncbi.nlm.nih.gov/nuccore/2167992552) or 10^4^ TCID_50_ in 50 μL volume of SARS-CoV-2 strain Omicron BA.5 (NCBI GenBank number: OP984772.1, details can be found in: https://www.ncbi.nlm.nih.gov/nuccore/OP984772.1). Body weight and mortality of the mice were monitored daily post infection. Three animals per group were euthanized at indicated time points in Fig 5a and Extended Fig. 10a. The nasal turbinates or lungs were collected for viral loads measurement and histopathological examination.

#### Vaccination and infection of hamsters

The Syrian Golden hamsters were immunized with a prime-boost vaccination strategy on day 0 and day 21. Each hamster received PoLixNano via nebulization at a dose of 200 μg RBD-mRNA, and received LNP (LNP-i.m.) via intramuscular injection at a dose of 10 μg RBD-mRNA. On day 35 post-vaccination, one naïve hamster (donor) was intranasally challenged with 10^4^ TCID_50_ in 50 μL volume of Omicron BA.5. On the second day post challenge, two vaccinated hamsters (recipients) (i.e. PBS-treated, LNP-i.m. and PoLixNano groups) were co-housing with a donor hamster for 48 h, sharing diet, bedding, and ventilation. The nasal rinsing samples were collected from recipients, and body weight was recorded daily for 7 days post co-housing. Hamsters were euthanized at the indicated time points to assess viral loads in the nasal turbinates and lung homogenates, and to perform histopathological and immunohistochemical analyses in the lungs.

#### Quantification of SARS-CoV-2 viral RNA

SARS-CoV-2 viral RNA in the nasal turbinates and lungs from challenged mice or hamsters was measured via quantitative reverse transcription PCR (RT-qPCR) as previously described^15^. Briefly, the tissue samples were homogenized with Tissuelyser-24 (Shanghai jingxin Industrial Development CO., LTD), total RNA was extracted and cDNA synthesis was performed. The RNA copies of SARS-CoV-2 was quantified by RT-qPCR targeting the S gene using the One Step PrimeScript RT-PCR Kit (RR057B, Takara) with specific primers and probes: CoV-F3 (5’-TCCTGGTGATTCTTCTTCAGGT-30). CoV-R3 (5’-TCTGAGAGAGGGTCAAGTGC-30), and CoV-P3 (5’-FAM-AGCTGCAGCACCAGCTGTCCA-BHQ1-30).

#### Histopathological and immunohistochemistry studies

For pathological analysis, lungs were collected, fixed in 4% paraformaldehyde for 72 h, dehydrated and embedded in paraffin. The wax block of lung tissues was cut into 5-µm-thick sections for H&E staining and analysis as described above. Immunohistochemical (IHC) staining for SARS-CoV-2 nucleocapsid protein (N) was performed to assess viral antigen expression and distribution in the lungs. SARS-CoV/SARS-CoV-2 Nucleocapsid (40143-T62, Sino Biological Inc, 1:200) and Peroxidase AffiniPure goat anti-rabbit IgG (H+L) (111-035-003, Jackson ImmunoResearch, 1:500) were used as the primary and secondary antibody of IHC. For hamsters IHC study, Rabbit Two-Step Kit (PV-6001, Beijing Zhongshan Golden Bridge Biotechnology Co., Ltd) was applied as the secondary antibody. The DAB Horseradish Peroxidase Color Development Kit (ZLI-9019, Beijing Zhongshan Golden Bridge Biotechnology Co., Ltd) was used for the chromogenic reaction. The tissue staining was visualized using a microscope (Eclipse Ci-L, Nikon).

#### Cytokine and chemokine measurements

Cytokines levels within lung homogenates from challenged mice were detected by high-throughput liquid-phase protein microarray platform. Briefly, samples were collected at the indicated time points and analyzed for cytokines and chemokines by Laizee Biotech (Beijing, China) using mouse Th1/Th2 Cytokine Panel (11 plex) kit which includes IFN-γ, IL-12p70, IL-13, IL-1β, IL-2, IL-4, IL-5, IL-6, TNF-α, GM-CSF and IL-18. Data were obtained using the Luminex 200 system (Thermo Fisher Scientific).

### Metastatic lung cancer model

The lung metastatic cancer model were established in C57BL/6 mice which received an intravenous injection of B16F10-Luc cells (1 ×10^6^ cells/mouse) suspended in 200 μL PBS on day 0. PoLixNano or LNP containing 100 μg IL-12 mRNA/mouse was administrated to mice via nebulization on day 6, day 9 and day 12. On day 21, the lungs from vaccinated mice were excised, weighed to determine lung mass, and examined for gross lesions. Tumors progression was monitored and quantified using the IVIS following intraperitoneal (i.p.) injection of the D-luciferin potassium salt.

### RT–qPCR analysis of tyrosinase-related protein 1 (Tyrp1) expression

Total RNA from the lung of mice was extracted using the MicroElute Total RNA Kit (R6731-01, OMEGA), and 1 μg RNA was reverse-transcribed into cDNA using the PrimeScript RT Reagent Kit (RR037B, Takara). RT-qPCR analysis was conducted using Hieff SYBR Green Master Mix Kit (11201ES08, YEASEN). The primer sequences of *Tyrp1* were as following: F—5′-CAGGTTGTCTGGAGCAGTAAC-3′, R—5′GGCTGAGGAGATACAATGCTG-3′. The primer sequences of internal control gene *GAPDH* were as following: F—5′-CAAAATGGTGAAGGTCGGTGTG-3′, R—5′-TGATGTTAGTGGGGTCTCGCTC-3′.

### Immunofluorescence of CFTR

PoLixNano or LNP was administered to mice via nebulization at a dose of 100 μg CFTR-mRNA or Fluc-mRNA per mouse with three repeated dosing. At indicated time points, B6CF mice were anesthetized with isoflurane and subsequently injected i.p. with D-luciferin potassium salt for bioluminescence imaging using the IVIS. At 72 h post first dosing, B6CF mice were humanely euthanized, and lungs were collected for further investigations. The *CFTR* expression within the lung sections was detected as previously described^16^. Briefly, lung sections (5 μm) were blocked with 10% donkey serum and subsequently incubated with the mixture of primary antibodies against CFTR (“570”, the University of North Carolina, 1:100) and ZO1 (ab99462, Abcam, 1:200). Secondary antibodies conjugated with Cy3 (donkey anti-mouse, 1:500) and Alexa 488 (donkey anti-goat, Abcam, 1:200) were applied.

### Western blot for CFTR detection

The *CFTR* expression within the lungs of B6CF mice treated by PoLixNano or LNP (LNP-neb) was detected via Western blot as previously described^16^. Briefly, the lungs were harvested, digested and lysed 72 h post-dosing. Total protein concentration was measured using BCA™ protein assay kit (23227, Thermo Fisher Scientific). Lung lysates were separated on SDS polyacrylamide gel (6%) overnight at 4℃, and the protein was transferred onto a nitrocellulose membrane (Protran supported 0.45 µm, Amersham) using a transfer buffer optimized for proteins of high molecular weight (125 mmol Tris, 950 mmol glycine and 0.02% SDS). A combination of CFTR primary antibodies (‘217’, ‘570’, ‘596’ and ‘660’, the University of North Carolina, 1: 400) was applied to detect the CFTR. The membrane was further incubated with polyclonal goat anti-mouse immunoglobulin secondary antibody (ab97040, Abcam, 1:500) and developed with a Pierce Femto Fast Western Blot Kit and Pierce Pico Fast Western Blot Kit (35065, Thermo Fisher Scientific). Signals were detected in consecutive series with chemiluminescence (Chemi-Bis, Bio-Rad).

### Statistical Analysis

Unless otherwise specified, data for all bar charts were plotted using means with error bars corresponding to standard error of the mean (SEM). Statistical analysis was performed using Prism 8 (GraphPad Software). Dual comparisons were made using Welch’s t-test, and comparisons between multiple conditions were analyzed using analysis of variance (ANOVA) followed by the appropriate post-hoc tests. All of the tests were two tailed. Differences were considered statistically significant when *P*<0.05.

## Acknowledgements

This work was supported by the National Natural Science Foundation of China (NSFC, Grant No. 82173764 and 32370993), the major project of Study on Pathogenesis and Epidemic Prevention Technology System (2021YFC2302500) by the Ministry of Science and Technology of China, NHMRC Emerging Leadership Investigator Grant (GNT2017974), and the Bundesministerium für Bildung und Forschung (BMBF, Grant No. 03VP10060, Zell-Trans). We appreciate Y. Qiu (Second Affiliated Hospital of Army Medical University) for performing organoids related investigations. We thank Dr. X. Zhu (Shanghai Vitalgen BioPharma Co., Ltd) and Prof. J. Zhang (Third Military Medical University) for helpful discussion. We extend our gratitude to X. Ding (Ludwig Maximilian University of Munich), Y. Zhang (Affiliated Hospital of North Sichuan Medical College, Sichuan, China), Y. Zhu (Southwest University, Chongqing, China), J. Yang (Chongqing University, Chongqing, China), C. Li (Fuzhou University, Fujian, China), C. Xu (MDPI, Wuhan, China), K. Wang, R. Xue, C. Liu, Y. Long and X. Xia (Third Military Medical University) for technical support and the design of schematic diagram. D. Zhang acknowledges the support of the China Scholarship Council (CSC) (202208500036).

## Author contributions

S.G. conceived and directed the project. S.G., J.R. and Q.Z. contributed experimental materials. D.Z., Q.X., Y.C., L.X., Y.L., L.P., Y.W., and S.H. performed experiments. S.G., D.Z., K. X. J.T., L.C. and Q.X. analyzed data. T.C. and C.L. was responsible for the bioinformatics and RNA sequencing data analysis. S.G., Q.Z., and D.S. supervised the research. S.G., D.Z., Q.X., K.X. and D.S. wrote the manuscripts with help and comments from all authors.

## Data availability

All data that support the findings of this study are provided within the paper and its Supplementary Information. Source data are provided with this paper.

## Extended Data Figures

**Extended Data Fig. 1.**
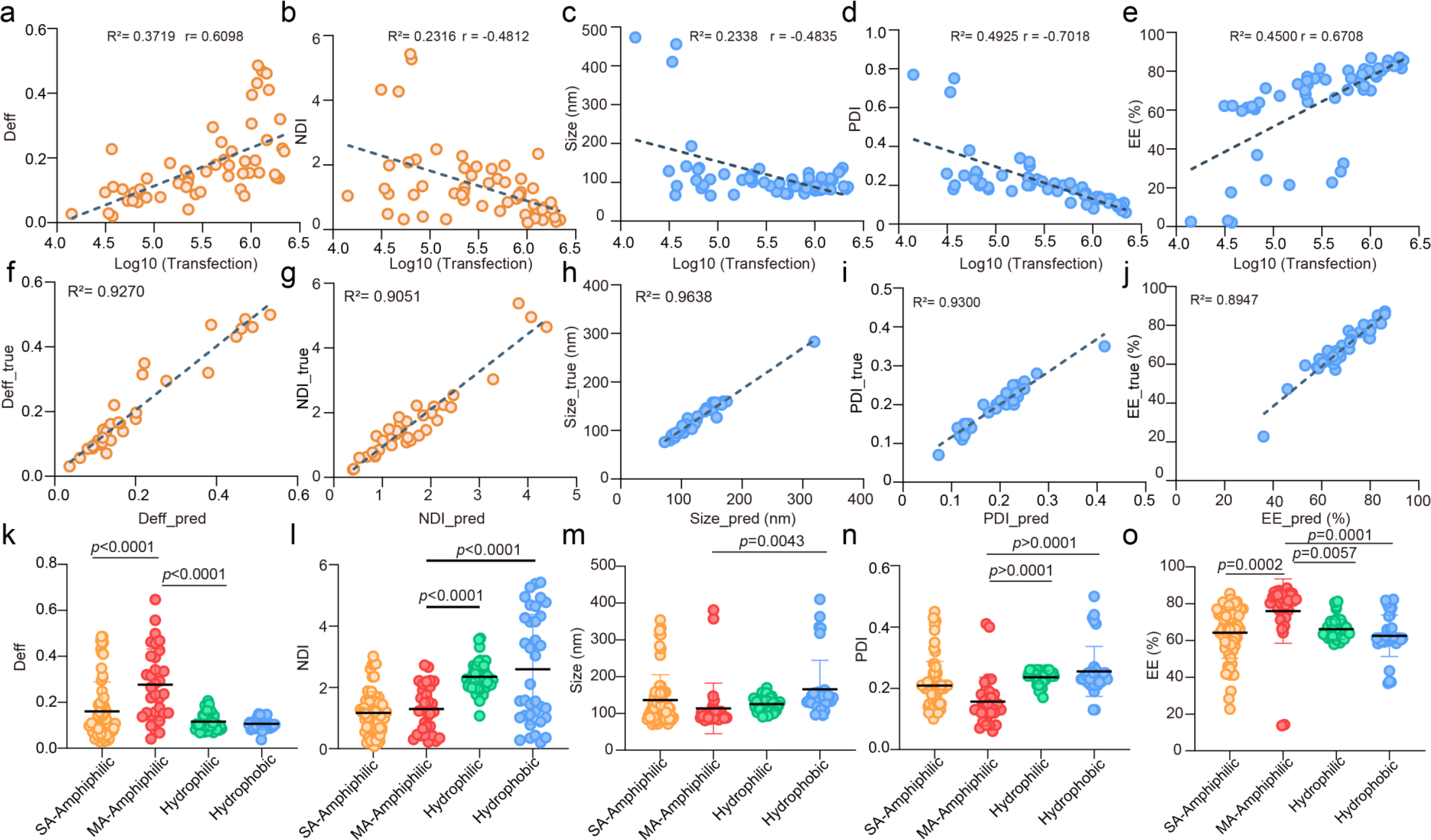
Machine learning model prediction and coefficients. Correlations of the diffusion coefficient (Deff) (**a**), nebulization shear force deformation index (NDI) (**b**), Size (**c**), polydispersity index (PDI) (**d**) and mRNA encapsulation efficiency (EE) (**e**) with Log10(transfection efficiency) (n=56). Predicted values vs experimental results of the Deff (**f**), NDI (**g**), Size (**h**), PDI (**i**) and EE (**j**) on test set (n=54). The Deff (**k**), NDI (**l**), Size (**m**), PDI (**n**) and EE (**o**) distributions of PoLixNano based on single-arm (SA) amphiphilic, multi-arm (MA) amphiphilic, hydrophilic, hydrophobic polymers. Statistical significance in **k-o** was calculated by one-way analysis of variance (ANOVA) followed by Tukey’s post-hoc test.

**Extended Data Fig. 2.**
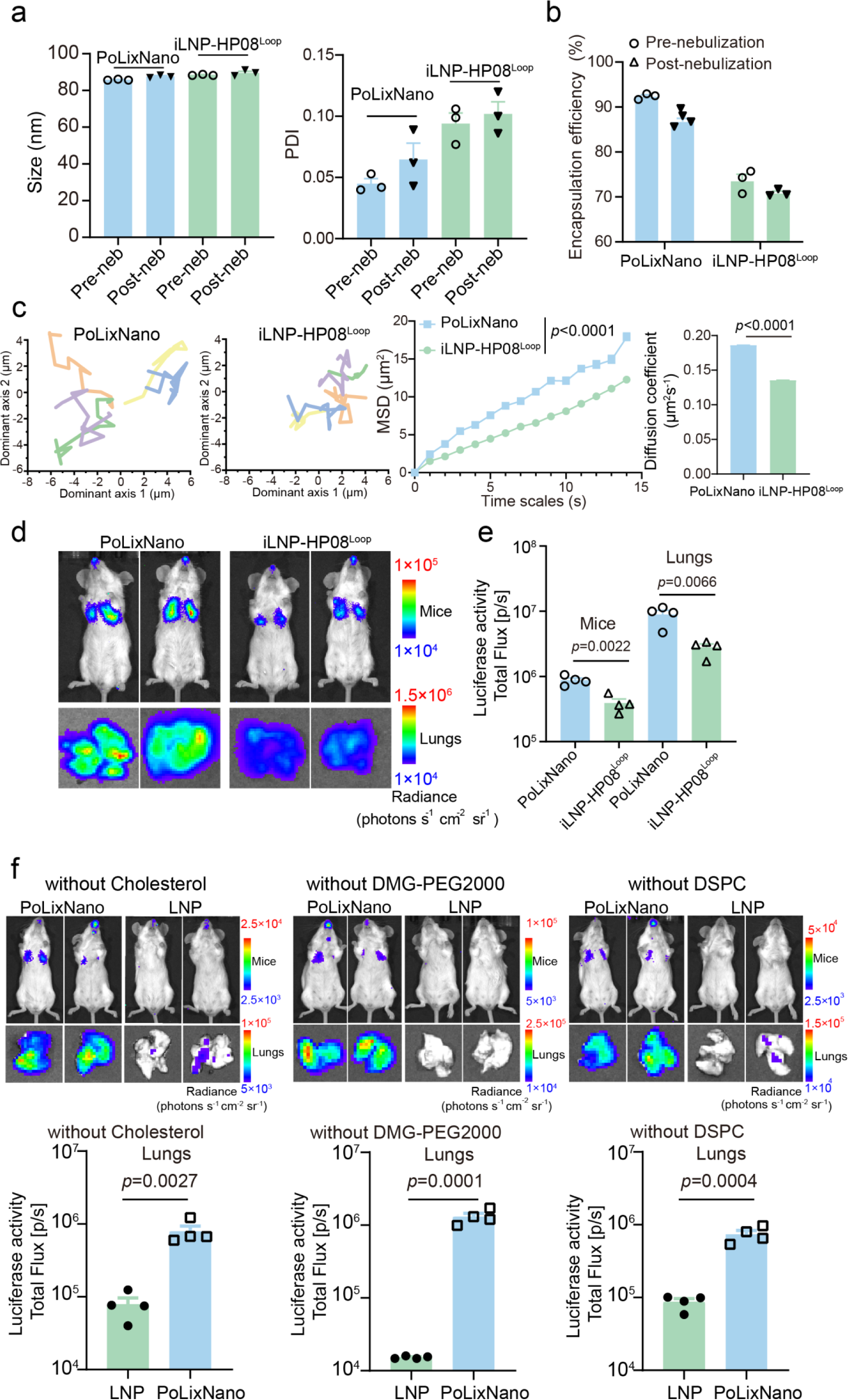
Comparison of PoLixNano vs iLNP-HP08^Loop^ and investigations on the transfection profiles of PoLixNano prepared with one lipid component removed. **a**, Measurements of particle size and polydispersity index (PDI) of PoLixNano and iLNP-HP08^Loop^ pre- and post-nebulization using DLS (n=3 independent replicates). **b**, The encapsulation efficiency (EE%) of PoLixNano and iLNP-HP08^Loop^ before and after nebulization (n=3 independent samples). **c**, Representative trajectories of nebulized PoLixNano and iLNP-HP08^Loop^ nanoparticles in a 3% (w/v) mucin solution over a 15-s period, with corresponding mean square displacements (MSD) and diffusion coefficient (Deff, μm²/s⁻¹) of the PoLixNano or iLNP-HP08^Loop^ nanoparticles (n=1148 independent nanoparticles). **d-e**, Representative bioluminescence images (**d**) of Fluc-mRNA expression 6 h post-nebulization of PoLixNano or iLNP-HP08^Loop^ in mice and excised lungs, with corresponding quantitative fluorescence intensities (**e**) (n=4 biologically independent animals). **f**, Representative bioluminescence images and the quantitative analysis of mouse lungs treated with PoLixNano or LNP without cholesterol, phospholipid or PEG-lipid at 6 h post-nebulization (n=4 biologically independent animals). Statistical significance was analyzed using two-tailed unpaired *t*-tests in **c**, **e** and **f**.

**Extended Data Fig. 3.**
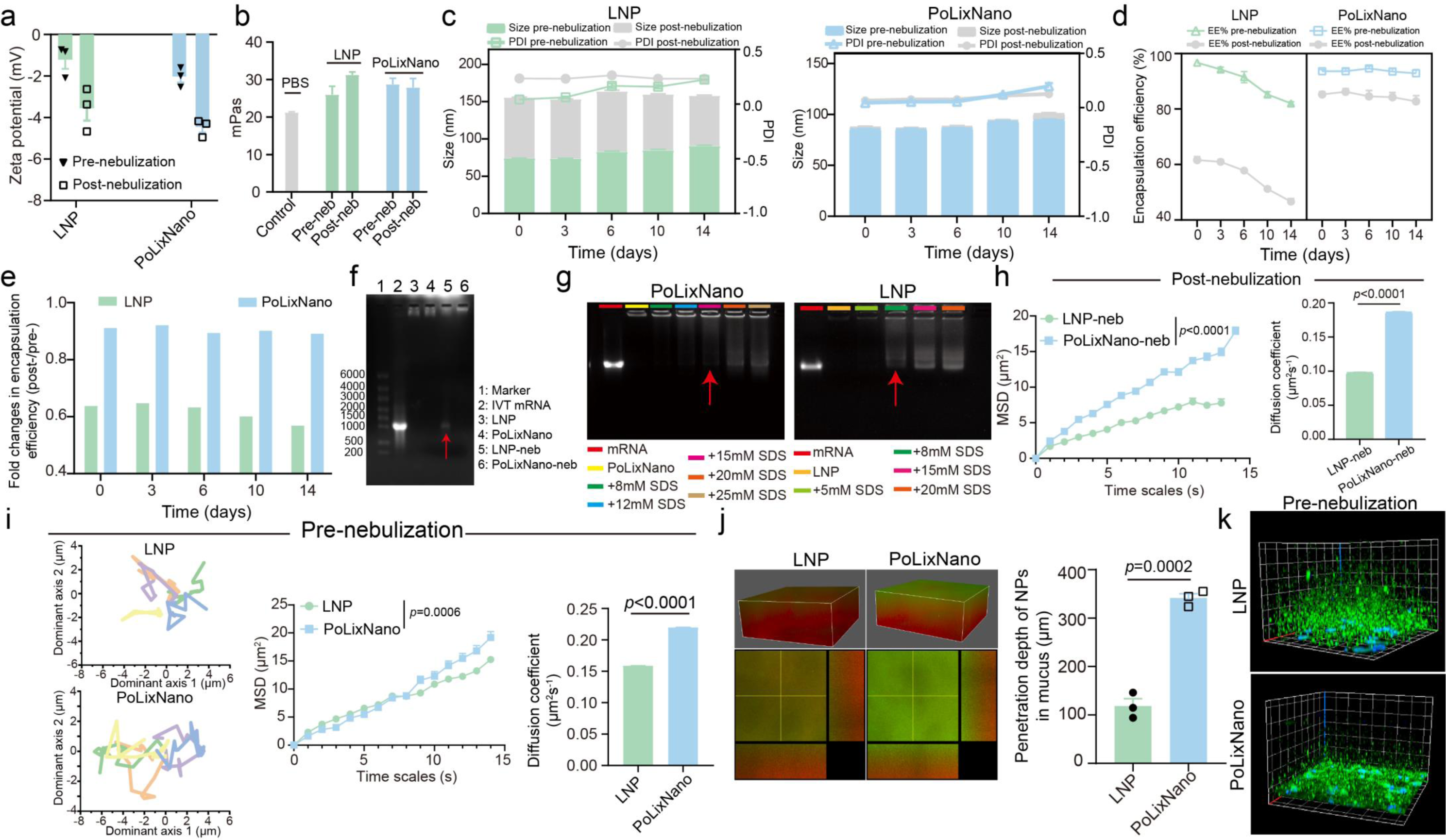
Supplementary physicochemical properties of the best-performing PoLixNano before and after nebulization. **a,** The zeta potential of PoLixNano or LNP before and after nebulization (n=3 independent samples). **b,** Viscosity assays (mPas) for PoLixNano and LNP pre- and post-nebulization (n=3 independent samples). **c-e,** Stability tests of PoLixNano or LNP stored at room temperature (RT) for indicated times. The particle size and PDI (**c**), EE % (**d**) and the fold changes in EE % (**e**) of LNP (left) and PoLixNano (right) were measured pre- and post-nebulization (n=3 independent samples). **f**, Agarose gel electrophoresis assay evaluating the leakage of mRNA from the indicated non-nebulized (lane 3-4) or nebulized (lane 5-6) formulations. Naked mRNA (lane 2) was served as a control. The red arrows indicate the sample showed observable mRNA leakage. **g,** Binding and condensation profiles of PoLixNano or LNP towards mRNA, investigated by the agarose gel retardation assay. The formulations were incubated with increasing amounts of sodium dodecyl sulfonate (SDS) to competitively release the encapsulated mRNA within indicated formulations. Naked mRNA was used as a positive control (mRNA lane) and untreated formulations (PoLixNano or LNP lanes) were served as negative controls. The red arrows indicate the initial sample that shows observable leakage. **h,** The mean square displacements (MSD) and diffusion coefficient (Deff, μm²/s⁻¹) of PoLixNano or LNP nanoparticles after nebulization (n > 970 independent nanoparticles per experiment). **i,** Representative trajectories of PoLixNano and LNP nanoparticles before nebulization in a 3% (w/v) mucin solution over a 15-s period, with corresponding MSD and Deff of the PoLixNano or LNP nanoparticles (n > 480 independent nanoparticles per experiment). Statistical significance was evaluated at 13-s (h) and 14-s (i) using two-tailed unpaired *t*-tests. **j,** 3D fluorescence images and sectional fluorescence images illustrating the penetrating profiles of PoLixNano or LNP pre-nebulization in a mucus-simulating hydrogel, accompanied by the corresponding quantitative comparation of penetration depths. **k,** 3D confocal microscopy of the mucus penentrating profiles of PoLixNano or LNP before nebulization in Calu-3 cells with secreted thick mucus layers. Green channel, fluorescence-labeled nanoparticles; blue channel, nuclei stained by DAPI. The results of **f**, **g**, **j** and **k** are representative images from at least three independent experiments with similar results. Statistical significance was analyzed using two-tailed unpaired *t*-tests.

**Extended Data Fig. 4.**
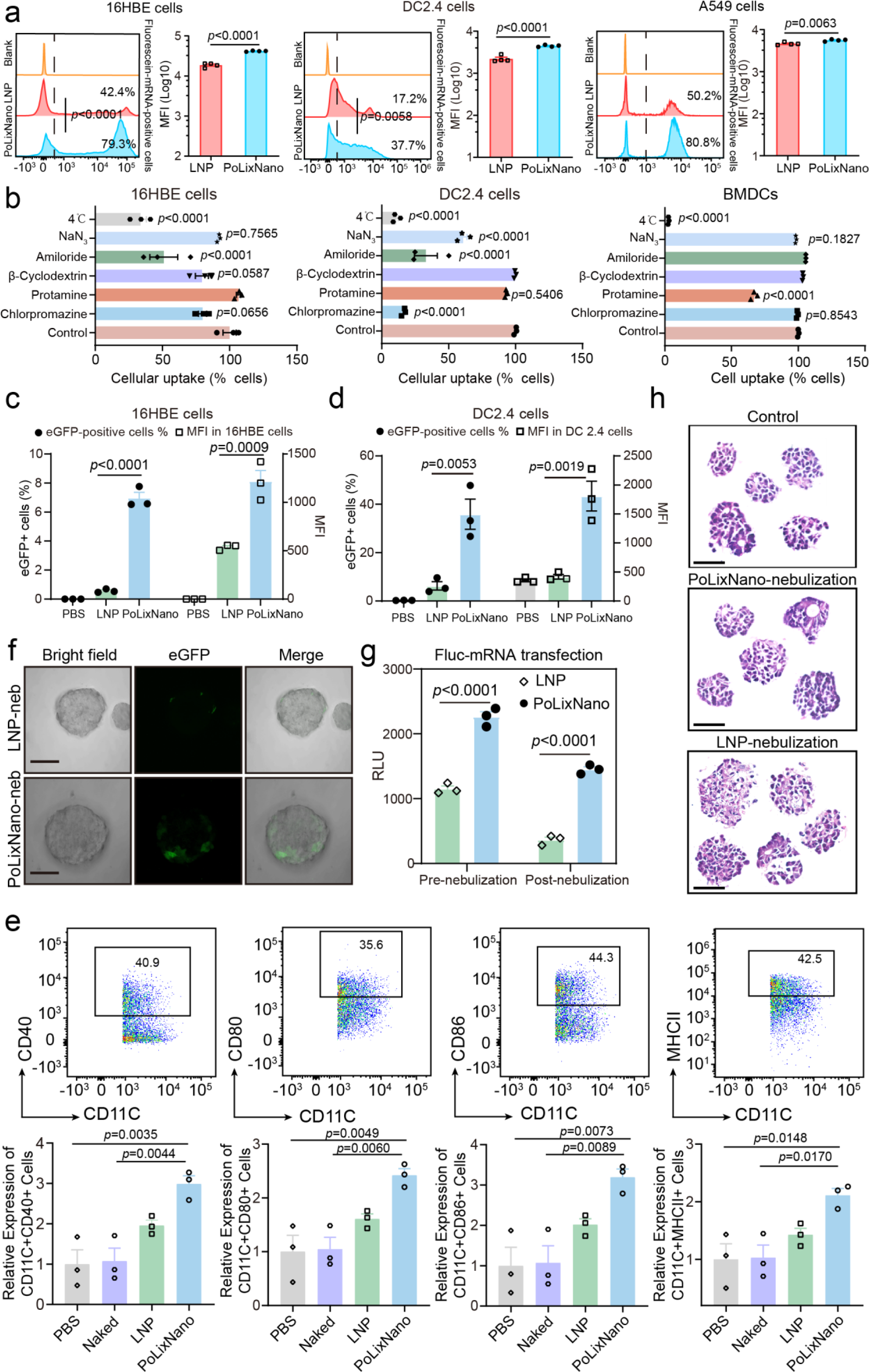
Additional in vitro investigations of the best-performing PoLixNano. **a**, The cellular uptake of Fluorescein-mRNA encapsulated in PoLixNano or LNP counterpart in 16HBE, DC2.4 and A549 cells measured by flow cytometry (n=4 biologically independent samples). The bar chart indicates the mean fluorescence intensity (MFI) per cell, and histograms display the percentage of Fluorescein-mRNA positive cells. Untreated cells were used as the blank control. **b**, Effect of low temperature (4℃) and endocytosis inhibitors on cellular uptake of PoLixNano in 16HBE, DC2.4 and BMDC cells. The non-inhibited internalization of Fluorescein-mRNA@PoLixNano was served as the control group. PoLixNano could utilize efficient energy-dependent cellular uptake mediated by macropinocytosis in 16HBE and DC2.4 cells since its endocytosis was significantly inhibited at 4℃ or by co-incubation with amiloride. Pre-treatment with sodium azide, a cytochrome oxidase inhibitor, or protamine which inhibits adsorptive-mediated endocytosis, substantially decreased the percentage of Fluorescein-mRNA positive cells in BMDCs. A lipid raft-mediated endocytosis inhibitor, β-cyclodextrin, suppressed the internalization of PoLixNano in 16HBE cells, and chlorpromazine (a clathrin-mediated endocytosis inhibitor) significantly down-regulated the uptake of PoLixNano in DC2.4 cells. (n=3 biologically independent replicates). **c**,**d**, Quantitative evaluating the expression levels of eGFP-mRNA in PoLixNano or LNP transfected 16HBE cells (c) and DC2.4 cells (d) via flow cytometry (n=3 biologically independent replicates). **e**, Flow cytometry analysis of CD40, CD80, CD86, and MHCII expression on BMDCs after incubation with PoLixNano or LNP for 24 h. Naked mRNA (“Naked”) and PBS treated counterparts were served as controls (n=4 biologically independent samples). **f**, Representative images showing eGFP-mRNA expression in individual organoids after 36 h treatment with nebulized PoLixNano or LNP-neb. Images from left to right depict bright-field microscopy images of individual organoid, eGFP signal (green channel), and an overlay of the two images. Scale bars: 50 µm. **g**, Fluc-mRNA transfection efficiency in organoids mediated by PoLixNano or LNP both pre- and post-nebulization. The luciferase activity was represented as RLU (n=3 biologically independent samples). **h**, Hematoxylin and eosin (H&E) analysis of organoids treated with PoLixNano or LNP at 24 h post-transfection. Untreated organoids were used control. Scale bars: 50 µm. The results of **f** and **h** are representative images from three independent experiments with similar results. Statistical analysis were determined by a two-tailed unpaired *t*-test in **a** and **g**, a one-way ANOVA with Dunnett’s post-hoc test was applied in **b**, and a one-way ANOVA with Tukey’s post-hoc test was applied in **c**, **d** and **e.**

**Extended Data Fig. 5.**
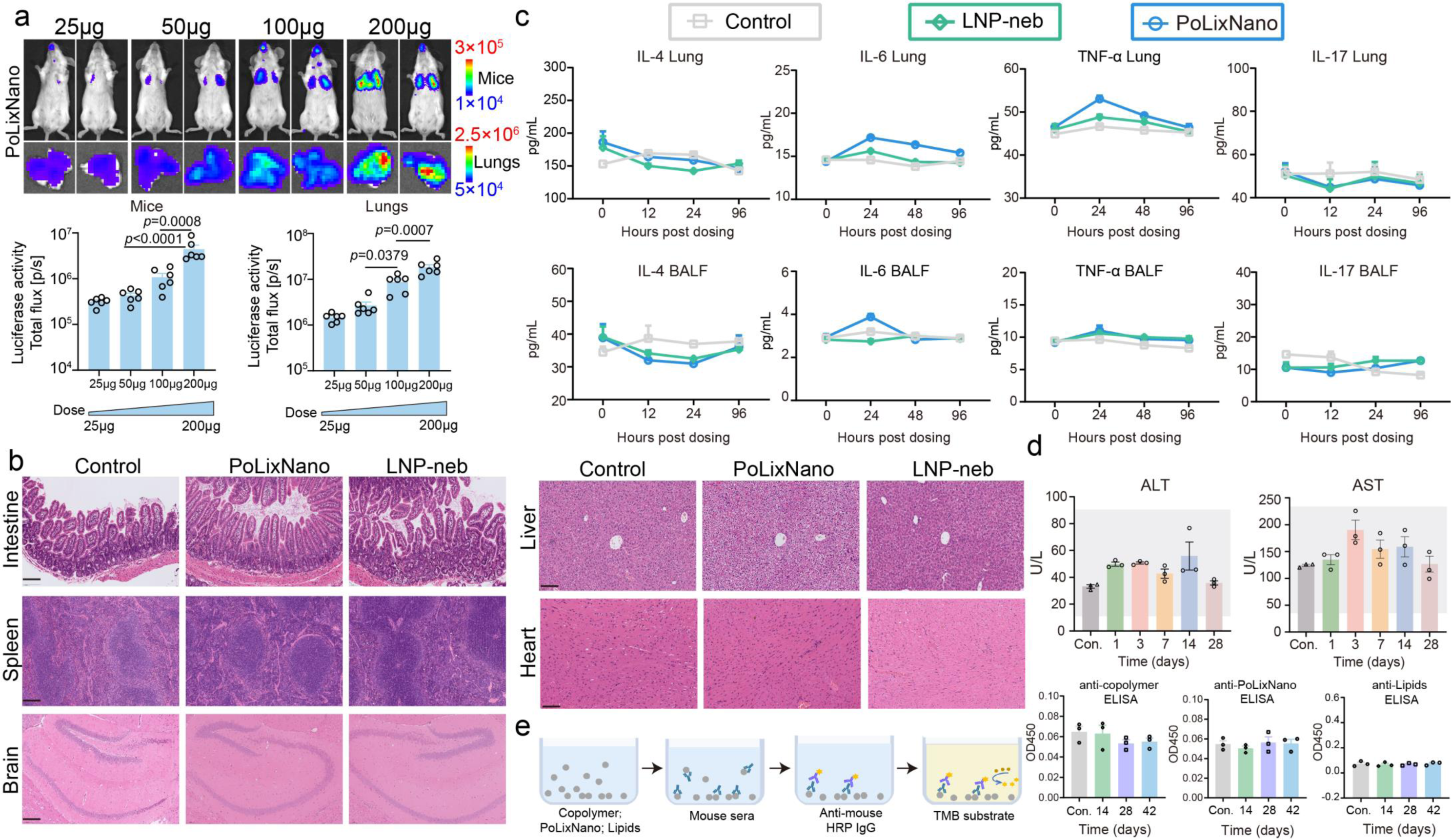
Additional in vivo evaluations of PoLixNano. **a,** Bioluminescence imaging and quantitative analysis of mice and excised lungs at 6 h post-nebulization with PoLixNano encapsulating 25 μg/mouse, 50 μg/mouse, 100 μg/mouse or 200 μg/mouse Fluc-mRNA, respectively (n=6 biologically independent mice). **b**, Histopathology analysis of major organs of mice, including intestine, liver, spleen, heart and brain, collected from mice nebulized with PoLixNano or LNP at 24 h post dosing. Scale bar: 100 μm. Samples collected from PBS treated mice were used as the control. The results of **a** and **b** are representative images from at least three independent experiments with similar results. **c**, Enzyme-linked immunoassay (ELISA) quantification of interleukin-4 (IL-4), interleukin-6 (IL-6), tumor necrosis factor-α (TNF-α) and interleukin-17 (IL-17) in lung homogenates and BALF samples of mice treated with nebulized PoLixNano or LNP (LNP-neb), measured at indicated time points (n=3 biologically independent samples). **d**, Complete blood chemistry metrics of mice treated with nebulized PoLixNano (n=3 biologically independent samples). The grey regions represent the normal range of each parameter as a reference. ALT: alanine aminotransferase, AST: aspartate aminotransferase. **e**, Schematic (left panel) and quantitation (right panel) of anti-PoLixNano antibody responses at 14 days, 28 days and 42 days post-dosing of PoLixNano via ELISA analysis. Horseradish peroxidase (HRP); 3,3′,5,5′-Tetramethylbenzidine (TMB) (n=3 biologically independent samples). Untreated mice were used as the control (Con.) in **c**-**e**. A one-way ANOVA with Tukey’s post-hoc test was applied in **a**.

**Extended Data Fig. 6.**
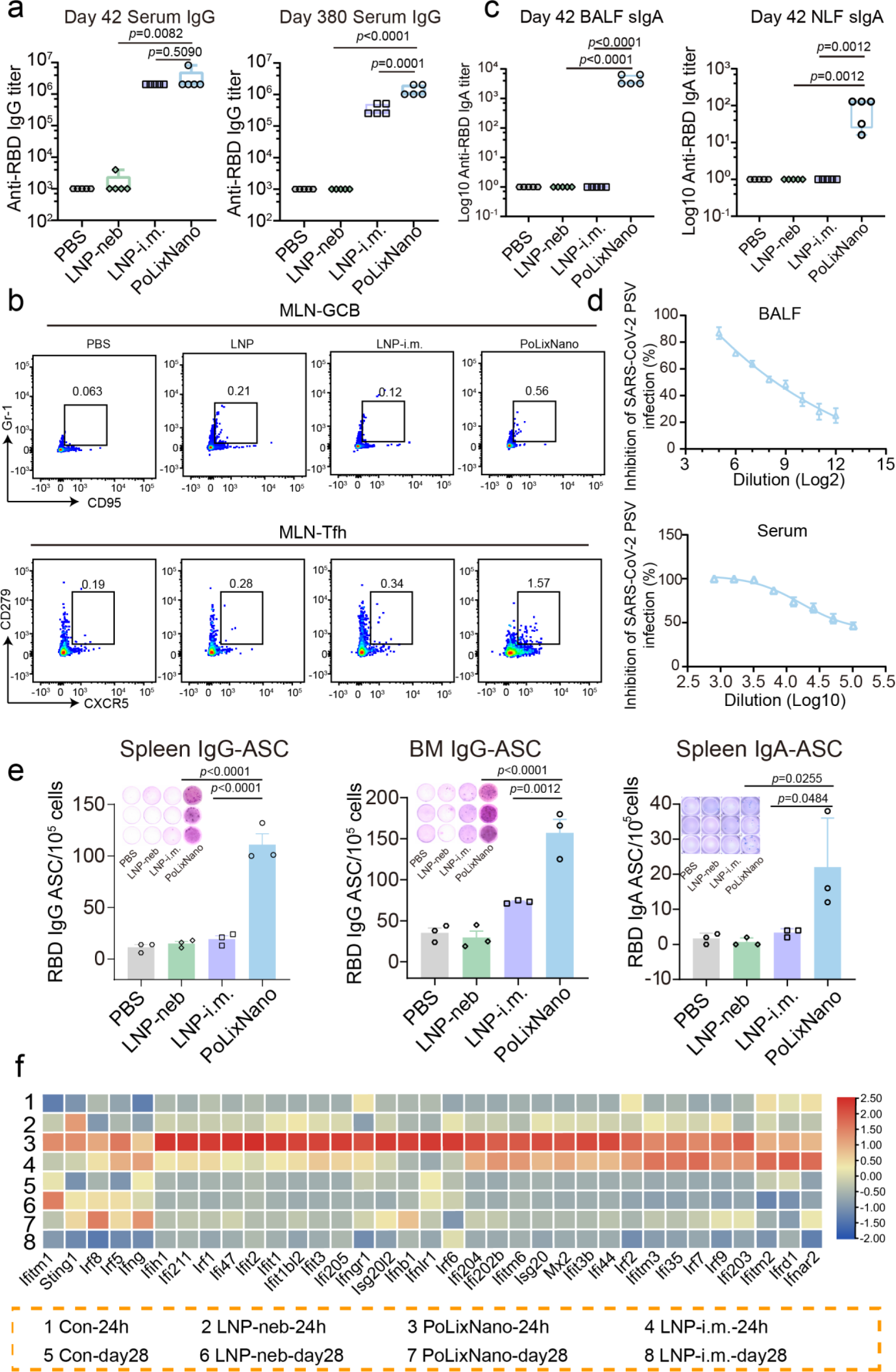
Supplementary evaluations of immune responses in mice induced by different formulations. **a,** Endpoint titers of RBD-specific IgG antibody within serum samples of vaccinated mice collected on day 42 and day 380 measured by ELISA (n=5 biologically independent animals). **b**, Representative flow cytometry plots showing proportion of germinal center B cells (GCB, CD19+B220+CD95+Gr1+) and T follicular helper cells (Tfh, CD4+CXCR5+PD-1+) in mediastinal lymph nodes (MLNs) of mice vaccinated by indicated formulations. **c**, Endpoint titers of RBD-specific sIgA antibody in BALF and NLF samples of vaccinated mice collected on day 42 (n=5 biologically independent animals). **d**, Pseudo-virus neutralizing antibody titer in the BALF and serum samples from PoLixNano vaccinated mice on day 42 post nebulization (n=5 biologically independent anmimals). **e**, ELISpot assay for the quantification of RBD-specific IgG antibody secreting cells (ASCs) in spleen and bone marrow (BM) and RBD-specific IgA ASCs in spleen collected from vaccinated mice on day 35 (n=3 biologically independent samples). Inserts: imagines of each well. **f**, A heatmap showing the expression of genes related to the IFN signaling pathway in lung samples collected at 24 h and 28 days post-prime. Lung samples form mice treated with PBS served as the control (Con). A one-way ANOVA with Tukey’s post-hoc test was used for statistical analysis in **a**, **c** and **e**.

**Extended Data Fig. 7.**
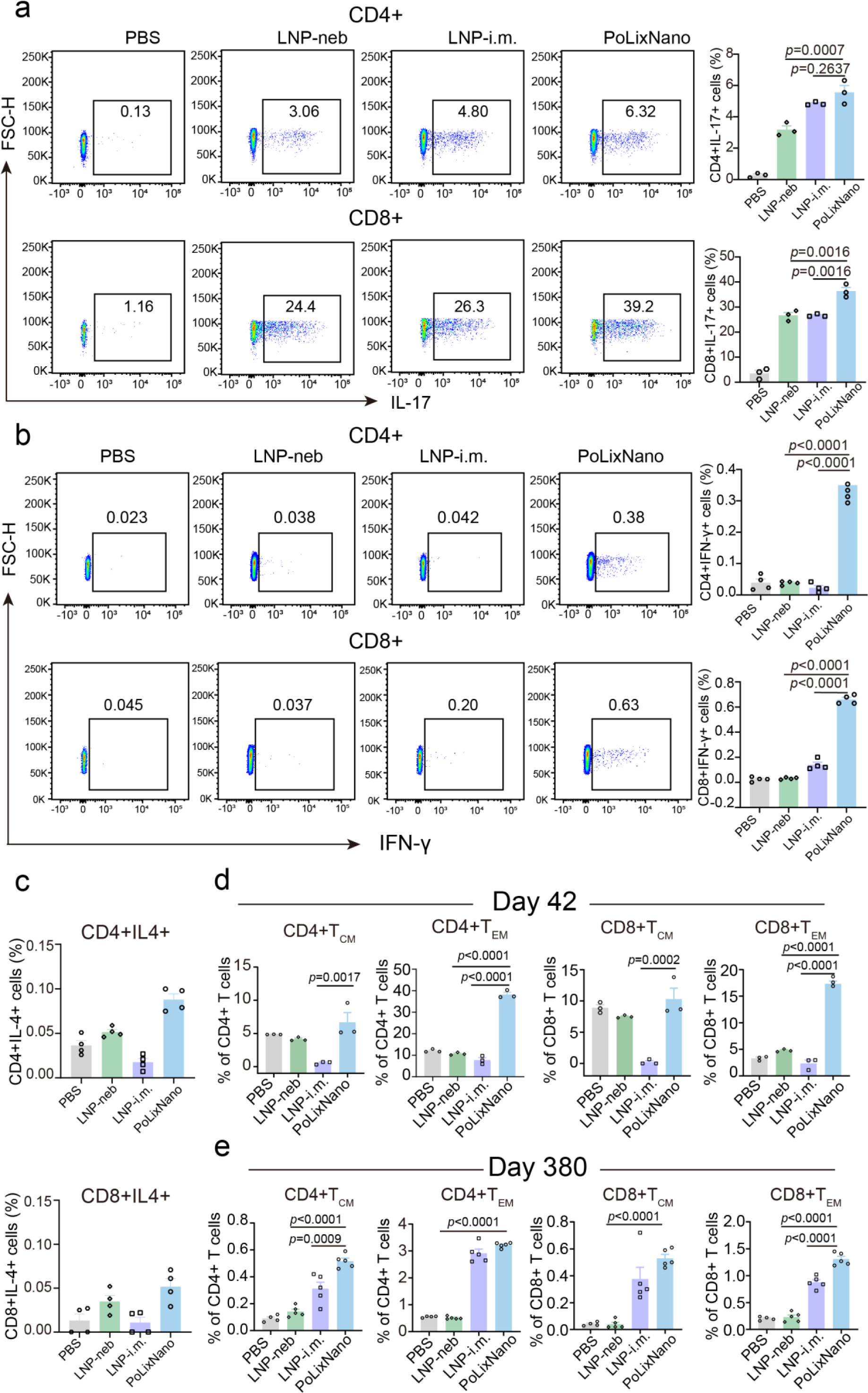
Supplementary evaluations of T helper (Th) cell responses and memory-biased immunity in the lung samples of mice vaccinated by different formulations. **a,** Representative flow cytometry plots (left) and proportions (right) of Th17 (CD4+IL17+) and CD8+IL17+ T cells in the pulmonary lymphocytes obtained from mice vaccinated by indicated formulations 42 days post-prime. **b,** Representative flow cytometry plots (left) and proportions (right) of Th1 (CD4+IFN-γ+) and cytotoxic T (CD8+IFN-γ+) cells in the pulmonary lymphocytes obtained from mice vaccinated by indicated formulations 42 days post-prime. **c**, Quantitative analysis of Th2 (CD4+IL-4+) cells (upper panel) and CD8+IL-4+ T cells (lower panel) in the pulmonary lymphocytes obtained from mice vaccinated by indicated formulations 42 days post-prime. **d** and **e**, The proportion of central memory T cells (T_CM_, CD44+CD62L+), effector memory T cell (T_EM_, CD44+CD62L-) among CD4+ or CD8+ T cells in lungs of mice vaccinated by indicated formulations 42 days (**d**) and 380 days (**e**) post-prime. A one-way ANOVA followed by Tukey’s post-hoc test was applied for statistical analysis in **a**-**e**.

**Extended Data Fig. 8.**
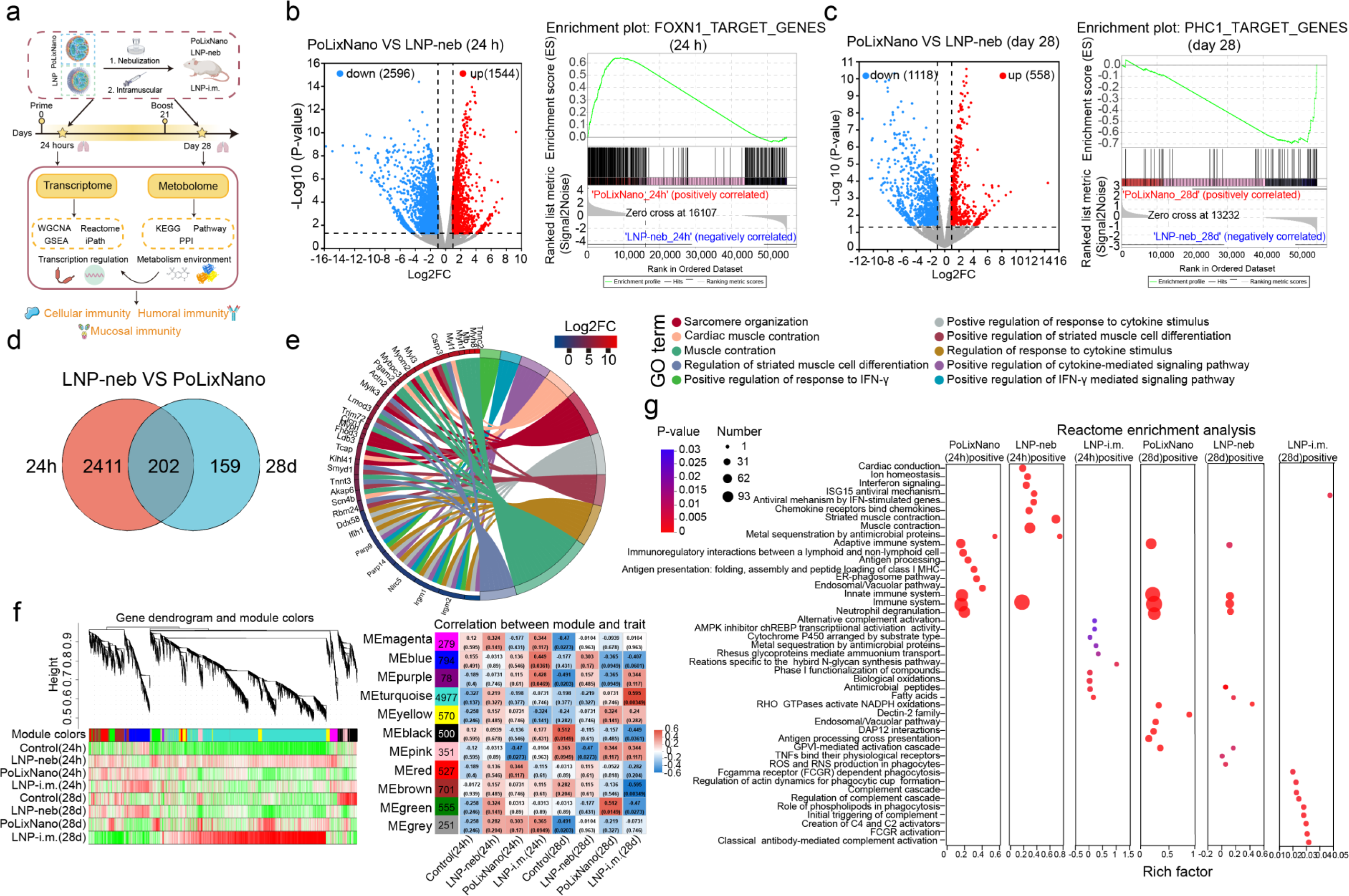
RNA sequencing analysis of lung tissues obtained from mice vaccinated by PoLixNano or LNP counterparts. **a**, Schematic illustration of dosing, sampling and experimental procedure. Samples from mice treated with PBS were served as the control. **b** and **c**, Volcano plot analysis of differentially expressed genes (DEGs) and Gene Set Enrichment Analysis (GSEA) of DEGs. Volcano plots (left) and GSEA (right) of DEGs in lung tissues at 24 h (b) and 28 days (c) post-prime, comparing PoLixNano and LNP-neb. Red dots: up regulation; blue dots: down regulation; gray dots: no significance. Significant DEGs are highlighted with *P* value < 0.05, and |log_2_FC|>1.2. **d**, Venn diagram shows 2613 DEGs between the PoLixNano and LNP-neb groups at 24 h and 361 DEGs at day 28. A total of 202 overlapping DEGs were observed between these time points in PoLixNano and LNP-neb groups. **e**, GO enrichment chord plot illustrating the relationship between the overlapping DEGs and enriched biological pathways. **f,** Weighted Gene Co-Expression Network Analysis (WGCNA) illustrating the association of DEGs with various enriched co-expression modules between various groups at 24 h and 28 days post-vaccination. **g,** Reactome enrichment analysis of genes within feature positive modules associated with PoLixNano, LNP-neb, or LNP-i.m. at 24 h and 28 days post-prime. Upregulated and downregulated DEGs were calculated using Hypergeometric Distribution Algorithm analysis in **e** and **f**. *P* values were adjusted with Benjamini-Hochberg (BH) Method, the reactome enrichment analysis with *P* value <0.05 were considered significantly enriched.

**Extended Data Fig. 9.**
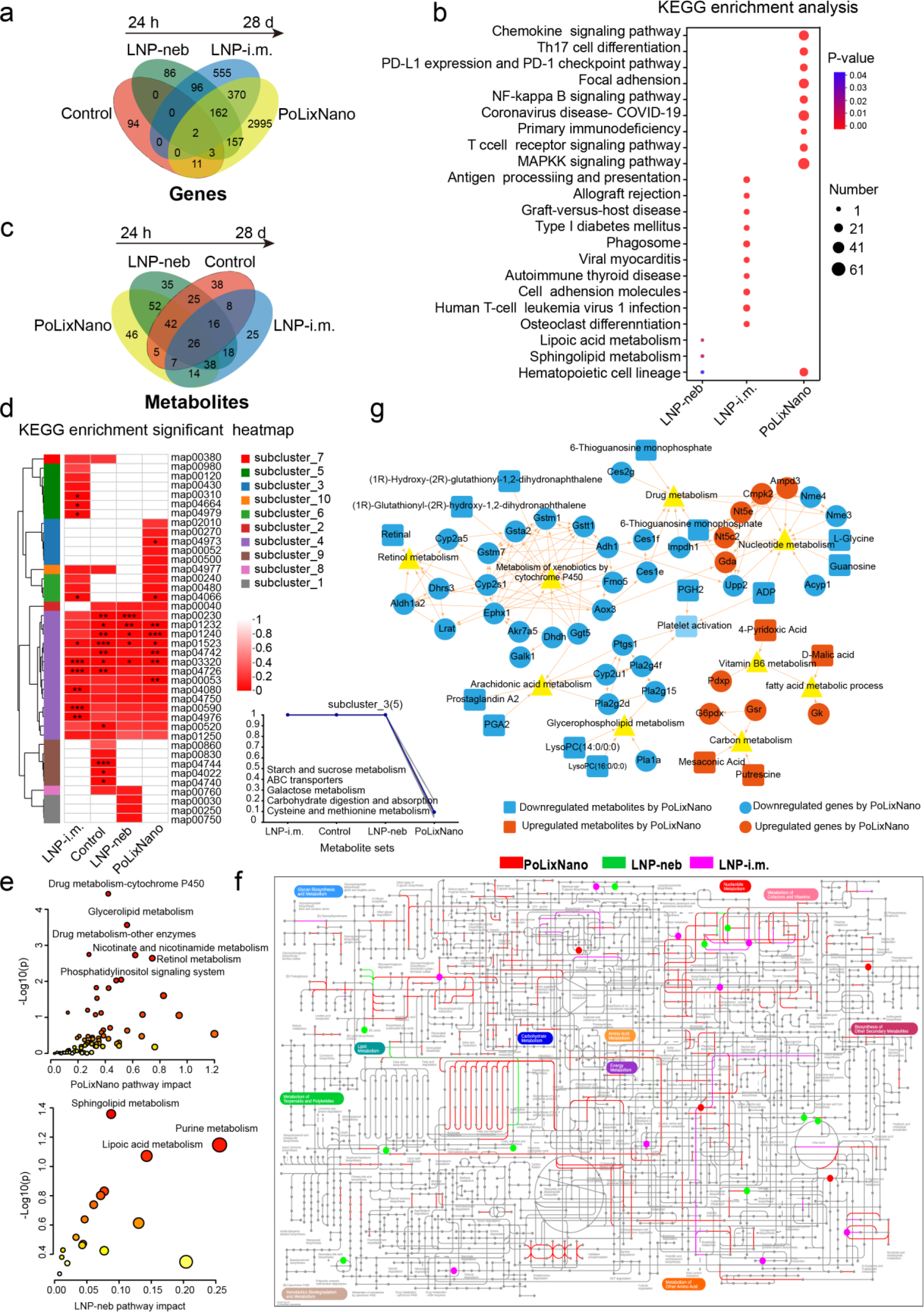
Additional transcriptomic and metabolomic analyses. **a**, Dynamic analysis of DEGs over time from 24 h to 28 days post-prime dosing. Venn diagram depicts various DEGs among the indicated four groups. There are 2995 DGEs between 24 h and 28 days post-vaccination samples for PoLixNano group, 555 DEGs and 86 DEGs for LNP-i.m. and LNP-neb groups respectively. **b**, KEGG terms display enriched pathway of the DEGs between the 24 h and 28 days after initial dosing of the PoLixNano, LNP-neb and LNP-i.m. **c**, Venn diagram shows various different expressed metabolites (DEMs) over time from 24 h to 28 days post-dosing among the indicated four groups. PoLixNano group has 46 DEMs, while the LNP-i.m. and LNP-neb groups have 25 and 35 DEMs respectively. **d**, KEGG terms displays enriched pathways of the DEMs between the 24 h and 28 days after initial vaccination of the PoLixNano, LNP-neb and LNP-i.m.. The KEGG pathway clustering algorithm utilized hierarchical clustering, with the distance metric set to Euclidean and the clustering method chosen as Complete. Enrichment results were clustered using the *P*_Value_uncorrected, with multiple testing correction performed using the Benjamini-Hochberg (BH) method. The data was additionally standardized using Z-score normalization. (\**P*<0.05, \*\**P*<0.01, \*\*\**P*<0.001). **e**, The metabolic pathways enriched in Metabolite cluster 6 and 3. MetaboAnalyst (https://www.metaboanalyst.ca) online analysis was used to evaluate metabolic pathways. **f**, Ingenuity pathway analysis (IPA) (https://pathways.embl.de) highlights metabolic pathways associated within the PoLixNano, LNP-neb, and LNP-i.m. groups. Each colored dot represents a metabolite, and the lines indicate metabolic pathway involving DEGs. **g**, Interactions among DEGs were analyzed using STRING (https://cn.string-db.org/), while relationships between metabolites and associated genes were examined using MetaCyc (https://metacyc.org) and the Human Metabolome Database (HMDB) (https://hmdb.ca/metabolites). Data visualization was performed with Cytoscape. Circles represent DEGs, squares indicate the metabolites, and triangles depict metabolic pathways. Blue means downregulation, and orange denotes upregulation in DEGs or DEMs. All DEGs and DEMs were compared between the PoLixNano and LNP groups at 24 h and 28 days post-prime.

**Extended Data Fig. 10.**
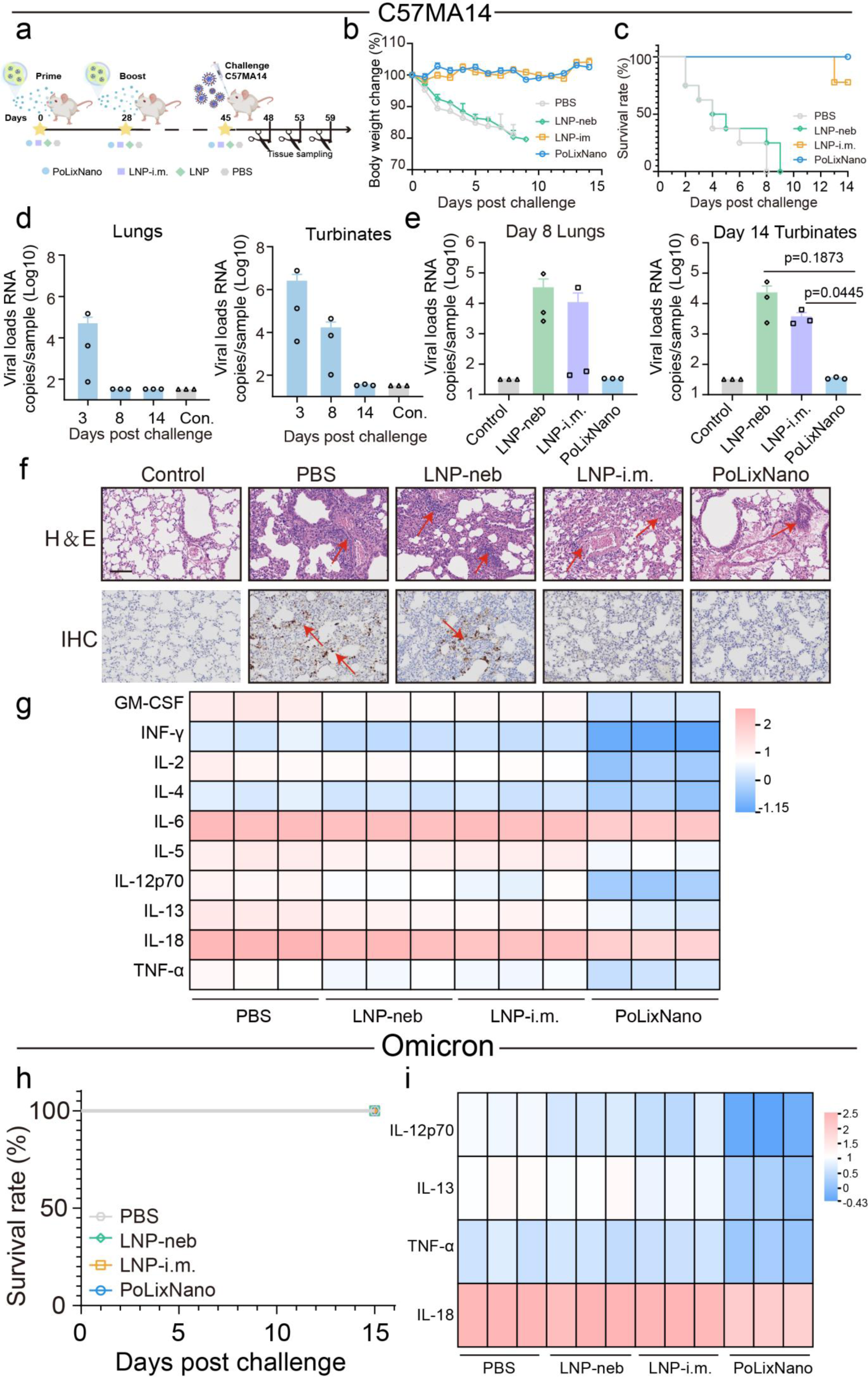
PoLixNano provides robust protection against a lethal challenge of SARS-CoV-2 C57MA14 strain in mice. **a**, Schematic illustration of immunization, sampling and live SARS-CoV-2 challenge in mice with a lethal dose (50 LD_50_) of C57MA14 strain via intranasal instillation 45 days post-prime with PoLixNano: nebulized RBD-mRNA@PoLixNano, LNP-neb: nebulized RBD-mRNA@LNP, LNP-i.m.: intramuscularly injected RBD-mRNA@LNP, and PBS: nebulized PBS. **b**, The body weight changes of mice vaccinated with the above-mentioned formulations after the SARS-CoV-2 challenge (n=10 biologically independent animals). **c**, The survival rate of mice immunized with PoLixNano, LNP-neb, LNP-i.m. and PBS after the C57MA14 strain challenge (n=10 biologically independent animals). **d**, The viral loads in the lung tissues and nasal turbinate of PoLixNano vaccinated mice at 3 days post-infection (dpi), 8 dpi and 14 dpi (n=3 biologically independent experiments). **e**, The viral RNA loads in lung tissues collected at 8 dpi and in nasal turbinates collected at 14 dpi of mice vacicnated with indicated formulations (n=3 biologically independent experiments). **f**, Histopathology analysis (top) and immunohistochemical (IHC) assay of the SARS-CoV-2 nucleocapsid (N) protein (bottom) in lung tissues at 8 dpi (n=3 biologically independent replicates). Red arrows indicate pathological changes and SARS-CoV-2 nucleocapsid (N) protein positive areas. Scale bars, 100 μm. **g**, A heat map showing levels of different cytokines in lung homogenates collected from the indicated groups at 3 dpi with the C57MA14 strain. **h**, The survival rate of mice immunized with PoLixNano, LNP-neb, LNP-i.m. and PBS over 15 days following Omicron BA.5 strain challenge. **i**, A heat map showing the levels of different cytokines in lung homogenates among indicated groups at 3 dpi with the Omicron BA.5 strain. Samples collected from untreated and unchallenged mice were served as the control (Con.) in **d-f**. A one-way ANOVA with Tukey’s post-hoc test was applied for statistical analysis in **e**.

## References

1. Mascola, J. R. & Fauci, A. S. Novel vaccine technologies for the 21st century. Nat Rev Immunol 20, 87–88 (2020).

2. Pardi, N., Hogan, M. J., Porter, F. W. & Weissman, D. mRNA vaccines - a new era in vaccinology. Nat Rev Drug Discov 17, 261–279 (2018).

3. Callaway, E. & Naddaf, M. Pioneers of mRNA COVID vaccines win medicine Nobel. Nature 622, 228–229 (2023).

4. Rohner, E., Yang, R., Foo, K. S., Goedel, A. & Chien, K. R. Unlocking the promise of mRNA therapeutics. Nat Biotechnol 40, 1586–1600 (2022).

5. Chaudhary, N., Weissman, D. & Whitehead, K. A. mRNA vaccines for infectious diseases: principles, delivery and clinical translation. Nat Rev Drug Discov 20, 817–838 (2021).

6. Liu, C. et al. mRNA-based cancer therapeutics. Nat Rev Cancer 23, 526–543 (2023).

7. Martini, P. G. V. & Guey, L. T. A New Era for Rare Genetic Diseases: Messenger RNA Therapy. Hum Gene Ther 30, 1180–1189 (2019).

8. Hameed, S. A., Paul, S., Dellosa, G. K. Y., Jaraquemada, D. & Bello, M. B. Towards the future exploration of mucosal mRNA vaccines against emerging viral diseases; lessons from existing next-generation mucosal vaccine strategies. NPJ Vaccines 7, 71 (2022).

9. Suberi, A. et al. Polymer nanoparticles deliver mRNA to the lung for mucosal vaccination. Sci Transl Med 15, eabq0603 (2023).

10. Baldeon Vaca, G., et al. Intranasal mRNA-LNP vaccination protects hamsters from SARS-CoV-2 infection. Sci Adv 9, eadh1655 (2023).

11. Lavelle, E. C. & Ward, R. W. Mucosal vaccines - fortifying the frontiers. Nat Rev Immunol 22, 236–250 (2022).

12. Sun, S. et al. Respiratory mucosal vaccination of peptide-poloxamine-DNA nanoparticles provides complete protection against lethal SARS-CoV-2 challenge. Biomaterials 292, 121907 (2023).

13. Liu, M., Hu, S., Yan, N., Popowski, K. D. & Cheng, K. Inhalable extracellular vesicle delivery of IL-12 mRNA to treat lung cancer and promote systemic immunity. Nat Nanotechnol 19, 565–575 (2024).

14. Guan, S. et al. Self-assembled peptide-poloxamine nanoparticles enable in vitro and in vivo genome restoration for cystic fibrosis. Nat Nanotechnol 14, 287–297 (2019).

15. Guan, S., Darmstädter, M., Xu, C. & Rosenecker, J. In Vitro Investigations on Optimizing and Nebulization of IVT-mRNA Formulations for Potential Pulmonary-Based Alpha-1-Antitrypsin Deficiency Treatment. Pharmaceutics 13, 1281 (2021).

16. Paff, T., Omran, H., Nielsen, K. G. & Haarman, E. G. Current and Future Treatments in Primary Ciliary Dyskinesia. Int J Mol Sci 22, 9834 (2021).

17. Guan, S. & Rosenecker, J. Nanotechnologies in delivery of mRNA therapeutics using nonviral vector-based delivery systems. Gene Ther 24, 133–143 (2017).

18. Kim, J., Eygeris, Y., Ryals, R. C., Jozić, A. & Sahay, G. Strategies for non-viral vectors targeting organs beyond the liver. Nat. Nanotechnol. 19, 428–447 (2024).

19. Tang, J. et al. Nanotechnologies in Delivery of DNA and mRNA Vaccines to the Nasal and Pulmonary Mucosa. Nanomaterials (Basel) 12, 226 (2022).

20. Johler, S. M., Rejman, J., Guan, S. & Rosenecker, J. Nebulisation of IVT mRNA Complexes for Intrapulmonary Administration. PLoS One 10, e0137504 (2015).

21. Lokugamage, M. P. et al. Optimization of lipid nanoparticles for the delivery of nebulized therapeutic mRNA to the lungs. Nat Biomed Eng 5, 1059–1068 (2021).

22. Blanchard, E. L. et al. Treatment of influenza and SARS-CoV-2 infections via mRNA-encoded Cas13a in rodents. Nat Biotechnol 39, 717–726 (2021).

23. Kim, N., Duncan, G. A., Hanes, J. & Suk, J. S. Barriers to inhaled gene therapy of obstructive lung diseases: A review. J Control Release 240, 465–488 (2016).

24. Huck, B. C. et al. Models using native tracheobronchial mucus in the context of pulmonary drug delivery research: Composition, structure and barrier properties. Adv Drug Deliv Rev 183, 114141 (2022).

25. Chen, D., Liu, J., Wu, J. & Suk, J. S. Enhancing nanoparticle penetration through airway mucus to improve drug delivery efficacy in the lung. Expert Opin Drug Deliv 18, 595– 606 (2021).

26. Cullis, P. R. & Felgner, P. L. The 60-year evolution of lipid nanoparticles for nucleic acid delivery. Nat Rev Drug Discov 23, 709–722 (2024).

27. Barbier, A. J., Jiang, A. Y., Zhang, P., Wooster, R. & Anderson, D. G. The clinical progress of mRNA vaccines and immunotherapies. Nat Biotechnol 40, 840–854 (2022).

28. Rowe, S. M. et al. Inhaled mRNA therapy for treatment of cystic fibrosis: Interim results of a randomized, double-blind, placebo-controlled phase 1/2 clinical study. J Cyst Fibros 22, 656–664 (2023).

29. Jiang, A. Y. et al. Combinatorial development of nebulized mRNA delivery formulations for the lungs. Nat Nanotechnol 19, 364–375 (2024).

30. Kim, J. et al. Engineering Lipid Nanoparticles for Enhanced Intracellular Delivery of mRNA through Inhalation. ACS Nano 16, 14792–14806 (2022).

31. Liu, S. et al. Charge-assisted stabilization of lipid nanoparticles enables inhaled mRNA delivery for mucosal vaccination. Preprint at 10.26434/chemrxiv-2023-n079h (2023).

32. Bai, X. et al. Optimized inhaled LNP formulation for enhanced treatment of idiopathic pulmonary fibrosis via mRNA-mediated antibody therapy. Nat Commun 15, 6844 (2024).

33. Li, B. et al. Accelerating ionizable lipid discovery for mRNA delivery using machine learning and combinatorial chemistry. Nat Mater 23, 1002–1008 (2024).

34. Rao, L., Yuan, Y., Shen, X., Yu, G. & Chen, X. Designing nanotheranostics with machine learning. Nat Nanotechnol (2024) doi:10.1038/s41565-024-01753-8.

35. Dumortier, G., Grossiord, J. L., Agnely, F. & Chaumeil, J. C. A review of poloxamer 407 pharmaceutical and pharmacological characteristics. Pharm Res 23, 2709–2728 (2006).

36. Caballero, I. et al. Tetrafunctional Block Copolymers Promote Lung Gene Transfer in Newborn Piglets. Mol Ther Nucleic Acids 16, 186–193 (2019).

37. Schneider, C. S. et al. Nanoparticles that do not adhere to mucus provide uniform and long-lasting drug delivery to airways following inhalation. Sci Adv 3, e1601556 (2017).

38. Fujii, M. & Sato, T. Somatic cell-derived organoids as prototypes of human epithelial tissues and diseases. Nat Mater 20, 156–169 (2021).

39. Kim, J., Koo, B.-K. & Knoblich, J. A. Human organoids: model systems for human biology and medicine. Nat Rev Mol Cell Biol 21, 571–584 (2020).

40. Ma, X. et al. Nanoparticle Vaccines Based on the Receptor Binding Domain (RBD) and Heptad Repeat (HR) of SARS-CoV-2 Elicit Robust Protective Immune Responses. Immunity 53, 1315–1330.e9 (2020).

41. Afkhami, S. et al. Respiratory mucosal delivery of next-generation COVID-19 vaccine provides robust protection against both ancestral and variant strains of SARS-CoV-2. Cell 185, 896–915.e19 (2022).

42. Ye, T. et al. Inhaled SARS-CoV-2 vaccine for single-dose dry powder aerosol immunization. Nature 624, 630–638 (2023).

43. Robinson, M. J., et al. Long-lived plasma cells accumulate in the bone marrow at a constant rate from early in an immune response. Sci Immunol 7, eabm8389 (2022).

44. Liao, M. et al. Single-cell landscape of bronchoalveolar immune cells in patients with COVID-19. Nat Med 26, 842–844 (2020).

45. Grifoni, A. et al. Targets of T Cell Responses to SARS-CoV-2 Coronavirus in Humans with COVID-19 Disease and Unexposed Individuals. Cell 181, 1489–1501.e15 (2020).

46. Frye, R. F., Schneider, V. M., Frye, C. S. & Feldman, A. M. Plasma levels of TNF-alpha and IL-6 are inversely related to cytochrome P450-dependent drug metabolism in patients with congestive heart failure. J Card Fail 8, 315–319 (2002).

47. Manikandan, P. & Nagini, S. Cytochrome P450 Structure, Function and Clinical Significance: A Review. Curr Drug Targets 19, 38–54 (2018).

48. Žuklys, S., et al. Foxn1 regulates key target genes essential for T cell development in postnatal thymic epithelial cells. Nat Immunol 17, 1206–1215 (2016).

49. Zhu, H. & Zheng, C. When PARPs Meet Antiviral Innate Immunity. Trends Microbiol 29, 776–778 (2021).

50. Xu, H. et al. Molecular and clinical characterization of PARP9 in gliomas: A potential immunotherapeutic target. CNS Neurosci Ther 26, 804–814 (2020).

51. Raupov, R., Suspitsin, E., Belozerov, K., Gabrusskaya, T. & Kostik, M. IFIH1 and DDX58 gene variants in pediatric rheumatic diseases. World J Clin Pediatr 12, 107–114 (2023).

52. Kim, D. et al. Nod2-mediated recognition of the microbiota is critical for mucosal adjuvant activity of cholera toxin. Nat Med 22, 524–530 (2016).

53. Strober, W. & Watanabe, T. NOD2, an intracellular innate immune sensor involved in host defense and Crohn’s disease. Mucosal Immunol 4, 484–495 (2011).

54. Liu, Z. et al. The role of NOD2 in intestinal immune response and microbiota modulation: A therapeutic target in inflammatory bowel disease. Int Immunopharmacol 113, 109466 (2022).

55. Zhou, Y. et al. RIPK3 signaling and its role in regulated cell death and diseases. Cell Death Discov 10, 200 (2024).

56. Nachbur, U. et al. A RIPK2 inhibitor delays NOD signalling events yet prevents inflammatory cytokine production. Nat Commun 6, 6442 (2015).

57. Geddes, K. et al. Identification of an innate T helper type 17 response to intestinal bacterial pathogens. Nat Med 17, 837–844 (2011).

58. Veldhoen, M. Interleukin 17 is a chief orchestrator of immunity. Nat Immunol 18, 612– 621 (2017).

59. Ferreira, M. C. et al. Interleukin-17-induced protein lipocalin 2 is dispensable for immunity to oral candidiasis. Infect Immun 82, 1030–1035 (2014).

60. Christensen, D., Mortensen, R., Rosenkrands, I., Dietrich, J. & Andersen, P. Vaccine-induced Th17 cells are established as resident memory cells in the lung and promote local IgA responses. Mucosal Immunol 10, 260–270 (2017).

61. Stetson, D. B. & Medzhitov, R. Type I interferons in host defense. Immunity 25, 373–381 (2006).

62. Casanova, J.-L., MacMicking, J. D. & Nathan, C. F. Interferon-γ and infectious diseases: Lessons and prospects. Science 384, eadl2016 (2024).

63. Uhl, L. F. K. et al. Interferon-γ couples CD8+ T cell avidity and differentiation during infection. Nat Commun 14, 6727 (2023).

64. Sun, L., Su, Y., Jiao, A., Wang, X. & Zhang, B. T cells in health and disease. Signal Transduct Target Ther 8, 235 (2023).

65. Zhu, M.-L., Nagavalli, A. & Su, M. A. Aire deficiency promotes TRP-1-specific immune rejection of melanoma. Cancer Res 73, 2104–2116 (2013).

## References in Methods

1. Zhang, J.J., et al. Airway applied IVT mRNA vaccine needs specific sequence design and high standard purification that removes devastating dsRNA contaminant. bioRxiv (2024).

2. Xiao, Q. et al. Efficient Transfection of In vitro Transcribed mRNA in Cultured Cells Using Peptide-Poloxamine Nanoparticles. J Vis Exp (2022).

3. Bai, X. et al. Optimized inhaled LNP formulation for enhanced treatment of idiopathic pulmonary fibrosis via mRNA-mediated antibody therapy. Nature communications 15, 6844 (2024).

4. Liu, M. et al. Efficient mucus permeation and tight junction opening by dissociable “mucus-inert” agent coated trimethyl chitosan nanoparticles for oral insulin delivery. J Control Release 222, 67–77 (2016).

5. Zhang, Y.D. et al. Rationally Designed Self-Assembling Nanoparticles to Overcome Mucus and Epithelium Transport Barriers for Oral Vaccines against. Adv Funct Mater 28 (2018).

6. Zhang, L. et al. Effect of mRNA-LNP components of two globally-marketed COVID-19 vaccines on efficacy and stability. NPJ vaccines 8, 156 (2023).

7. Zhong, J. et al. Hyodeoxycholic acid ameliorates nonalcoholic fatty liver disease by inhibiting RAN-mediated PPARalpha nucleus-cytoplasm shuttling. Nature communications 14, 5451 (2023).

8. Kim, D., Langmead, B. & Salzberg, S.L. HISAT: a fast spliced aligner with low memory requirements. Nature methods 12, 357–360 (2015).

9. Love, M.I., Huber, W. & Anders, S. Moderated estimation of fold change and dispersion for RNA-seq data with DESeq2. Genome biology 15, 550 (2014).

10. Wang, L., Feng, Z., Wang, X., Wang, X. & Zhang, X. DEGseq: an R package for identifying differentially expressed genes from RNA-seq data. Bioinformatics 26, 136–138 (2010).

11. Xie, C. et al. KOBAS 2.0: a web server for annotation and identification of enriched pathways and diseases. Nucleic acids research 39, W316–322 (2011).

12. Sun, X. et al. Mirabegron displays anticancer effects by globally browning adipose tissues. Nature communications 14, 7610 (2023).

13. Wishart, D.S. et al. HMDB 5.0: the Human Metabolome Database for 2022. Nucleic acids research 50, D622–D631 (2022).

14. Ren, Y., et al. Majorbio Cloud: A one-stop, comprehensive bioinformatic platform for multiomics analyses. iMeta 1, e12 (2022).

15. Sun, S. et al. Respiratory mucosal vaccination of peptide-poloxamine-DNA nanoparticles provides complete protection against lethal SARS-CoV-2 challenge. Biomaterials 292, 121907 (2023).

16. Guan, S. et al. Self-assembled peptide-poloxamine nanoparticles enable in vitro and in vivo genome restoration for cystic fibrosis. Nat Nanotechnol 14, 287–297 (2019).

