## Supplementary information for "Machine learning engineered PoLixNano nanoparticles overcome delivery barriers for nebulized mRNA therapeutics"

**This file includes:**

Supplementary Table 1-3

Supplementary Figure 1-13

**SUPPLEMENTARY TABLES**

**Supplementary Table 1.** The open reading frame (ORF) sequences of in vitro transcribed mRNA adopted in this study

| **mRNA name** | **sequence** |
| --- | --- |
| **Fluc** | ATGGAAGATGCCAAAAACATTAAGAAGGGCCCAGCGCCATTCTACCCACTCGAAGACGGGACCGCCGGCGAGCAGCTGCACAAAGCCATGAAGCGCTACGCCCTGGTGCCCGGCACCATCGCCTTTACCGACGCACATATCGAGGTGGACATTACCTACGCCGAGTACTTCGAGATGAGCGTTCGGCTGGCAGAAGCTATGAAGCGCTATGGGCTGAATACAAACCATCGGATCGTGGTGTGCAGCGAGAATAGCTTGCAGTTCTTCATGCCCGTGTTGGGTGCCCTGTTCATCGGTGTGGCTGTGGCCCCAGCTAACGACATCTACAACGAGCGCGAGCTGCTGAACAGCATGGGCATCAGCCAGCCCACCGTCGTATTCGTGAGCAAGAAAGGGCTGCAAAAGATCCTCAACGTGCAAAAGAAGCTACCGATCATACAAAAGATCATCATCATGGATAGCAAGACCGACTACCAGGGCTTCCAAAGCATGTACACCTTCGTGACTTCCCATTTGCCACCCGGCTTCAACGAGTACGACTTCGTGCCCGAGAGCTTCGACCGGGACAAAACCATCGCCCTGATCATGAACAGTAGTGGCAGTACCGGATTGCCCAAGGGCGTAGCCCTACCGCACCGCACCGCTTGTGTCCGATTCAGTCATGCCCGCGACCCCATCTTCGGCAACCAGATCATCCCCGACACCGCTATCCTCAGCGTGGTGCCATTTCACCACGGCTTCGGCATGTTCACCACGCTGGGCTACTTGATCTGCGGCTTTCGGGTCGTGCTCATGTACCGCTTCGAGGAGGAGCTATTCTTGCGCAGCTTGCAAGACTATAAGATTCAATCTGCCCTGCTGGTGCCCACACTATTTAGCTTCTTCGCTAAGAGCACTCTCATCGACAAGTACGACCTAAGCAACTTGCACGAGATCGCCAGCGGCGGGGCGCCGCTCAGCAAGGAGGTAGGTGAGGCCGTGGCCAAACGCTTCCACCTACCAGGCATCCGCCAGGGCTACGGCCTGACAGAAACAACCAGCGCCATTCTGATCACCCCCGAAGGGGACGACAAGCCTGGCGCAGTAGGCAAGGTGGTGCCCTTCTTCGAGGCTAAGGTGGTGGACTTGGACACCGGTAAGACACTGGGTGTGAACCAGCGCGGCGAGCTGTGCGTCCGTGGCCCCATGATCATGAGCGGCTACGTTAACAACCCCGAGGCTACAAACGCTCTCATCGACAAGGACGGCTGGCTGCACAGCGGCGACATCGCCTACTGGGACGAGGACGAGCACTTCTTCATCGTGGACCGGCTGAAGAGCCTGATCAAATACAAGGGCTACCAGGTAGCCCCAGCCGAACTGGAGAGCATCCTGCTGCAACACCCCAACATCTTCGACGCCGGGGTCGCCGGCCTGCCCGACGACGATGCCGGCGAGCTGCCCGCCGCAGTCGTCGTGCTGGAACACGGTAAAACCATGACCGAGAAGGAGATCGTGGACTATGTGGCCAGCCAGGTTACAACCGCCAAGAAGCTGCGCGGTGGTGTTGTGTTCGTGGACGAGGTGCCTAAAGGACTGACCGGCAAGTTGGACGCCCGCAAGATCCGCGAGATTCTCATTAAGGCCAAGAAGGGCGGCAAGATCGCCGTGTGATAATAG |
| **eGFP** | ATGGTGAGCAAGGGCGAGGAGCTGTTCACCGGGGTGGTGCCCATCCTGGTCGAGCTGGACGGCGACGTAAACGGCCACAAGTTCAGCGTGTCCGGCGAGGGCGAGGGCGATGCCACCTACGGCAAGCTGACCCTGAAGTTCATCTGCACCACCGGCAAGCTGCCCGTGCCCTGGCCCACCCTCGTGACCACCCTGACCTACGGCGTGCAGTGCTTCAGCCGCTACCCCGACCACATGAAGCAGCACGACTTCTTCAAGTCCGCCATGCCCGAAGGCTACGTCCAGGAGCGCACCATCTTCTTCAAGGACGACGGCAACTACAAGACCCGCGCCGAGGTGAAGTTCGAGGGCGACACCCTGGTGAACCGCATCGAGCTGAAGGGCATCGACTTCAAGGAGGACGGCAACATCCTGGGGCACAAGCTGGAGTACAACTACAACAGCCACAACGTCTATATCATGGCCGACAAGCAGAAGAACGGCATCAAGGTGAACTTCAAGATCCGCCACAACATCGAGGACGGCAGCGTGCAGCTCGCCGACCACTACCAGCAGAACACCCCCATCGGCGACGGCCCCGTGCTGCTGCCCGACAACCACTACCTGAGCACCCAGTCCGCCCTGAGCAAAGACCCCAACGAGAAGCGCGATCACATGGTCCTGCTGGAGTTCGTGACCGCCGCCGGGATCACTCTCGGCATGGACGAGCTGTACAAGTAA |
| **RBD** | ATGAGAGTCCAACCAACAGAATCTATTGTTAGATTTCCTAATATTACAAACTTGTGCCCTTTTGATGAAGTTTTTAACGCCACCAGATTTGCATCTGTTTATGCTTGGAACAGGAAGAGAATCAGCAACTGTGTTGCTGATTATTCTGTCCTATATAATCTCGCATCATTTTTCACTTTTAAGTGTTATGGAGTGTCTCCTACTAAATTAAATGATCTCTGCTTTACTAATGTCTATGCAGATTCATTTGTAATTAGAGGTGATGAAGTCAGACAAATCGCTCCAGGGCAAACTGGAAATATTGCTGATTATAATTATAAATTACCAGATGATTTTACAGGCTGCGTTATAGCTTGGAATTCTAACAAGCTTGATTCTAAGGTTAGTGGTAATTATAATTACCTGTATAGATTGTTTAGGAAGTCTAATCTCAAACCTTTTGAGAGAGATATTTCAACTGAAATCTATCAGGCCGGTAACAAACCTTGTAATGGTGTTGCAGGTTTTAATTGTTACTTTCCTTTACAATCATATGGTTTCCAACCCACTAATGGTGTTGGTTACCAACCATACAGAGTAGTAGTACTTTCTTTTGAACTTCTACATGCACCAGCAACTGTTTGTGGACCTAAAAAGTAA |
| **IL-12** | ATGTGTCCTCAGAAGCTAACCATCTCCTGGTTTGCCATCGTTTTGCTGGTGTCTCCACTCATGGCCATGTGGGAGCTGGAGAAAGACGTTTATGTTGTAGAGGTGGACTGGACTCCCGATGCCCCTGGAGAAACAGTGAACCTCACCTGTGACACGCCTGAAGAAGATGACATCACCTGGACCTCAGACCAGAGACATGGAGTCATAGGCTCTGGAAAGACCCTGACCATCACTGTCAAAGAGTTTCTAGATGCTGGCCAGTACACCTGCCACAAAGGAGGCGAGACTCTGAGCCACTCACATCTGCTGCTCCACAAGAAGGAAAATGGAATTTGGTCCACTGAAATTTTAAAAAATTTCAAAAACAAGACTTTCCTGAAGTGTGAAGCACCAAATTACTCCGGACGGTTCACGTGCTCATGGCTGGTGCAAAGAAACATGGACTTGAAGTTCAACATCAAGAGCAGTAGCAGTTCCCCTGACTCTCGGGCAGTGACATGTGGAATGGCGTCTCTGTCTGCAGAGAAGGTCACACTGGACCAAAGGGACTATGAGAAGTATTCAGTGTCCTGCCAGGAGGATGTCACCTGCCCAACTGCCGAGGAGACCCTGCCCATTGAACTGGCGTTGGAAGCACGGCAGCAGAATAAATATGAGAACTACAGCACCAGCTTCTTCATCAGGGACATCATCAAACCAGACCCGCCCAAGAACTTGCAGATGAAGCCTTTGAAGAACTCACAGGTGGAGGTCAGCTGGGAGTACCCTGACTCCTGGAGCACTCCCCATTCCTACTTCTCCCTCAAGTTCTTTGTTCGAATCCAGCGCAAGAAAGAAAAGATGAAGGAGACAGAGGAGGGGTGTAACCAGAAAGGTGCGTTCCTCGTAGAGAAGACATCTACCGAAGTCCAATGCAAAGGCGGGAATGTCTGCGTGCAAGCTCAGGATCGCTATTACAATTCCTCATGCAGCAAGTGGGCATGTGTTCCCTGCAGGGTCCGATCCGGCGGCGGCGGCAGCGGCGGCGGCGGCAGCGGCGGCGGCGGCAGCATGTGTCAATCACGCTACCTCCTCTTTTTGGCCACCCTTGCCCTCCTAAACCACCTCAGTTTGGCCAGGGTCATTCCAGTCTCTGGACCTGCCAGGTGTCTTAGCCAGTCCCGAAACCTGCTGAAGACCACAGATGACATGGTGAAGACGGCCAGAGAAAAACTGAAACATTATTCCTGCACTGCTGAAGACATCGATCATGAAGACATCACACGGGACCAAACCAGCACATTGAAGACCTGTTTACCACTGGAACTACACAAGAACGAGAGTTGCCTGGCTACTAGAGAGACTTCTTCCACAACAAGAGGGAGCTGCCTGCCCCCACAGAAGACGTCTTTGATGATGACCCTGTGCCTTGGTAGCATCTATGAGGACTTGAAGATGTACCAGACAGAGTTCCAGGCCATCAACGCAGCACTTCAGAATCACAACCATCAGCAGATCATTCTAGACAAGGGCATGCTGGTGGCCATCGATGAGCTGATGCAGTCTCTGAATCATAATGGCGAGACTCTGCGCCAGAAACCTCCTGTGGGAGAAGCAGACCCTTACAGAGTGAAAATGAAGCTCTGCATCCTGCTTCACGCCTTCAGCACCCGCGTCGTGACCATCAACAGGGTGATGGGCTATCTGAGCTCCGCCTGA |
| **CFTR** | ATGCAGAGGTCGCCTCTGGAAAAGGCCAGCGTTGTCTCCAAACTTTTTTTCAGCTGGACCAGACCAATTTTGAGGAAAGGATACAGACAGCGCCTGGAATTGTCAGACATATACCAAATCCCTTCTGTTGATTCTGCTGACAATCTATCTGAAAAATTGGAAAGAGAATGGGATAGAGAGCTGGCTTCAAAGAAAAATCCTAAACTCATTAATGCCCTTCGGCGATGTTTTTTCTGGAGATTTATGTTCTATGGAATCTTTTTATATTTAGGGGAAGTCACCAAAGCAGTACAGCCTCTCTTACTGGGAAGAATCATAGCTTCCTATGACCCGGATAACAAGGAGGAACGCTCTATCGCGATTTATCTAGGCATAGGCTTATGCCTTCTCTTTATTGTGAGGACACTGCTCCTACACCCAGCCATTTTTGGCCTTCATCACATTGGAATGCAGATGAGAATAGCTATGTTTAGTTTGATTTATAAGAAGACTTTAAAGCTGTCAAGCCGTGTTCTAGATAAAATAAGTATTGGACAACTTGTTAGTCTCCTTTCCAACAACCTGAACAAATTTGATGAAGGACTTGCATTGGCACATTTCGTGTGGATCGCTCCTTTGCAAGTGGCACTCCTCATGGGGCTAATCTGGGAGTTGTTACAGGCGTCTGCCTTCTGTGGACTTGGTTTCCTGATAGTCCTTGCCCTTTTTCAGGCTGGGCTAGGGAGAATGATGATGAAGTACAGAGATCAGAGAGCTGGGAAGATCAGTGAAAGACTTGTGATTACCTCAGAAATGATTGAAAATATCCAATCTGTTAAGGCATACTGCTGGGAAGAAGCAATGGAAAAAATGATTGAAAACTTAAGACAAACAGAACTGAAACTGACTCGGAAGGCAGCCTATGTGAGATACTTCAATAGCTCAGCCTTCTTCTTCTCAGGGTTCTTTGTGGTGTTTTTATCTGTGCTTCCCTATGCACTAATCAAAGGAATCATCCTCCGGAAAATATTCACCACCATCTCATTCTGCATTGTTCTGCGCATGGCGGTCACTCGGCAATTTCCCTGGGCTGTACAAACATGGTATGACTCTCTTGGAGCAATAAACAAAATACAGGATTTCTTACAAAAGCAAGAATATAAGACATTGGAATATAACTTAACGACTACAGAAGTAGTGATGGAGAATGTAACAGCCTTCTGGGAGGAGGGATTTGGGGAATTATTTGAGAAAGCAAAACAAAACAATAACAATAGAAAAACTTCTAATGGTGATGACAGCCTCTTCTTCAGTAATTTCTCACTTCTTGGTACTCCTGTCCTGAAAGATATTAATTTCAAGATAGAAAGAGGACAGTTGTTGGCGGTTGCTGGATCCACTGGAGCAGGCAAGACTTCACTTCTAATGGTGATTATGGGAGAACTGGAGCCTTCAGAGGGTAAAATTAAGCACAGTGGAAGAATTTCATTCTGTTCTCAGTTTTCCTGGATTATGCCTGGCACCATTAAAGAAAATATCATCTTTGGTGTTTCCTATGATGAATATAGATACAGAAGCGTCATCAAAGCATGCCAACTAGAAGAGGACATCTCCAAGTTTGCAGAGAAAGACAATATAGTTCTTGGAGAAGGTGGAATCACACTGAGTGGAGGTCAACGAGCAAGAATTTCTTTAGCAAGAGCAGTATACAAAGATGCTGATTTGTATTTATTAGACTCTCCTTTTGGATACCTAGATGTTTTAACAGAAAAAGAAATATTTGAAAGCTGTGTCTGTAAACTGATGGCTAACAAAACTAGGATTTTGGTCACTTCTAAAATGGAACATTTAAAGAAAGCTGACAAAATATTAATTTTGCATGAAGGTAGCAGCTATTTTTATGGGACATTTTCAGAACTCCAAAATCTACAGCCAGACTTTAGCTCAAAACTCATGGGATGTGATTCTTTCGACCAATTTAGTGCAGAAAGAAGAAATTCAATCCTAACTGAGACCTTACACCGTTTCTCATTAGAAGGAGATGCTCCTGTCTCCTGGACAGAAACAAAAAAACAATCTTTTAAACAGACTGGAGAGTTTGGGGAAAAAAGGAAGAATTCTATTCTCAATCCAATCAACTCTATACGAAAATTTTCCATTGTGCAAAAGACTCCCTTACAAATGAATGGCATCGAAGAGGATTCTGATGAGCCTTTAGAGAGAAGGCTGTCCTTAGTACCAGATTCTGAGCAGGGAGAGGCGATACTGCCTCGCATCAGCGTGATCAGCACTGGCCCCACGCTTCAGGCACGAAGGAGGCAGTCTGTCCTGAACCTGATGACACACTCAGTTAACCAAGGTCAGAACATTCACCGAAAGACAACAGCATCCACACGAAAAGTGTCACTGGCCCCTCAGGCAAACTTGACTGAACTGGATATATATTCAAGAAGGTTATCTCAAGAAACTGGCTTGGAAATAAGTGAAGAAATTAACGAAGAAGACTTAAAGGAGTGCTTTTTTGATGATATGGAGAGCATACCAGCAGTGACTACATGGAACACATACCTTCGATATATTACTGTCCACAAGAGCTTAATTTTTGTGCTAATTTGGTGCTTAGTAATTTTTCTGGCAGAGGTGGCTGCTTCTTTGGTTGTGCTGTGGCTCCTTGGAAACACTCCTCTTCAAGACAAAGGGAATAGTACTCATAGTAGAAATAACAGCTATGCAGTGATTATCACCAGCACCAGTTCGTATTATGTGTTTTACATTTACGTGGGAGTAGCCGACACTTTGCTTGCTATGGGATTCTTCAGAGGTCTACCACTGGTGCATACTCTAATCACAGTGTCGAAAATTTTACACCACAAAATGTTACATTCTGTTCTTCAAGCACCTATGTCAACCCTCAACACGTTGAAAGCAGGTGGGATTCTTAATAGATTCTCCAAAGATATAGCAATTTTGGATGACCTTCTGCCTCTTACCATATTTGACTTCATCCAGTTGTTATTAATTGTGATTGGAGCTATAGCAGTTGTCGCAGTTTTACAACCCTACATCTTTGTTGCAACAGTGCCAGTGATAGTGGCTTTTATTATGTTGAGAGCATATTTCCTCCAAACCTCACAGCAACTCAAACAACTGGAATCTGAAGGCAGGAGTCCAATTTTCACTCATCTTGTTACAAGCTTAAAAGGACTATGGACACTTCGTGCCTTCGGACGGCAGCCTTACTTTGAAACTCTGTTCCACAAAGCTCTGAATTTACATACTGCCAACTGGTTCTTGTACCTGTCAACACTGCGCTGGTTCCAAATGAGAATAGAAATGATTTTTGTCATCTTCTTCATTGCTGTTACCTTCATTTCCATTTTAACAACAGGAGAAGGAGAAGGAAGAGTTGGTATTATCCTGACTTTAGCCATGAATATCATGAGTACATTGCAGTGGGCTGTAAACTCCAGCATAGATGTGGATAGCTTGATGCGATCTGTGAGCCGAGTCTTTAAGTTCATTGACATGCCAACAGAAGGTAAACCTACCAAGTCAACCAAACCATACAAGAATGGCCAACTCTCGAAAGTTATGATTATTGAGAATTCACACGTGAAGAAAGATGACATCTGGCCCTCAGGGGGCCAAATGACTGTCAAAGATCTCACAGCAAAATACACAGAAGGTGGAAATGCCATATTAGAGAACATTTCCTTCTCAATAAGTCCTGGCCAGAGGGTGGGCCTCTTGGGAAGAACTGGATCAGGGAAGAGTACTTTGTTATCAGCTTTTTTGAGACTACTGAACACTGAAGGAGAAATCCAGATCGATGGTGTGTCTTGGGATTCAATAACTTTGCAACAGTGGAGGAAAGCCTTTGGAGTGATACCACAGAAAGTATTTATTTTTTCTGGAACATTTAGAAAAAACTTGGATCCCTATGAACAGTGGAGTGATCAAGAAATATGGAAAGTTGCAGATGAGGTTGGGCTCAGATCTGTGATAGAACAGTTTCCTGGGAAGCTTGACTTTGTCCTTGTGGATGGGGGCTGTGTCCTAAGCCATGGCCACAAGCAGTTGATGTGCTTGGCTAGATCTGTTCTCAGTAAGGCGAAGATCTTGCTGCTTGATGAACCCAGTGCTCATTTGGATCCAGTAACATACCAAATAATTAGAAGAACTCTAAAACAAGCATTTGCTGATTGCACAGTAATTCTCTGTGAACACAGGATAGAAGCAATGCTGGAATGCCAACAATTTTTGGTCATAGAAGAGAACAAAGTGCGGCAGTACGATTCCATCCAGAAACTGCTGAACGAGAGGAGCCTCTTCCGGCAAGCCATCAGCCCCTCCGACAGGGTGAAGCTCTTTCCCCACCGGAACTCAAGCAAGTGCAAGTCTAAGCCCCAGATTGCTGCTCTGAAAGAGGAGACAGAAGAAGAGGTGCAAGATACAAGGCTTTAG |
| **MetLuc** | ATGGACATCAAGGTGGTGTTCACCCTGGTGTTCAGCGCCCTGGTGCAGGCCAAGAGCACCGAGTTCGACCCCAACATCGACATCGTGGGCCTGGAAGGCAAGTTCGGCATCACCAACCTGGAAACCGACCTGTTCACCATCTGGGAGACCATGGAAGTGATGATCAAGGCCGACATCGCCGACACCGACCGGGCCAGCAACTTCGTGGCCACCGAGACCGACGCCAACCGGGGCAAGATGCCCGGCAAGAAGCTGCCCCTGGCCGTCATCATGGAAATGGAAGCCAACGCCTTCAAGGCCGGCTGCACCCGGGGCTGCCTGATCTGCCTGAGCAAGATCAAGTGCACCGCCAAGATGAAGGTGTACATCCCCGGCAGGTGCCACGACTACGGCGGCGACAAGAAAACCGGCCAGGCCGGCATCGTGGGCGCCATCGTGGACATCCCCGAGATCAGCGGCTTCAAAGAAATGGCCCCCATGGAACAGTTCATCGCCCAGGTGGACAGATGCGCCAGCTGCACCACCGGCTGCCTGAAGGGCCTGGCCAACGTGAAGTGCAGCGAGCTGCTGAAGAAGTGGCTGCCCGACCGCTGCGCCAGCTTCGCCGACAAGATCCAGAAAGAGGTGCACAACATCAAGGGCATGGCCGGCGACAGGTAG |
| **NanoLuc** | ATGGTCTTCACACTCGAAGATTTCGTTGGGGACTGGCGACAGACAGCCGGCTACAACCTGGACCAAGTCCTTGAACAGGGAGGTGTGTCCAGTTTGTTTCAGAATCTCGGGGTGTCCGTAACTCCGATCCAAAGGATTGTCCTGAGCGGTGAAAATGGGCTGAAGATCGACATCCATGTCATCATCCCGTATGAAGGTCTGAGCGGCGACCAAATGGGCCAGATCGAAAAAATTTTTAAGGTGGTGTACCCTGTGGATGATCATCACTTTAAGGTGATCCTGCACTATGGCACACTGGTAATCGACGGGGTTACGCCGAACATGATCGACTATTTCGGACGGCCGTATGAAGGCATCGCCGTGTTCGACGGCAAAAAGATCACTGTAACAGGGACCCTGTGGAACGGCAACAAAATTATCGACGAGCGCCTGATCAACCCCGACGGCTCCCTGCTGTTCCGAGTAACCATCAACGGAGTGACCGGCTGGCGGCTGTGCGAACGCATTCTGGCGCACGAGACAACCCCTAACAAAGGCAGCGGCACCACCTCTGGCACCACAAGACTGCTGTCTGGCCACACCTGTTTCACACTGACCGGCCTGCTGGGCACACTGGTTACAATGGGACTGCTGACCTGATAATAG |

“Fluc” represents firefly luciferase. “eGFP” is the abbreviation of enhanced green fluorescent protein. “RBD” means the receptor binding domain of SARS-CoV-2. “IL-12” is interleukin-12. “CFTR” denotes the cystic fibrosis transmembrane conductance regulator. “MetLuc” represents Metridia luciferase. “NanoLuc” is the abbreviation of nano luciferase.

**Supplementary Table 2. The information on polymeric compounents of PoLixNano with diverse chemical sturctures that applied for machine learning.**

| Polymer candidates | Polarity | Molecular weight | Ethylene oxide (EO) units | Propylene oxide (PO) units | HLB |
| --- | --- | --- | --- | --- | --- |
| T304 | Amphiphilic | 1650 | 4×3.7 | 4×4.3 | 12-18 |
| T701 | Amphiphilic | 3600 | 4×2.1 | 4×14 | 1-7 |
| T704 | Amphiphilic | 5500 | 4×13 | 4×14 | 15 |
| T707 | Amphiphilic | 12500 | 4×54 | 4×13 | 27 |
| T803 | Amphiphilic | 5500 | 4×11 | 4×15 | 12-18 |
| T901 | Amphiphilic | 4700 | 4×2.7 | 4×18.2 | 1-7 |
| T904 | Amphiphilic | 6700 | 4×15 | 4×17 | 12-18 |
| T908 | Amphiphilic | 25000 | 4×114 | 4×21 | >24 |
| T90R4 | Amphiphilic | 6900 | 4×16 | 4×18 | 1-7 |
| T1107 | Amphiphilic | 15000 | 4×60 | 4×20 | 18-24 |
| T1301 | Amphiphilic | 6800 | 4×4 | 4×27 | 1-7 |
| T1304 | Amphiphilic | 10500 | 4×21.4 | 4×27.1 | 12-18 |
| T1307 | Amphiphilic | 18000 | 4×72 | 4×23 | >24 |
| T1508 | Amphiphilic | 29900 | 4×129 | 4×31 | >24 |
| T150R1 | Amphiphilic | 7900 | 4×5 | 4×29 | 1-7 |
| P105 | Amphiphilic | 1900 | 2×11 | 16 | 18.5 |
| P124 | Amphiphilic | 2090-2360 | 2×12 | 20 | 4.5 |
| P181 | Amphiphilic | 2000 | 2×2 | 31 | 1-7 |
| P184 | Amphiphilic | 2900 | 2×13 | 30 | 13 |
| P188 | Amphiphilic | 8400 | 2×76 | 29 | 29 |
| Polymer candidates | Polarity | Molecular weight | Ethylene oxide (EO) units | Propylene oxide (PO) units | HLB |
| P237 | Amphiphilic | 6840-8830 | 2×64 | 37 | 24 |
| P338 | Amphiphilic | 12700-17400 | 2×141 | 44 | 27 |
| P401 | Amphiphilic | 4400 | 2×4 | 68 | 1 |
| P403 | Amphiphilic | 5800 | 2×20 | 70 | 8 |
| P407 | Amphiphilic | 9840-14600 | 2×101 | 56 | 22 |
| P84 | Amphiphilic | 4200 | 2×19 | 43 | 15-18 |
| P85 | Amphiphilic | 4600 | 2×25 | 40 | 12-18 |
| P335 | Amphiphilic | 6500 | 2×74 | 56 | 15 |
| P10R5 | Amphiphilic | 2000 | 22 | 2×8 | 12-18 |
| L31 | Amphiphilic | 1100 | 2×2 | 16 | 3.5 |
| PEG6K | Hydrophilic | 6000 | 136 | _ | n/a |
| PEG20K | Hydrophilic | 20000 | 150 | _ | n/a |
| 4-arm PEG2K | Hydrophilic | 2000 | 4×11 | _ | n/a |
| 4-arm PEG5K | Hydrophilic | 5000 | 4×28 | _ | n/a |
| 4-arm PEG10K | Hydrophilic | 10000 | 4×57 | _ | n/a |
| 4-arm PEG20K | Hydrophilic | 20000 | 4×114 | _ | n/a |
| 8-arm PEG10K | Hydrophilic | 10000 | 8×28 | _ | n/a |
| 8-arm PEG20K | Hydrophilic | 20000 | 8×57 | _ | n/a |
| 8-arm PEG40 | Hydrophilic | 40000 | 8×114 | _ | n/a |
| PPG-200 | Hydrophobic | 200 | _ | 3 | n/a |
| PPG-400 | Hydrophobic | 400 | _ | 7 | n/a |
| Polymer candidates | Polarity | Molecular weight | Ethylene oxide (EO) units | Propylene oxide (PO) units | HLB |
| PPG-600 | Hydrophobic | 600 | _ | 10 | n/a |
| PPG-1K | Hydrophobic | 1000 | _ | 17 | n/a |
| PPG-2K | Hydrophobic | 2000 | _ | 34 | n/a |
| PPG-3K | Hydrophobic | 3000 | _ | 50 | n/a |
| PPG-4K | Hydrophobic | 4000 | _ | 70 | n/a |
| Tween 20 | Amphiphilic | 1226 | _ | _ | 16.7 |
| Tween 40 | Amphiphilic | 840 | _ | _ | 16.9 |
| Tween 60 | Amphiphilic | 609 | _ | _ | 14.9 |
| Tween 80 | Amphiphilic | 429 | _ | _ | 15 |
| Glycerol ethoxylate-co-propoxylate triol | Amphiphilic | 2600 | _ | _ | n/a |
| PLA2K-PEG5K | Amphiphilic | 7000 | 114 | _ | n/a |
| PLA5K-PEG5K | Amphiphilic | 10000 | 114 | _ | n/a |
| PEG5K-b-PCL2K | Amphiphilic | 7000 | 114 | _ | n/a |
| PLGA(50:50) | Amphiphilic | 10000-20000 | _ | _ | n/a |
| PLGA(65:35) | Amphiphilic | 24000-38000 | _ | _ | n/a |
| PLGA(75:25) | Amphiphilic | 4000-15000 | _ | _ | n/a |
| PLGA(90:10) | Amphiphilic | 25000-35000 | _ | _ | n/a |
| PVA-27K | Hydrophilic | 27000 | _ | _ | >13 |
| PVA-47K | Hydrophilic | 47000 | _ | _ | >13 |
| PAM | Hydrophilic | >3000 | _ | _ | n/a |
| PEI | Hydrophilic | 25000 | _ | _ | n/a |
| Branched PEI | Hydrophilic | 25000 | _ | _ | n/a |
| Polymer candidates | Polarity | Molecular weight | Ethylene oxide (EO) units | Propylene oxide (PO) units | HLB |
| PLA10K | Hydrophobic | 10000 | _ | _ | n/a |
| PLA20K | Hydrophobic | 20000 | _ | _ | n/a |
| PLA60K | Hydrophobic | 60000 | _ | _ | n/a |

The P105, P124, P181, P184, P188, P237, P338, P401, P403, P407, P84, P85, P335, P10R5 and L31 represent poloxamer (Pluronic^®^) block copolymers. The poloxamine (Tetronic^®^) block copolymers encompass the T304, T701, T704, T707, T803, T901, T904, T908, T90R4, T1107, T1301, T1304, T1307, T1508 and T150R1. PEG is the polyethylene glycol. PPG represents the polypropylene glycol. PLA means the polylactic acid. PCL is the abbreviation of polycaprolactone. PLGA represents the poly (lactic-co-glycolic acid). PVA denotes the polyvinyl alcohol. PAM is the polyacrylamide. PEI implies the polyethyleneimine. HLB is the hydrophilic-lipophilic balance. "-" represents polymers which do not contain ethylene oxide (EO) units or propylene oxide (PO) units; “n/a” represents data is not applicable.

**Supplementary Table 3. Peptide pools of 14-mer overlapping peptides spanning the SARS-CoV-2 RBD protein**

| NO. | Sequence | NO. | Sequence |
| --- | --- | --- | --- |
| 1 | RVQPTESIVRFPNI | 23 | FTGCVIAWNSNNLD |
| 2 | ESIVRFPNITNLCP | 24 | IAWNSNNLDSKVGG |
| 3 | FPNITNLCPFGEVF | 25 | NNLDSKVGGNYNYL |
| 4 | NLCPFGEVFNATRF | 26 | KVGGNYNYLYRLFR |
| 5 | GEVFNATRFASVYA | 27 | YNYLYRLFRKSNLK |
| 6 | ATRFASVYAWNRKR | 28 | RLFRKSNLKPFERD |
| 7 | SVYAWNRKRISNCV | 29 | SNLKPFERDISTEI |
| 8 | NRKRISNCVADYSV | 30 | FERDISTEIYQAGS |
| 9 | SNCVADYSVLYNSA | 31 | STEIYQAGSTPCNG |
| 10 | DYSVLYNSASFSTF | 32 | QAGSTPCNGVEGFN |
| 11 | YNSASFSTFKCYGV | 33 | PCNGVEGFNCYFPL |
| 12 | FSTFKCYGVSPTKL | 34 | EGFNCYFPLQSYGF |
| 13 | CYGVSPTKLNDLCF | 35 | YFPLQSYGFQPTNG |
| 14 | PTKLNDLCFTNVYA | 36 | SYGFQPTNGVGYQP |
| 15 | DLCFTNVYADSFVI | 37 | PTNGVGYQPYRVVV |
| 16 | NVYADSFVIRGDEV | 38 | GYQPYRVVVLSFEL |
| 17 | SFVIRGDEVRQIAP | 39 | RVVVLSFELLHAPA |
| 18 | GDEVRQIAPGQTGK | 40 | SFELLHAPATVCGP |
| 19 | QIAPGQTGKIADYN | 41 | HAPATVCGPKKSTN |
| 20 | QTGKIADYNYKLPD | 42 | VCGPKKSTNLVKNK |
| 21 | ADYNYKLPDDFTGC | 43 | KSTNLVKNKCVNF |
| 22 | KLPDDFTGCVIAWN |  |  |

**SUPPLEMENTARY FIGURES**

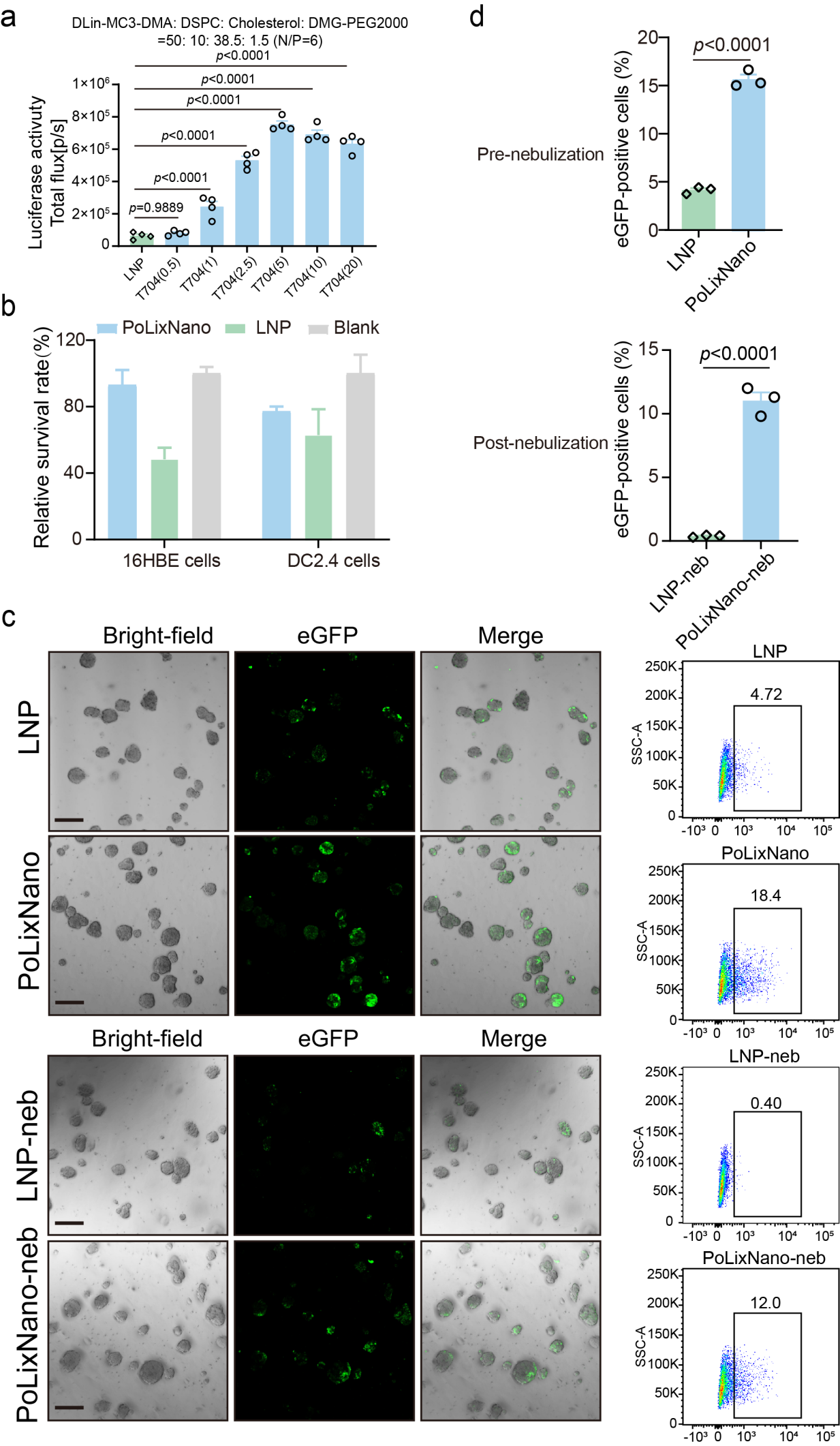

**Supplementary Fig. 1** **|** **a,** Transfection efficiency of mRNA encoding firefly luciferase (Fluc-mRNA) encapsulated in PoLixNano comprising various concentrations of poloxamine 704 (T704) and indicated lipids compositions (the molar ratio of DLin-MC3-DMA:DSPC:Cholesterol:DMG-PEG2000 was fixed at 50:10:38.5:1.5, and the ratio between nitrogen residues in ionizable lipid and nucleic acid phosphate groups (N/P ratio) is 6). The PoLixNano was administered to mice via nebulization, the bioluminescence signal in the lungs of mice was detected 6 h post-dosing (0.1 mg Fluc-mRNA per mouse). (n=3 biologically independent samples). **b**, Quantitative evaluating the expression levels of eGFP-mRNA in PoLixNano or LNP transfected DC 2.4 cells via flow cytometry (n=3 biologically independent replicates). **c,** Cytotoxicity assessment of PoLixNano and LNP (all formulations were prepared under conditions showing the highest transgene expression) after 4 h incubation with indicated cell lines. Untreated cells were used as the blank. **d**, Representative expression of eGFP-mRNA expression in organoids after 36 h of incubation with PoLixNano or LNP, both pre-nebulization (top) and post-nebulization (bottom), and corresponding flow cytometry analysis with representative flow cytometry plots. Scale bars: 200 µm. **e,** Quantitative flow cytometry analysis of enhanced green fluorescent protein (eGFP) expression in organoids transfected by PoLixNano or LNP containing eGFP-mRNA, both pre- and post-nebulization (n=3 biologically independent replicates). Statistical significance was analyzed using one-way ANOVA with Dunnett's multiple comparisons test in **a** and one-way ANOVA with Tukey's post-hoc test in **b**. A two-tailed unpaired *t*-tests was applied in **e**.

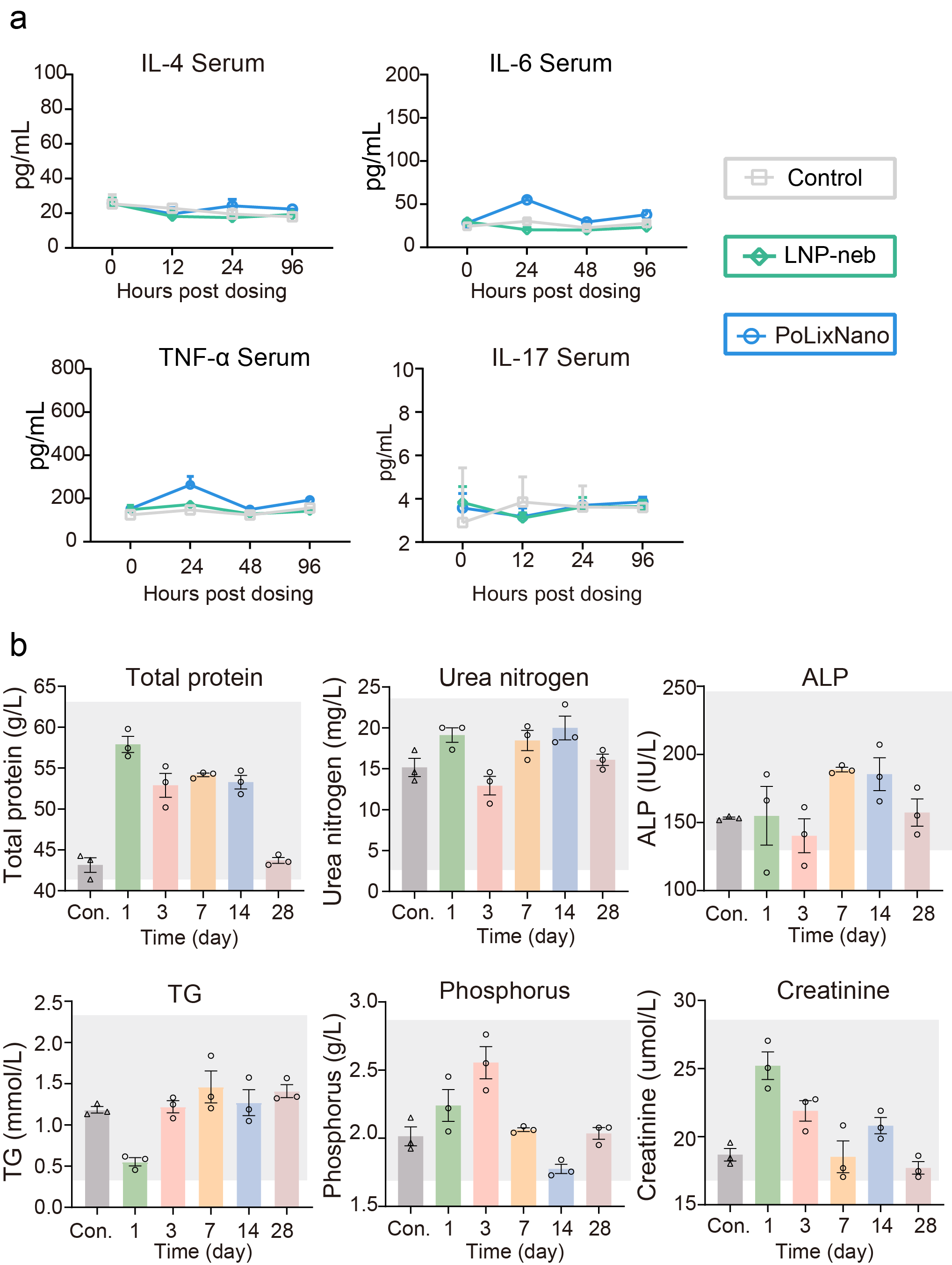

**Supplementary Fig. 2 | a,** Enzyme-linked immunoassay (ELISA) quantification of interleukin-4 (IL-4), interleukin-6 (IL-6), tumor necrosis factor-α (TNF-α) and interleukin-17 (IL-17) in serum samples of mice treated by nebulized PoLixNano or LNP (LNP-neb) at the indicated time points (n=3 biologically independent samples). **b,** Complete blood chemistry metrics of mice treated with nebulized PoLixNano (n=3 biologically independent samples). The grey regions represent the normal range of each parameter as a reference. TG: triglycerides; ALP: alkaline phosphatase. Untreated mice were used as the control (Con.).

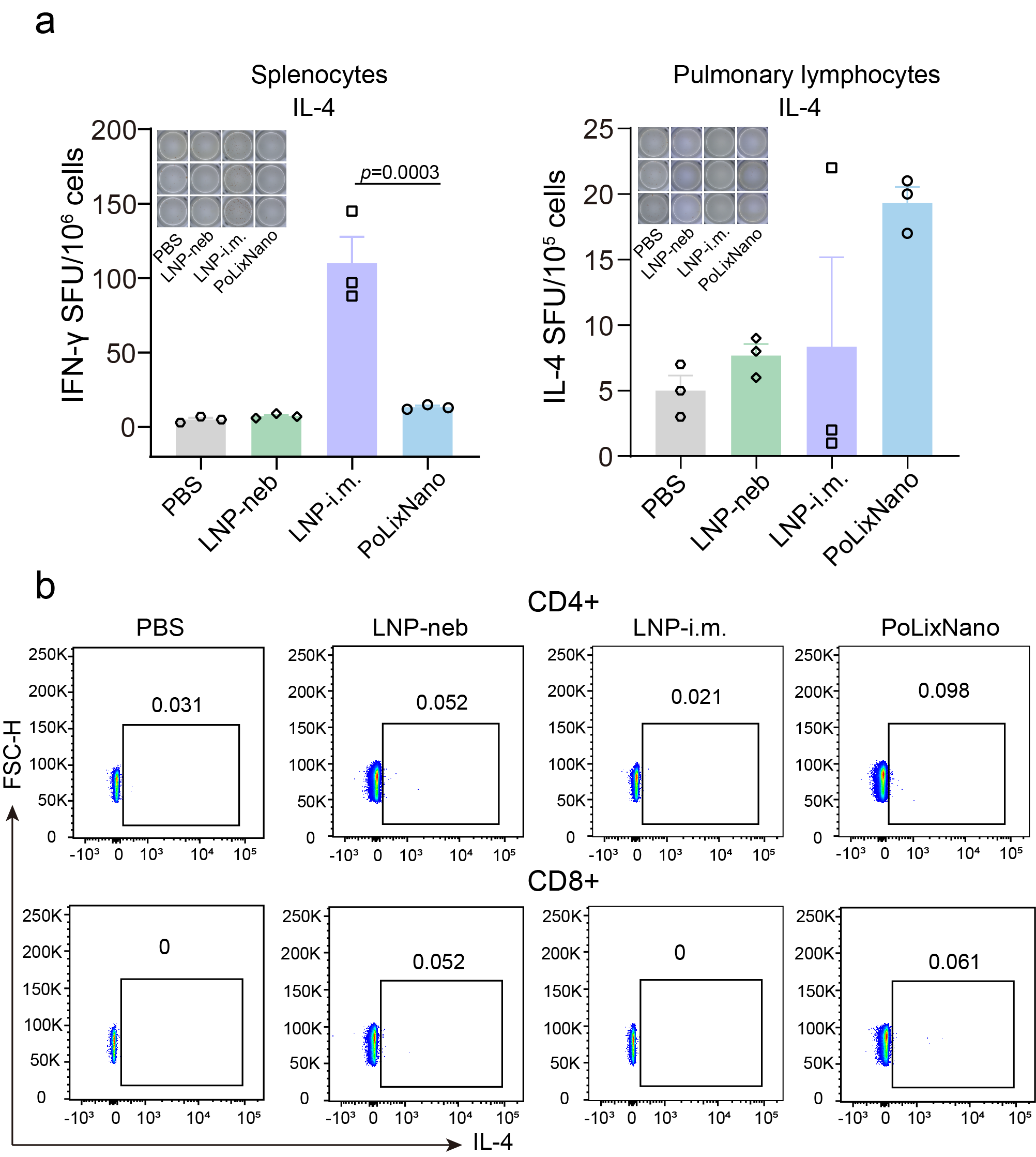

**Supplementary Fig. 3** **|** **a,** Enzyme-linked immunospot (ELISpot) assay of IL-4 spot-forming cells in splenocytes or pulmonary lymphocytes of indicated formulations vaccinated mice. The samples were collected on day 42 and re-stimulated with peptide pools of overlapping peptides spanning the SARS-CoV-2 receptor binding domain (RBD) region (n=3 biologically independent animals). **b**, Representative flow cytometry plots of CD4+IL-4+ and CD8+IL-4+ T cells among pulmonary lymphocytes on day 42 post-prime (n=3 biologically independent animals). Significance was calculated using a one-way ANOVA with Tukey's post-hoc test in **a**.

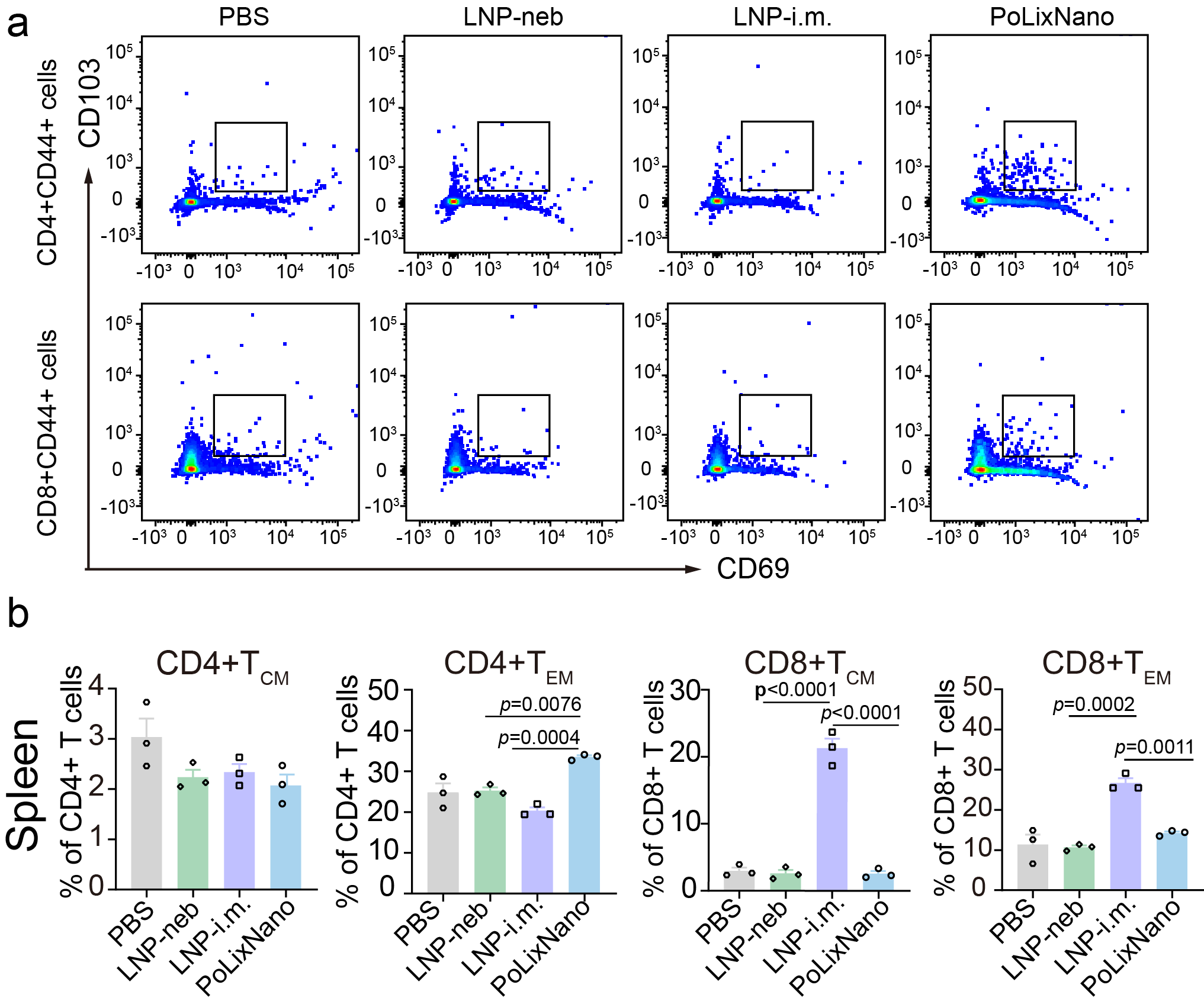

**Supplementary Fig. 4 | a**, Representative flow cytometry plots of CD4+ or CD8+ lung tissue resident memory T cells (T_RM_, CD69+CD103+) in the pulmonary lymphocytes of mice treated by indicated formulations. **b**, Flow cytometry analysis of CD4+ and CD8+ effector memory T cells (T_EM_) expressing CD44+CD62L- or central memory T cells (T_CM_) co-expressing CD44+CD62L+ in the splenocytes of mice vaccinated by indicated formulations. Significance was calculated using a one-way ANOVA with Tukey's post-hoc test.

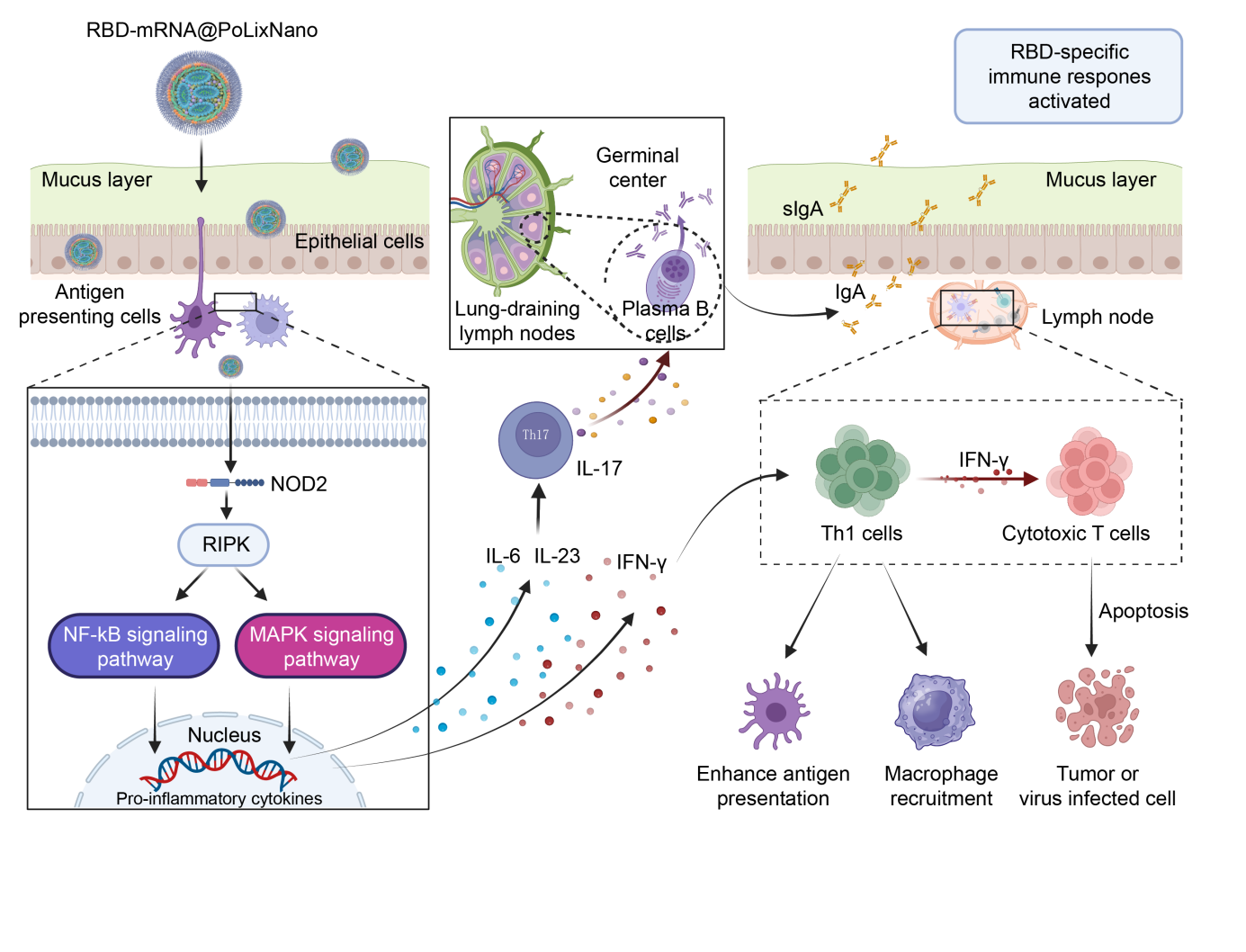

**Supplementary Fig. 5 |** Schematic diagram of the speculated mechanism of the mucosal adjuvant effects that are exerted by PoLixNano to enhance the immune responses.

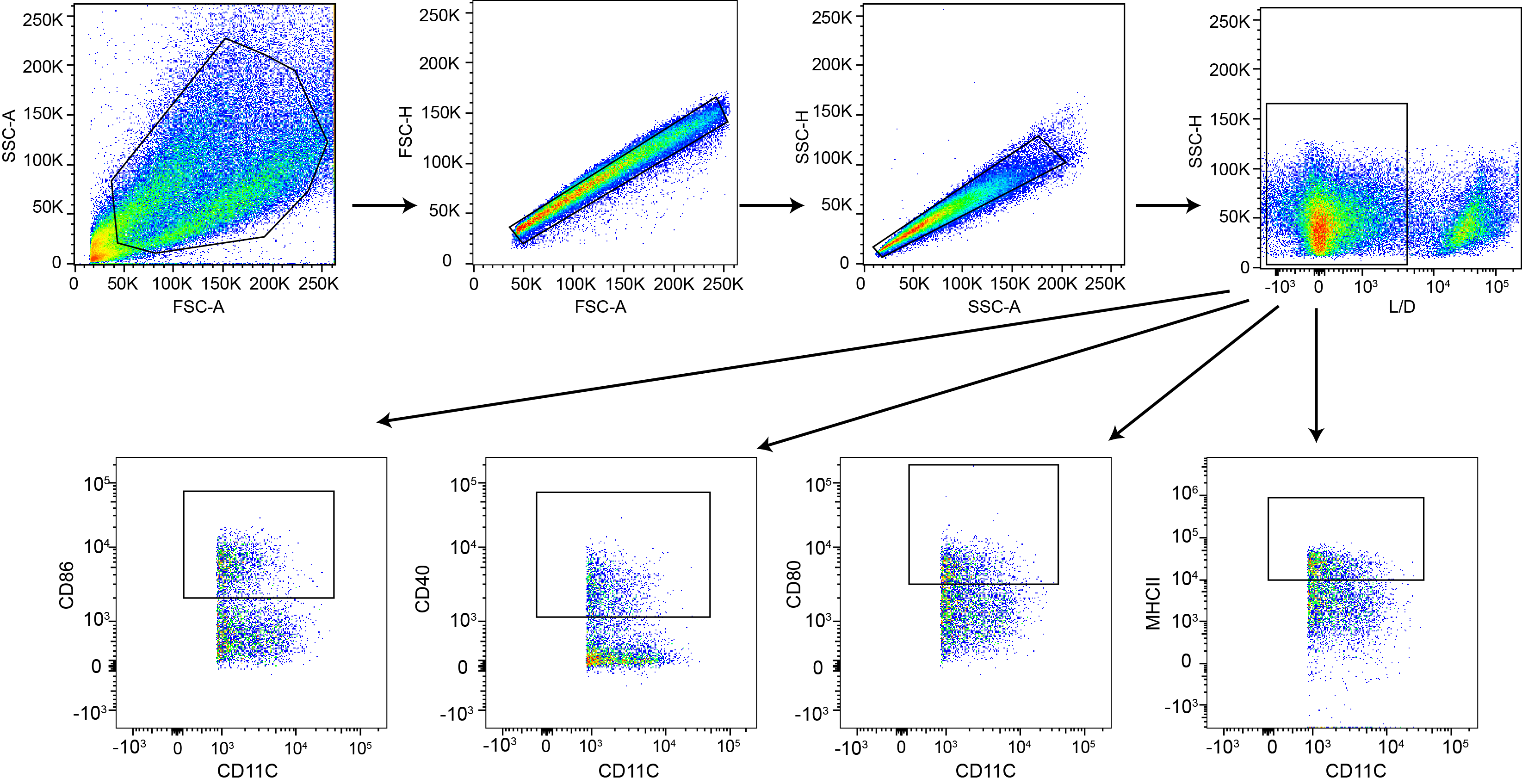

**Supplementary Fig. 6 |** Flow cytometry gating strategy for analyzing bone-marrow-derived dendritic cells (BMDCs), investigating the cells of expressing activation markers, i.e. CD86, CD40, CD80, or MHCII.

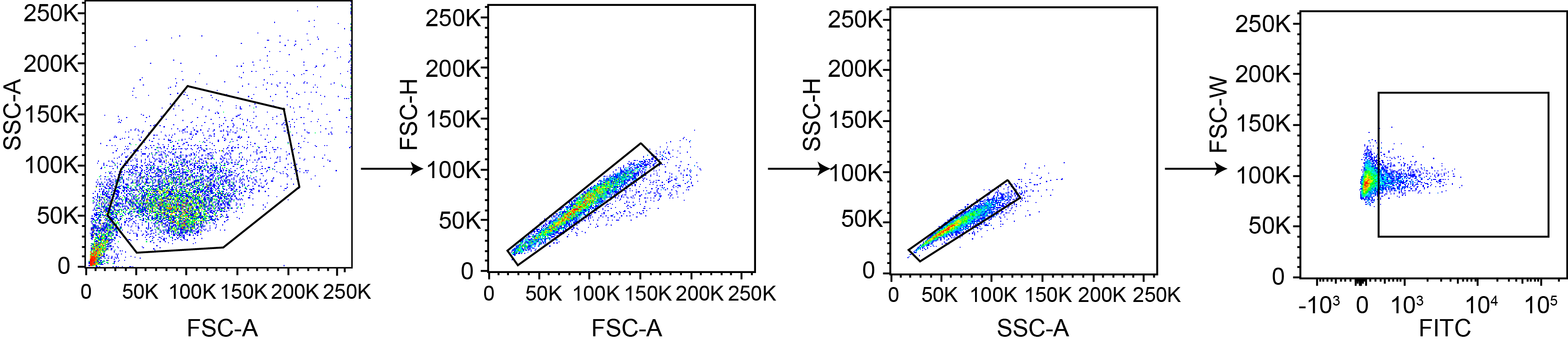

**Supplementary Fig. 7 |** Flow cytometry gating strategy for evaluating the expression of eGFP in single cells generated from organoids.

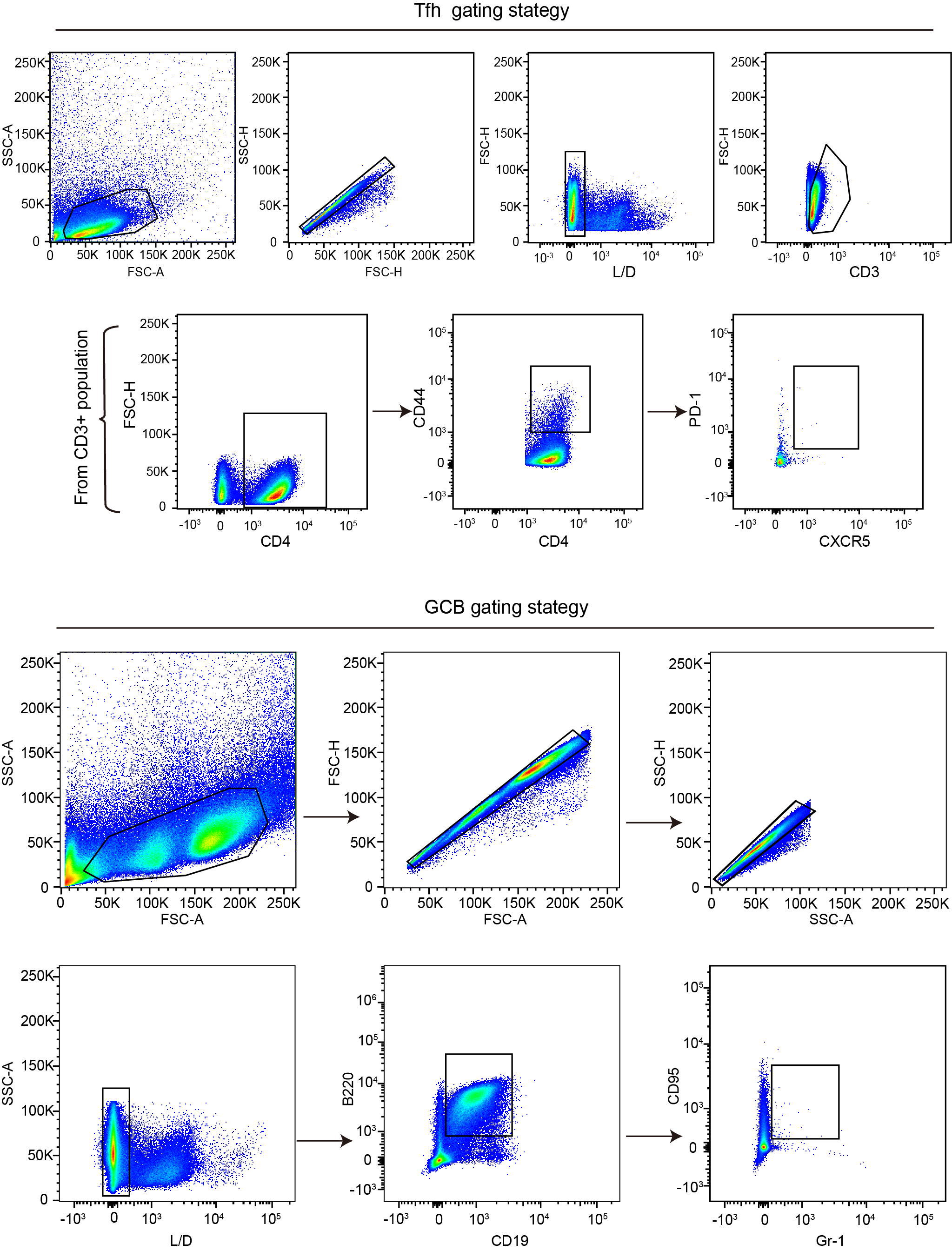

**Supplementary Fig. 8 |** Representative flow cytometry gating strategy for identifying T follicular helper cells (Tfh) and germinal center B cells (GCB) cells in mediastinal lymph nodes (MLN).

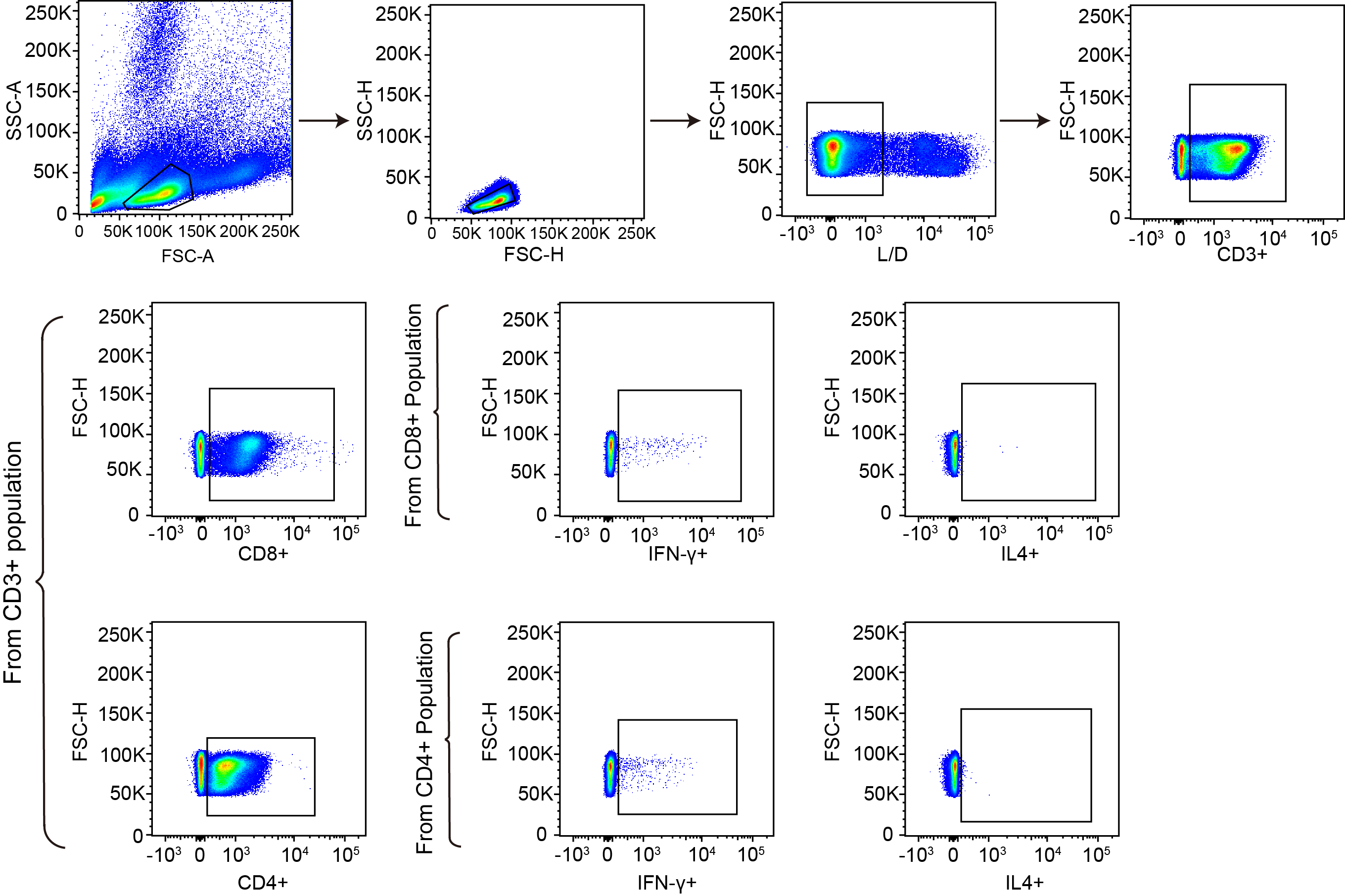

**Supplementary Fig. 9 |** Gating strategy applied for the flow cytometry of IFN-γ and IL-4 secreting CD4+ T cells and CD8+ T cells upon SARS-CoV-2 RBD peptide pools re-stimulation.

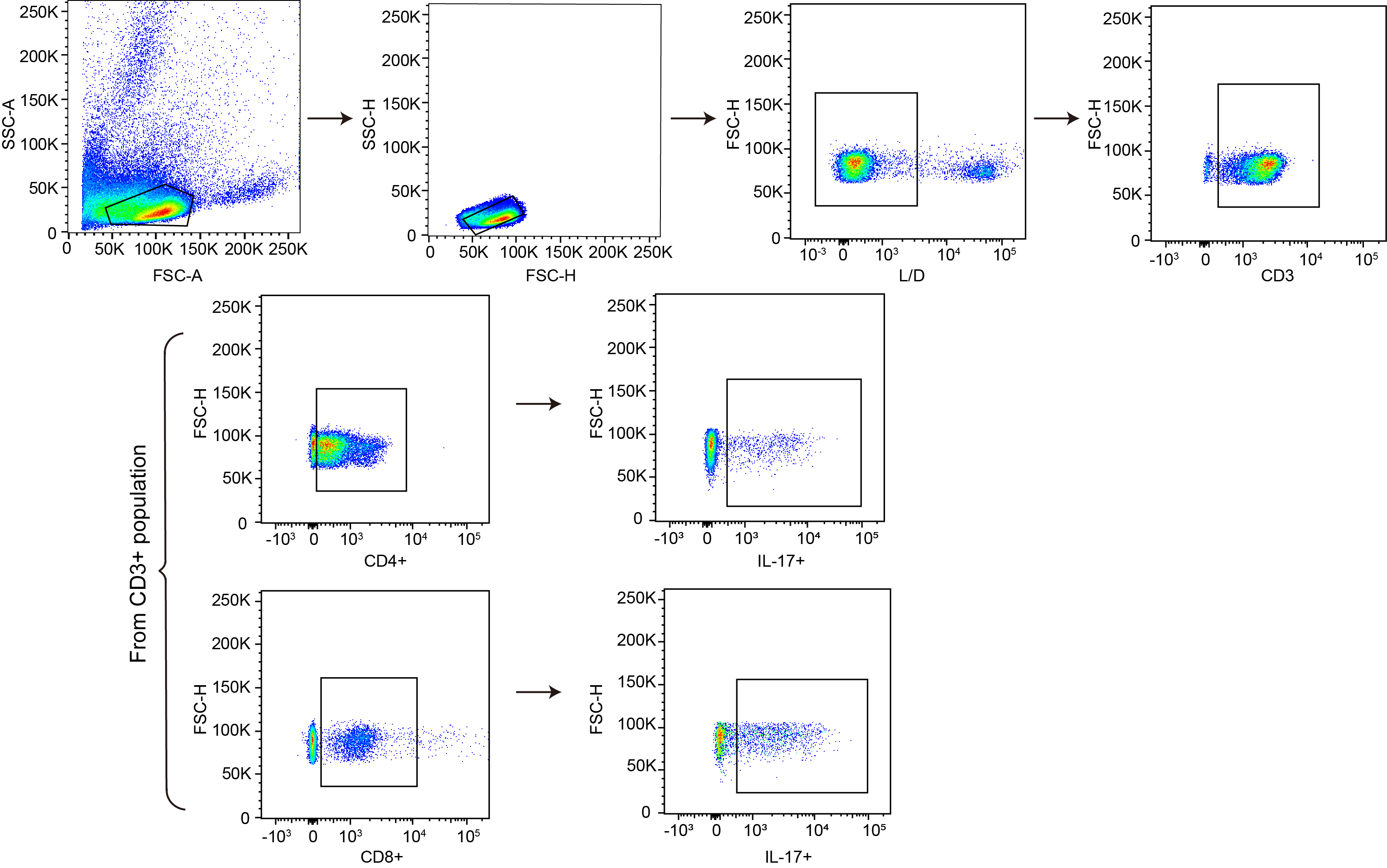

**Supplementary Fig. 10 |** Gating strategy applied for the flow cytometry of IL-17 secreting CD4+ T cells and CD8+ T cells upon SARS-CoV-2 RBD peptide pools re-stimulation.

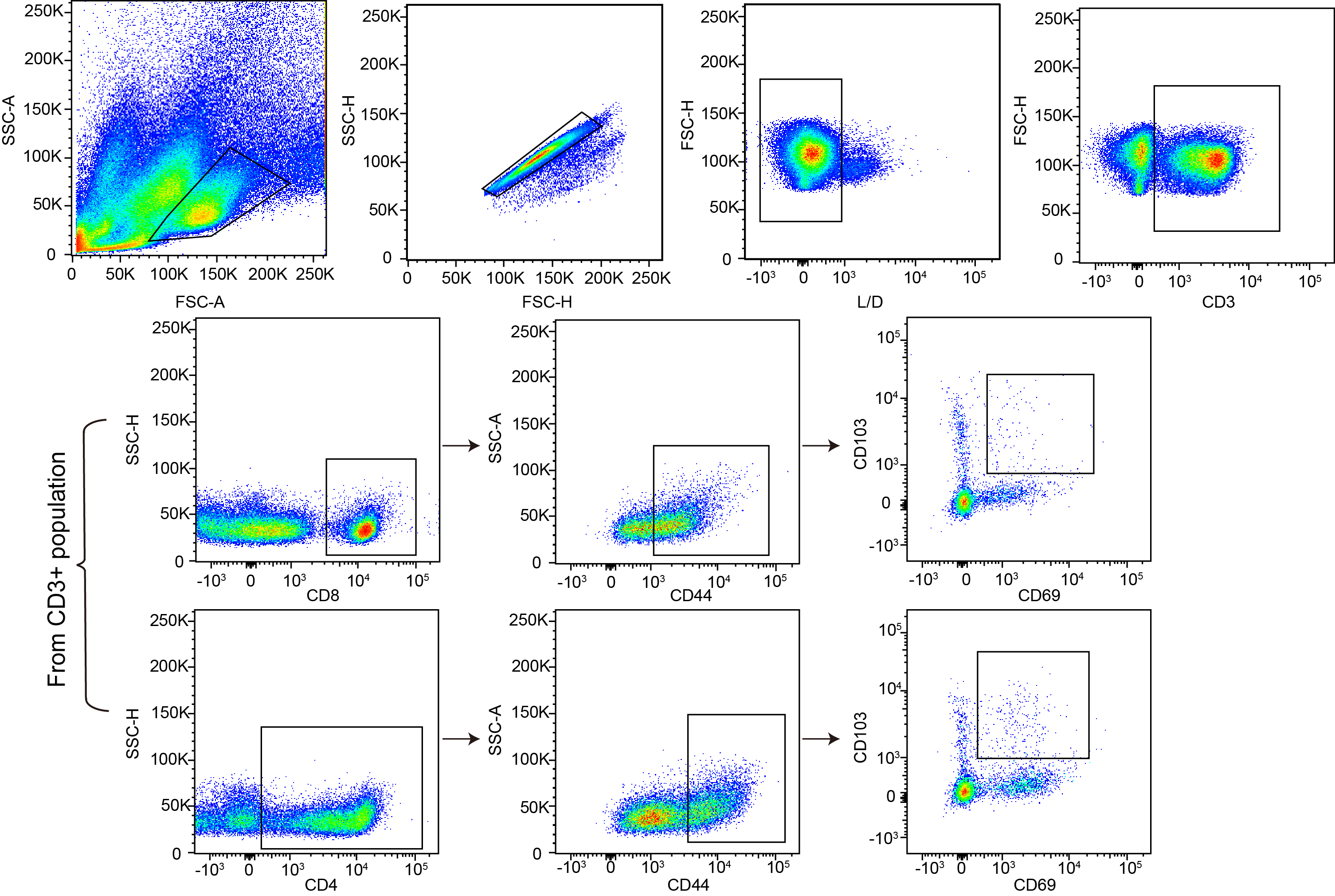

**Supplementary Fig. 11 |** Gating strategy applied for the flow cytometry analysis of pulmonary tissue-resident memory (T_RM_) CD8 + or CD4+ T cells.

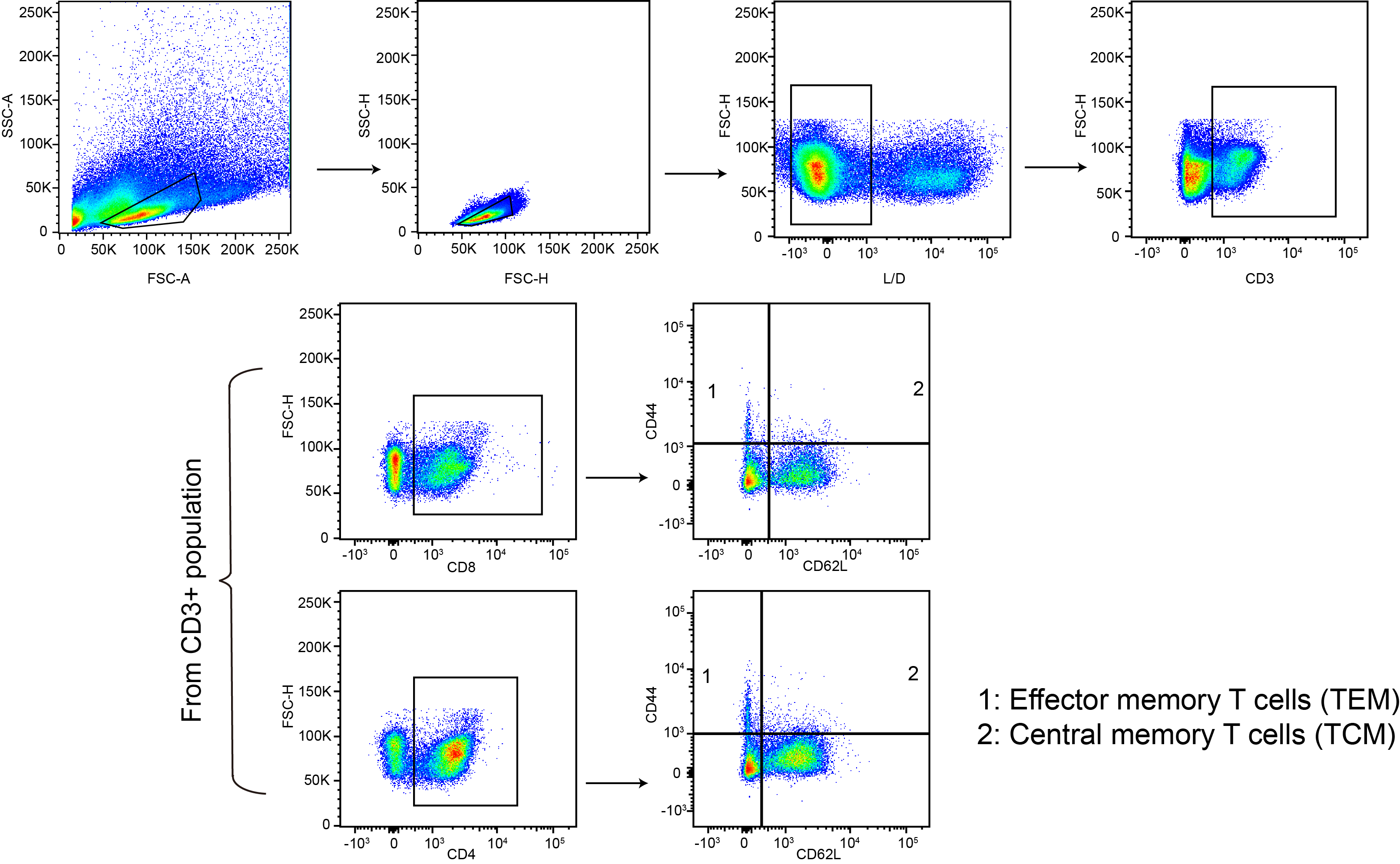

**Supplementary Fig. 12 |** Flow cytometry gating strategy for identifying pulmonary CD8+ or CD4+ effector memory (T_EM_) and central memory (T_CM_) T cells.

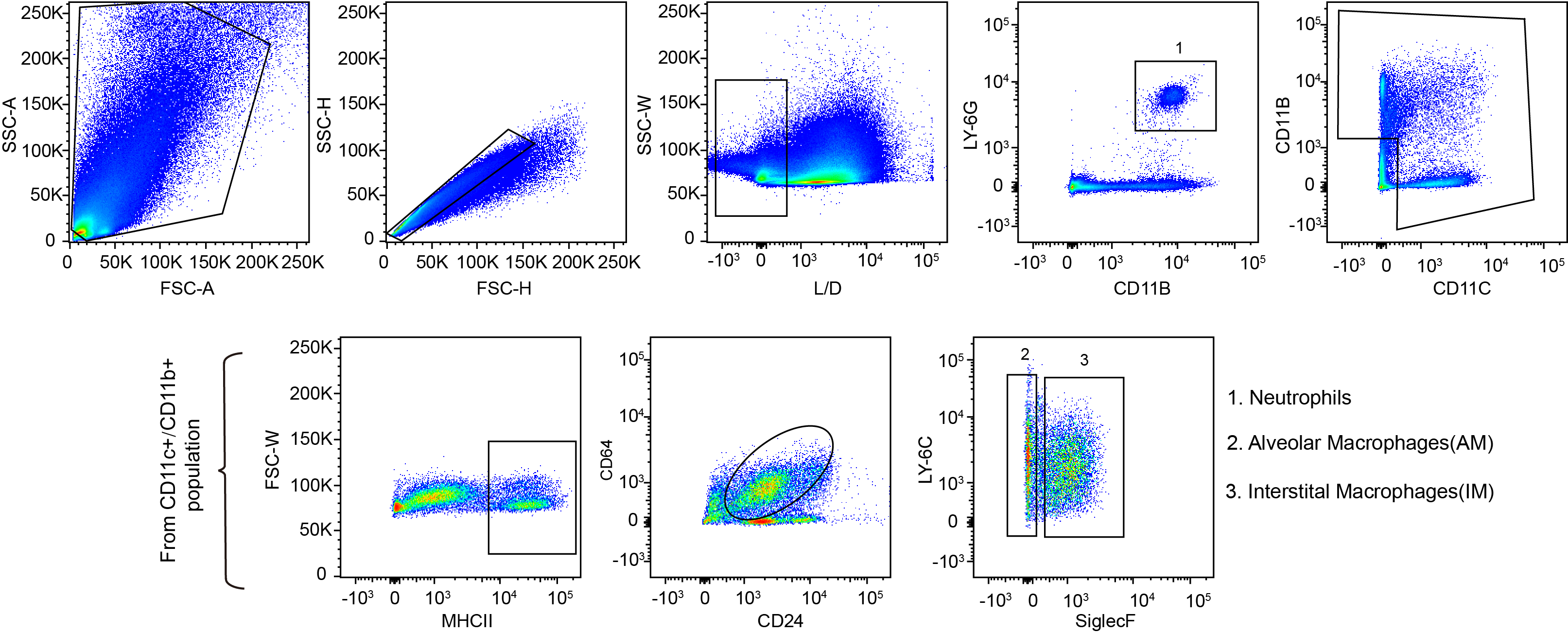

**Supplementary Fig. 13 |** Flow cytometry gating strategy used to distinguish neutrophils, alveolar macrophages (AMs), and interstitial macrophages (IMs) from pulmonary myeloid cell populations.
